# IRSp53 shapes the plasma membrane and controls polarized transport at the nascent lumen during epithelial morphogenesis

**DOI:** 10.1101/856369

**Authors:** Sara Bisi, Syed Abrar Rizvi, Stefano Marchesi, Davide Carra, Galina V. Beznoussenko, Ines Ferrara, Gianluca Deflorian, Alexander Mironov, Giovanni Bertalot, Federica Pisati, Amanda Oldani, Angela Cattaneo, Salvatore Pece, Giuseppe Viale, Angela Bachi, Claudio Tripodo, Giorgio Scita, Andrea Disanza

## Abstract

Establishment of apical–basal cell polarity is necessary for generation of luminal and tubular structures during epithelial morphogenesis. Molecules acting at the membrane/ actin interface are expected to be crucial in governing these processes. Here, we show that the I-BAR-containing IRSp53 protein is restricted to the luminal side of epithelial cells of various glandular organs, and is specifically enriched in renal tubules in human, mice, and zebrafish. Using three-dimensional cultures of renal MDCK and intestinal Caco-2 cysts, we show that IRSp53 is recruited early after the first cell division along the forming apical lumen, and is essential for formation of a single lumen and for positioning of the polarity determinants aPKC and podocalyxin. Molecularly, IRSp53 directly binds to and controls localization of the inactive form of the small GTPase RAB35, a tethering factor for apical determinants. The interaction of IRSp53 with the actin capping protein EPS8 is critical for restricting IRSp53 localization. Correlative light and electron microscopy shows that IRSp53 loss perturbs the shape and continuity of the opposing apical membrane during the initial phase of lumenogenesis, which leads to preservation of multiple cytoplasmic bridges that interrupt the continuity of the nascent lumen. At the organism level, genetic removal of IRSp53 results in abnormal renal tubulogenesis, with defects in tubular polarity and architectural organization in both IRSp53 zebrafish mutant lines and IRSp53-KO murine models. Thus, IRSp53 acts as a platform for spatiotemporal regulation of assembly of the multi-protein complexes that shape the luminal membrane during the early steps of epithelial lumen morphogenesis.

## Introduction

Many internal epithelial organs consist of a polarized cell monolayer that surrounds a central apical lumen. Polarization requires interactions between the signaling complexes and scaffolds that define cortical domains with membrane-sorting machinery^1^. This architectural organization and the morphogenesis processes that underlie it can be reproduced by plating cells on pliable matrigel in a matrigel-containing medium that provides the required mechanochemical cues for formation of a polarized hollow sphere (cyst). The *de-novo* apical–basal polarity in cysts arises from successive divisions of a single, nonpolarized cyst-forming cell^2, 3^. The first symmetry-breaking event occurs already during late telophase, at the first cell division, when the midbody is formed after cytokinetic furrow ingression^4, 5^. Around the midbody, the apical membrane initiation site (AMIS) is assembled, which establishes the location of the nascent lumen. Assembly of the AMIS at the midbody is mediated by both microtubules and branched actin filaments that are generated via the RAC1-WAVE axis, which promotes recruitment and anchoring of vesicles via the adhesion protein Cingulin^6^. In addition, transmembrane proteins, such as podocalyxin (PODXL; classical apical marker, also known as GP135) and Crumbs3, are transcytosed from the plasma membrane facing the extracellular matrix (ECM) towards the first cell–cell contact site^7–9^. This occurs via RAB11-RAB8 endo/exosomes trafficking and through a direct anchoring with with RAB35 at the AMIS^7, 10^. Thus, coordination between actin cytoskeletal dynamics and membrane trafficking is essential for the initiation of a polarized central lumen.

Despite this wealth of knowledge, the molecular players that link the spatially restricted actin dynamics and the polarized delivery of endosomal vesicles remain to be completely understood. What determines the shape of the opposing apical membrane during the initial phase of lumenogenesis is also not clear. Proteins that can sense and shape the curvature of the plasma membrane at the AMIS and physically link it with the underlying cytoskeleton will be critical here. We postulated that IRSp53 can fulfill this function.

IRSp53 regulates the dynamic interplay between the plasma membrane and the actin cytoskeleton during directional migration and invasion of cells^11–15^. Accordingly, IRSp53 localizes at the tips of both filopodia and lamellipodia^16^. At these sites, IRSp53 senses and promotes membrane curvature through its I-BAR domain, and acts as an effector of either CDC42 or RAC1 GTPases by ensuring the localized and temporally-regulated recruitment of a number of actin regulatory proteins^11–13, 17–19^. These proteins include the nucleator-promoting WAVE complex, essential for branched polymerization of actin during lamellipodia extension and the linear actin elongators VASP and mDia1, which together with EPS8 (an actin capping and cross-linking protein^20^) initiate and promote the extension of filopodia protrusions^12, 21^. For the formation of the these structures, through its I-BAR domain, IRSp53 undergoes a phase separation that facilitates protein clustering^22^ and the recruitment of Ezrin, a member of the ezrin-radixin-moesin (ERM) protein family, that links the actin cortex to cortex to the cell membranes^23^. The subsequent binding of activated CDC42 to this cluster leads to inhibition of the the weak capping activity of IRSp53. It further promotes a structural change in IRSp53 that facilitates the interaction of IRSp53, via its now-liberated SH3 domain, with actin linear elongators and cross-linkers to support the growth of filopodia^12, 24^. In addition, IRSp53 has been reported to have a role in the assembly of cell–cell and cell–ECM adhesions downstream of ‘polarity-regulating kinase partitioning-defective 1b’ (PAR1B), and to be involved in the polarized architectural organization of Madin Darby canine kidney (MDCK) epithelial cells *in vitro*^26, 27^. The molecular pathways and interactors of IRSp53 in the control of these processes and their physiological relevance at the organism level, however, remains poorly understood.

Here, we show that IRSp53 is apically restricted at the luminal side of various epithelial tubular and glandular human, murine, and zebrafish tissues. Further, IRSp53 is recruited early after the first cell division at the AMIS, where it ensures the continuity and ultra-structural architecture of the opposing plasma membrane, and formation of a single apical domain and localized recruitment of aPKC and PODXL. Molecularly, through a positively charged patch on its I-BAR-domain, IRSp53 functions by binding directly to the inactive form of RAB35, which colocalizes with IRSp53 at the onset of AMIS, while also controlling PODXL trafficking. Additionally, interactions with the plasma membrane and EPS8 are required for correct localization of IRSp53, and thus for its activity. The critical physiological role of IRSp53 in epithelial morphogenesis is supported by the finding that genetic deletion of IRSp53 results in abnormal renal tubulogenesis, with defects in tubular polarity, architectural organization, and lumen formation during kidney development and in the adult kidney in IRSp53 zebrafish mutant lines and IRSp53-KO murine models. We propose that IRSp53 is pivotal for the assembly of the RAB35–IRSp53–EPS8 macromolecular complex that is necessary for correct kidney morphogenesis and lumenogenesis, through its control of polarized transport and the shape of the apical plasma membrane.

## Results

### IRSp53 localizes to the apical lumen of different organs/ tissues and in three-dimensional cysts *in vitro*

To gain information regarding the physiological role of IRSp53 in epithelial morphogenesis, we examined its expression (Supplementary Fig. S1A) and localization across a variety of epithelial tissues characterized by glandular or tubular structures, in human, mice, and zebrafish by immunohistochemistry and immunofluorescence (Fig. 1A-D). In all of the tissues examined, IRSp53 localized to the luminal-facing side of the epithelial cells, such as in human renal convoluted tubules, colon mucosal crypts, gastric glands, prostate, and salivary and mammary gland ducts and acini (Fig. 1A, B). We observed a similar luminal distribution in the kidney, colon, stomach and mammary gland of adult mice (Fig. 1C), and in the pronephric ducts of developing zebrafish (Fig. 1D).

**Figure 1.**
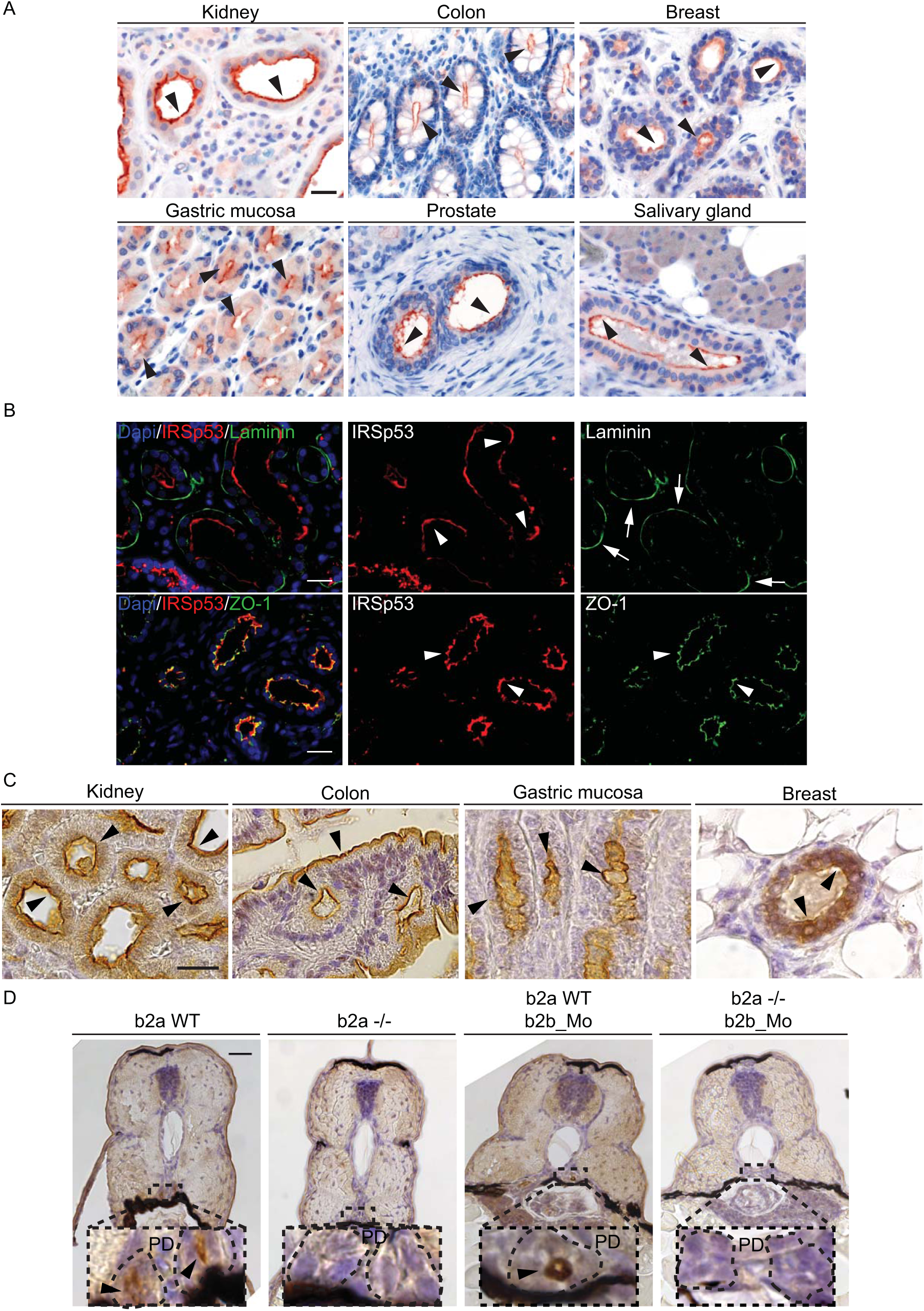
IRSp53 is apically restricted to the luminal side of various epithelial human, murine and zebrafish tissues. A) IHC analysis of IRSp53 expression and localization in the indicated human tissues and organs. Arrowheads indicate the apical/luminal enrichment of IRSp53. Scale bar, 50 µm. B) IF analysis of human kidney. Samples were stained with anti-IRSp53 (red), anti-Laminin (green) and DAPI (upper panels) or with anti-IRSp53 (red) anti-ZO-1 (green) and DAPI (lower panels). Arrowheads and arrows indicate the apical/luminal localization of IRSp53 and ZO-1, or the basal enrichment of Laminin, respectively. Scale bar, 50 µm. C) IHC analysis of the expression and localization of IRSp53 in the indicated murine tissues and organs. Arrowheads indicate the apical/luminal enrichment of IRSp53. Scale bar, 20 µm. D) IHC analysis of Baiap2a and Baiap2b expression and localization in zebrafish embryo (Inset: PD) at 72 hpf in the indicated genetic backgrounds: *baiap2a wild-type* (b2a WT), *baiap2a* mutant (b2a -/-), *baiap2b* morphant (b2a WT b2b_Mo) and *baiap2a* mutant *baiap2b* morphant (b2a -/- b2b_Mo). Arrowheads indicate the apical/luminal enrichment of IRSp53 in the pronephric duct. Scale bar, 50 µm (insets, 200 µm).

The apical localization of IRSp53 in luminal epithelial cells was further confirmed using well-established *in-vitro* two-dimensional (2D) monolayers and 3D cysts of MDCK and colorectal adenocarcinoma (Caco-2) epithelial cells. These cells form polarized monolayers when grown to confluency on 2D substrates. However, on cultivation on top of a matrigel cushion (for MDCK cells) and when embedded in a matrigel–collagen matrix (both MDCK, Caco-2 cells) they instead generated single spheroids with a lumen (cysts). This process is believed to reproduce the morphogenesis programs of the respective tissues of origin^1, 28^. Endogenous IRSp53, like ectopically expressed GFP-IRSp53, localized at the apical membranes and decorated the lumen of MDCK cysts with a continuous pattern (Fig. 2A and Supplementary Fig. S2A). There was also a similar apically restricted distribution in the Caco-2 cell hollow spheroids (Supplementary Fig. S2B-C). Collectively, these findings indicated that IRSp53 is apically restricted at the luminal side of these different epithelial tissues *in vitro* and *in vivo*.

**Figure 2.**
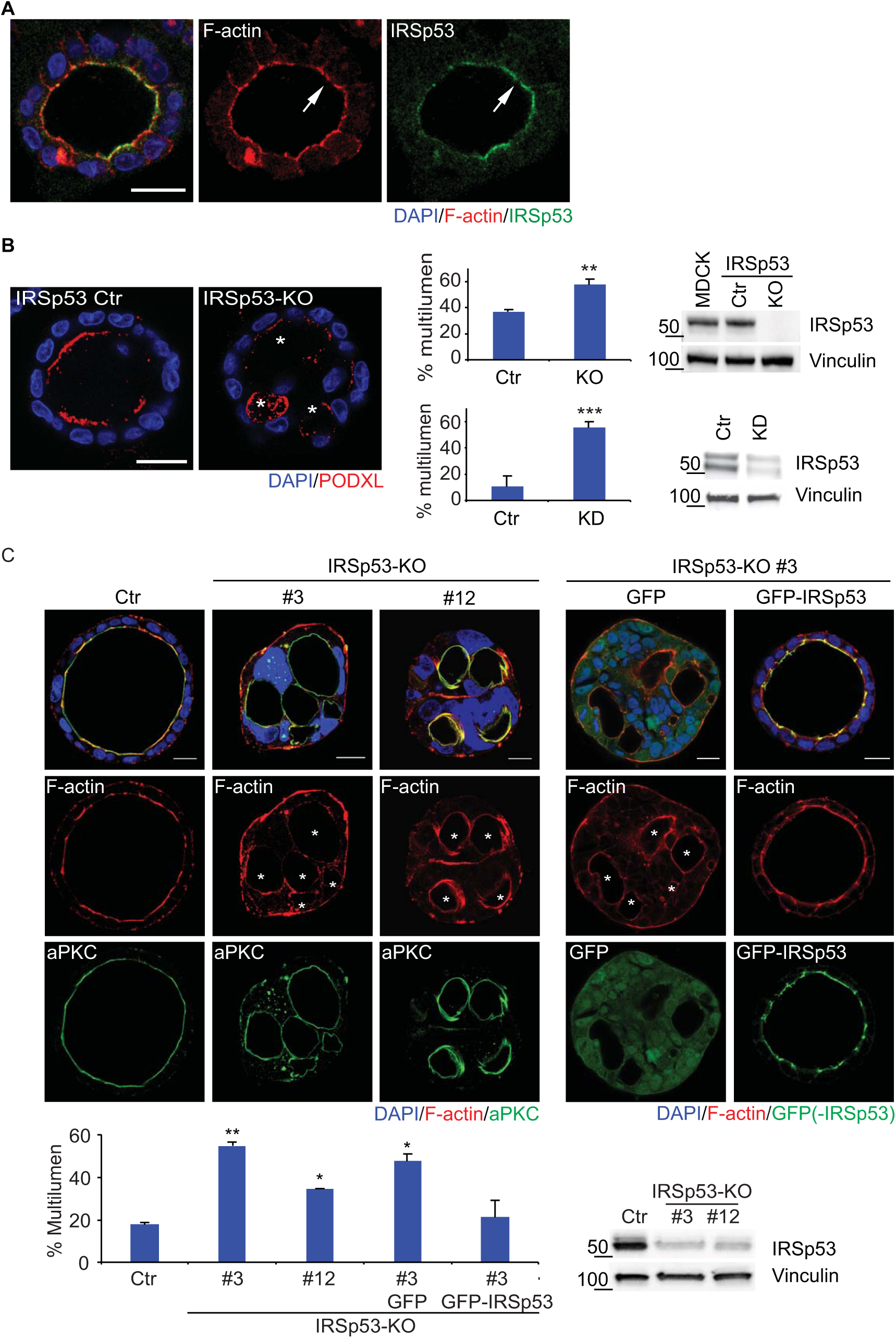
IRSp53 removal increased the formation of cysts with multiple lumens. A) IRSP53 decorates the apical lumen of MDCK epithelial cysts. MDCK were seeded as single cells on a Matrigel layer and left growing to form three-dimensional (3D) cysts. The cysts were fixed and stained an anti-IRSp53 antibody (green), and rhodamine phalloidin (red) to visualized F-actin and DAPI (blue). Arrows indicate IRSp53 and F-actin enrichment at the luminal side of the cyst. Scale bar, 18 µm. B) MDCK control cells (IRSp53 ctr) or IRSp53-KO obtained by CRIPR/Cas9 genome editing, or MDCK control or IRSp53-KD (not shown), were seeded as single cells on a Matrigel layer and left to grow for 5 days to form 3D cysts. *Left*: the cysts were fixed and stained with anti-podocalyxin (PODXL, red) and DAPI (blue). Asterisks, multiple lumens in IRSp53-KO cyst. Scale bar, 18 µm. *Central*: Quantification of cysts with multiple lumens. Data are expressed as means ± SD. At least 30 cysts/experiment were analysed in three independent experiments. **p < 0.01; ***p < 0.001. *Right*: Immunoblotting of expression levels of IRSp53 and vinculin. Molecular weight markers are indicated. C) *Top left*: Caco-2 control (Ctr) or IRSp53-KO clones #3 and #12, obtained by CRIPR/Cas9 genome editing, were embedded as single cells into Matrigel/ collagen matrix and left to grow to form 3D cysts (see Methods for details). The cysts were fixed and stained with an anti-atypical-PKC antibody (a-PKC, green), rhodamine-phalloidin to detect F-actin (red) and DAPI (blue). Asterisks, multi-lumens in IRSp53-KO cyst. Scale bar, 10 µm. *Top right*: Single cells from IRSp53-KO clone #3 stably-infected to express GFP or murine GFP-IRSp53 embedded into Matrigel/ collagen matrix and left to grow to form 3D cysts. The cysts were fixed and processed for epifluorescence to visualize GFP or GFP-IRSp53 and stained with rhodamine-phalloidin to detect F-actin (red) and DAPI (blue). Asterisks, multiple lumens in IRSp53-KO GFP cyst. Scale bar, 10 µm. *Lower left*: Quantification of cysts with multiple lumens. Data are expressed as mean ± SD. At least 30 cysts/experiment were analysed in two independent experiments. *p < 0.05; **p < 0.01. *Lower right:* Immunoblotting of expression levels of IRSp53 and vinculin (for GFP and GFP-IRSp53 expression levels, see Supplementary Fig. S5A).

### The loss of IRSp53 disrupts cyst morphogenesis and results in formation of multiple lumens

To determine whether IRSp53 is involved in the establishment of apical–basal polarity and lumen formation, IRSp53 was depleted in MDCK cells using CRISPR/CAS9, shRNA^26^, and siRNAs, and in Caco-2 cells using CRISPR/CAS9. Single MDCK-control and IRSp53-depleted cells were seeded onto Matrigel for 6 days. As expected, most of the control cysts (>75%) developed a single, actin-positive central lumen that was surrounded by an epithelial monolayer, which was decorated by apical proteins, such as PODXL (Fig. 2B). Conversely, <50% of the MDCK IRSp53-silenced cysts formed a single apical lumen, while the remaining showed either multiple or aberrant lumens (Fig. 2B). A significant increase in multi-luminal aberrant cysts was also seen in two independently generated Caco-2 cell CRISPR–IRSp53-KO clones, as compared to the control Caco-2 cells (Fig. 2C). Importantly, the aberrant multi-luminal phenotype was rescued upon re-expression of GFP-IRSp53 in both the IRSp53-silenced MDCK and Caco-2 *IRSp53*-KO cells (Fig. 2C, Fig. 5E and Supplementary Fig. S5A). Thus, IRSp53 has a crucial role in the morphogenesis of these epithelial organoids.

### IRSp53 controls the distribution and trafficking of the apical protein PODXL

To investigate how IRSp53 contributes to epithelial cell polarity and lumenogenesis programs, we examined the localization and dynamics of IRSp53 in the very early phases of cystogenesis (Fig. 3A)^7^. In the first 8 h to 12 h after plating of the control cells, PODXL localized to the peripheral surface of the cells, before being internalized into vesicles. Subsequently, PODXL was delivered to the opposing membrane between the two daughter cells, where the apical lumen was formed *de novo* (Fig. 3A, B). IRSp53 became prominently enriched along the newly formed, opposing intercellular membrane well before PODXL recruitment (Fig. 3B, Movie S1). At later stages, IRSp53 and PODXL were concentrated at the AMIS, and when the cyst was formed, IRSp53 and PODXL co-stained along the lumen (Fig. 3B). Importantly, IRSp53 and PODXL were also co-localized in vesicle-like structures in the early phases of cystogenesis (Fig. 3B), which were subsequently targeted to the AMIS and the nascent lumen. These findings suggested that IRSp53 and PODXL might move together to or from endosomal recycling compartments. They also raise the possibility that IRSp53 is involved in the trafficking or targeting of PODXL to the AMIS.

**Figure 3.**
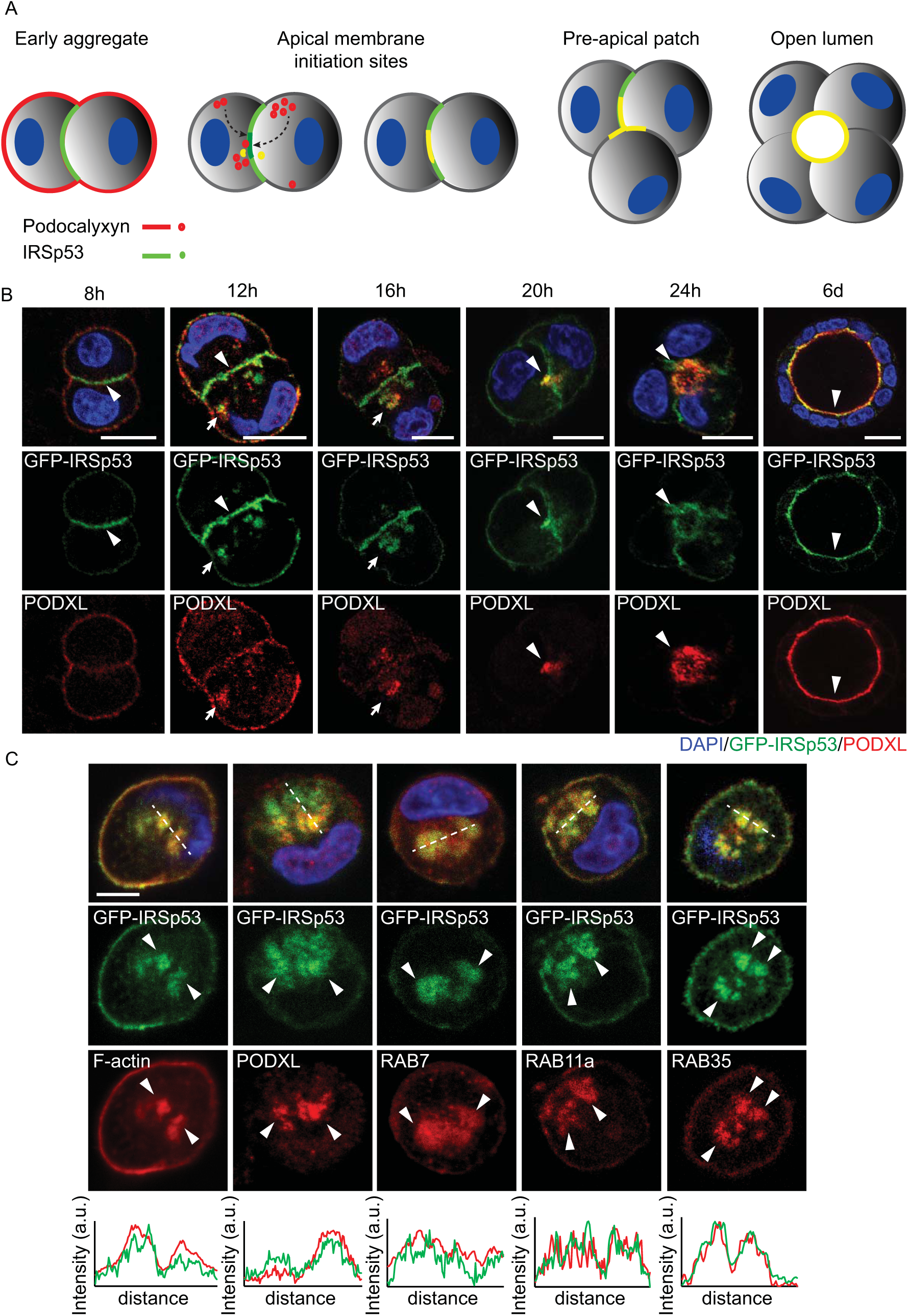
IRSP53 precedes PODXL localization to the apical membrane after the first cytokinesis during cyst development. A) Schematic representation of IRSp53 and PODXL trafficking during the early phases of lumenogenesis. B) MDCK cells expressing GFP-IRSp53 were seeded as single cells on a Matrigel layer and left to grow to form three-dimensional (3D) cysts. The cysts were fixed at the indicated time points, processed for epifluorescence to visualize GFP-IRSp53 (green) and stained with anti-PODXL (red) and DAPI (blue). Arrowheads, IRSp53 preceding PODXL relocalization at the AMIS at early time points; colocalization of IRSp53 and PODXL at the AMIS at later time points; and enrichment at the luminal side in the mature cyst. Arrows, IRSp53 and PODXL colocalization in vesicle-like structures. Scale bar, 18 µm. C) *Top*: MDCK cells, expressing GFP-IRSp53, were not transfected or were transfected with RFP-RAB7 or RFP-RAB35, and seeded as single cells on a Matrigel layer and fixed after 4 h of growth. The cysts were processed for epifluorescence to visualize GFP-IRSp53 (green) and RFP-RAB7 or RFP-RAB35, or stained with rhodamine-phalloidin (red), anti-PODXL (red) or anti-RAB11a (red), and DAPI (blue). Arrowheads, IRSp53 colocalization with the different trafficking markers. Scale bar, 10 µm. *Bottom*: Intensity profiles of the green and red channels over the vesicle-like structures (dashed lines in the merge channels), generated using the ImageJ software.

To distinguish among these possibilities, the cellular localization of IRSp53 was initially analyzed relative to a variety of proteins that can be used to label distinct trafficking compartments. MDCK cells that were stably expressing GFP-IRSp53 were used here, where this GFP-tagged IRSp53 showed a localization nearly identical to that of the endogenous protein (Fig. 2A, and Supplementary Fig. S2A). There was partial co-localization of GFP-IRSp53 with PODXL and actin in vesicle-like structures (Fig. 3C). These structures were also positive for RAB8, RAB7, RAB11a, and RAB35 (Fig. 3C), but not for the early and late endosome markers ‘early endosome antigen 1’ (EEA1) and ‘lysosome-associated membrane glycoprotein 1’ (LAMP1), respectively, or the Golgi marker Giantin (Supplementary Fig. S3A).

To determine whether IRSp53 is involved in the control of PODXL trafficking, we took advantage of the observation that PODXL is internalized from the apical membrane and undergoes trafficking to the so-called vacuolar apical compartment (VAC) upon calcium removal in MDCK polarized monolayers ^29^. First, we verified that PODXL is indeed internalized under these conditions. MDCK cell monolayers were incubated with a PODXL antibody that recognized the extracellular portion of PODXL *in vivo*, in the absence and presence of calcium. Detection of PODXL with a secondary fluorescence-conjugated anti-PODXL IgG after stripping of the cell surface with acid washes revealed that PODXL was internalized in the VACs (Supplementary Fig. S3B). More importantly, IRSp53 removal significantly delayed the relocalization of PODXL to the VACs, in comparison to the control cells (Supplementary Fig. S3C-E).

If IRSp53 is required for correct trafficking or anchoring of PODXL, its ablation should lead to mislocalization of PODXL during the early phases of cystogenesis. Consistent with this, while in control cells PODXL was enriched in a single, centrally located luminal spot at the four-cell stage, in the *IRSp53*-KO and knocked-down cells, two or more PODXL-positive foci were observed (Fig. 4A, B). Of note, the majority of *IRSp53*-silenced, early phase, multi-luminal cysts retained an apically restricted distribution of PODXL. However, a significant, albeit small, proportion of the cysts showed an inverted apical–basal polarity that was characterized by mislocalization of PODXL to the outer, ECM-facing, plasma membrane (Fig. 4A, B). We performed a similar experiment also in *IRSp53*-KO Caco-2 cells by monitoring the distribution of aPKC, which is the most commonly-used apical marker in this in for the distribution of aPKC^28, 30, 31^. Likewise in MDCK cells, IRSp53 depletion resulted in formation of aPKC multi-foci at the 3-4-cell stage during cystogenesis, as compared to the control Caco-2 cells (Supplementary Fig. S4A).

**Figure 4.**
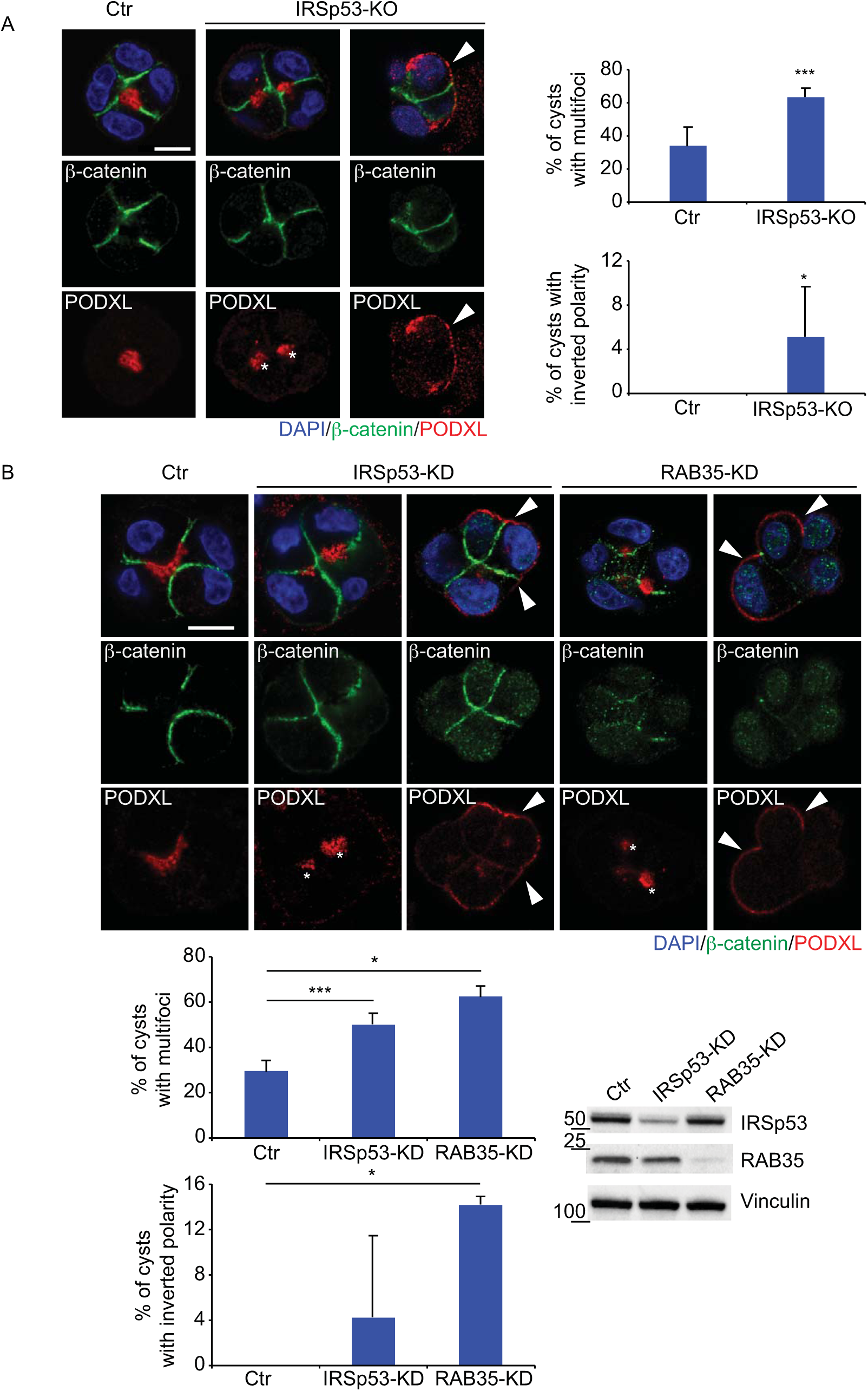
Loss of IRSp53 alters PODXL trafficking at the AMIS and apical-basal polarity. A) *Left*: MDCK cell control (Ctr) and *IRSp53*-KO were seeded as single cells on a matrigel layer and left to grow for 24/36 h. The cysts were fixed and stained with anti-β-catenin (green) and anti-PODXL (red) antibodies, and DAPI (blue). Asterisks, PODXL multi-foci in *IRSp53*-KO cyst. Arrowheads, PODXL staining at the basal membrane in *IRSp53*-KO cyst. Scale bar, 10 µm. *Right*: Quantification of multi-foci cysts (top) and inverted polarity cysts (bottom). Data are means ±SD. Four-cell stage cysts were analysed, as at least 20 cysts/experiment in five independent experiments. *p < 0.05; ***p < 0.001. B) *Top*: MDCK cell control (Ctr), *IRSp53*-KD or *RAB35*-KD cells were seeded as single cells on a Matrigel layer and left to grow for 24/36 h. The cysts were fixed and stained with anti-β-catenin (green) and anti-PODXL (red) antibodies, and DAPI (blue). Asterisks, PODXL multi-foci in *IRSp53*-KD and *RAB35*-KD cysts. Arrowheads, PODXL staining at the basal membrane in *IRSp53*-KD and *RAB35*-KD cysts. Scale bar, 10 µm. *Bottom left*: Quantification of multi-foci cysts (top) and inverted polarity cysts (bottom). Data are means ±SD. Four-cell stage cysts were analysed, as at least 20 cysts/experiment in five (multi-foci) or two (multi-foci, inverted polarity) independent experiments. *p < 0.05; ***p < 0.001. *Right bottom*: Immunoblotting of expression levels of IRSp53, RAB35 and vinculin to assess their down-regulation.

Collectively, these findings indicated that IRSp53 is implicated in the trafficking and/or correct targeting of apical proteins, and specifically of PODXL, to the *de-novo* forming lumen.

### IRSp53 is at the core of the RAB35 and EPS8 pathways in the control of lumen formation

The RAB GTPases are key regulators of PODXLtranscytosis^32^. Among these, we focused our attention on RAB35. RAB35 is involved in clathrin-mediated endocytosis^33^ and was recently shown to localize early at the AMIS, where it serves as a physical anchor for the targeting of PODXL in MDCK cell cystogenesis^10, 32^. During the initial phases of cyst development, IRSp53 and RAB35 were enriched along the opposing cell–cell membrane prior to the arrival of PODXL (Fig. 3B and Supplementary Fig. S4B). At the later stages, IRSp53, RAB35, and PODXL were concentrated at the AMIS (Fig. 3B and Supplementary Fig. S4B). IRSp53, RAB35, and PODXL also colocalized in intracellular vesicle-like structures (Fig. 3B, C, Supplementary Fig. S4B). More relevantly, silencing of RAB35, as previously reported^10, 32^ and similar to IRSp53 depletion, resulted in the formation of multi-lumen cysts, and in some cases, led to complete inversion of the apical–basal polarity (Fig. 4B). Of note, IRSp53 retained its localization to the AMIS in RAB35-silenced cells that showed inverted PODXL polarity (Supplementary Fig. S4C).

To determine whether IRSp53 and RAB35 interact physically, co-immunoprecipitation experiments were performed. Here, ectopically expressed IRSp53 interacted with RAB35 only upon serum starvation, which suggested that IRSp53 might preferentially associate with the inactive GDP-bound form of RAB35 (Fig. 5A). This possibility was verified using the recombinant purified dominant-negative RAB35S22N and constitutively active RAB35Q67L mutants of RAB35 in *in-vitro* overlay assays. Here, IRSp53 bound directly to RAB35 and showed higher apparent affinity for inactive RAB35S22N compared to the RAB35Q67L form (Supplementary Fig. S4D). Mapping of the interaction surfaces using various IRSp53 fragments or single-point mutants showed that the I-BAR domain of IRSp53 is necessary and sufficient for this association with RAB35S22N (Fig. 5B). We further restricted the binding interaction to a basic stretch of amino acids on the I-BAR domain. Accordingly, an IRSp53 mutant that lacked these critical positively charged residues (the I-BAR-4K domain mutant; IRSp53 I-BAR*: K142E, K143E, K146E, K147E^34^) did not interact with RAB35S22N under *in-vitro* binding or co-immunoprecipitation conditions (Fig. 5C, D).

**Figure 5.**
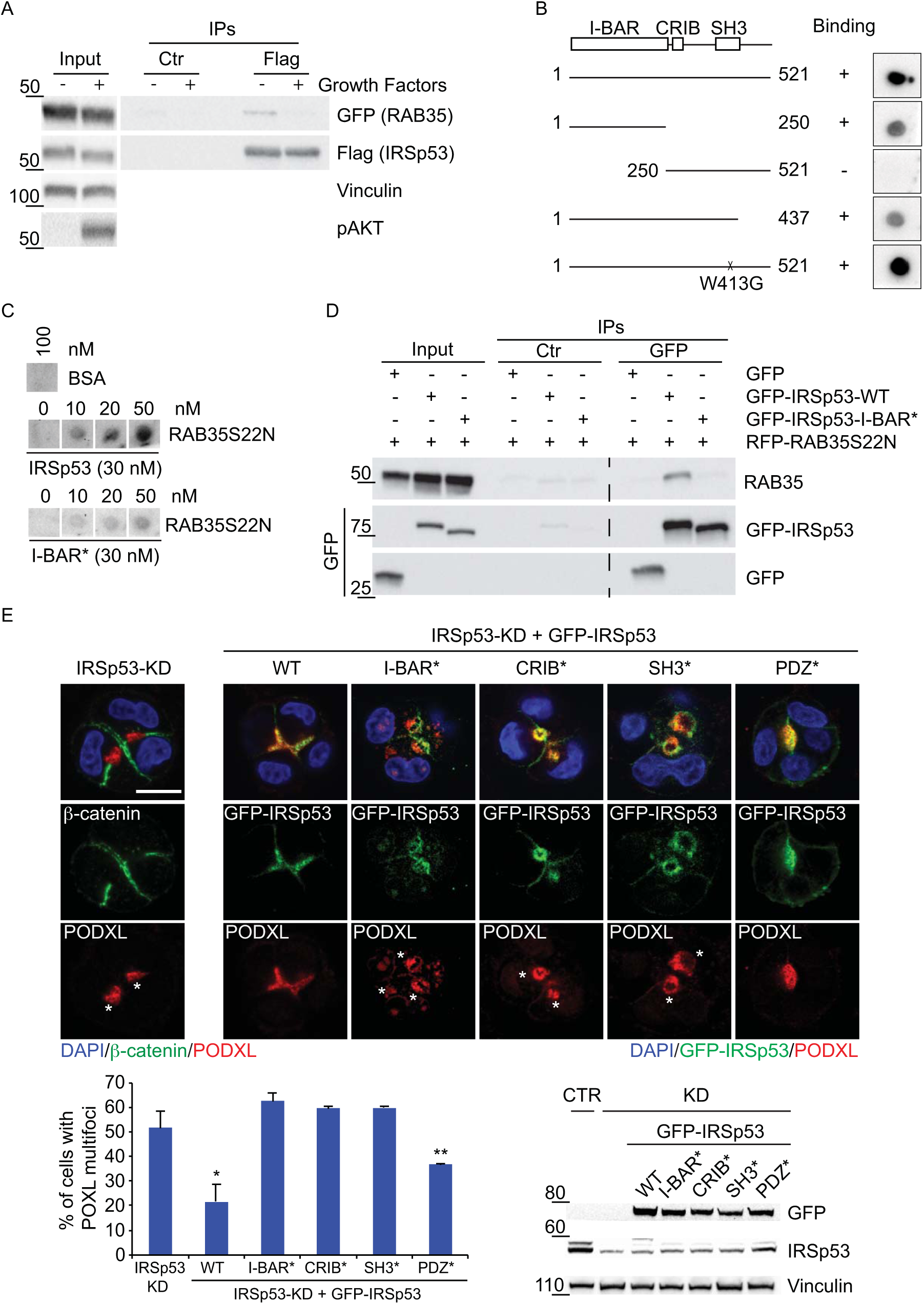
IRSp53 directly binds RAB35 in its inactive GDP-bound state. A) HeLa cell lysates (1 mg) from cells transfected with GFP-RAB35 and Flag-IRSp53, serum starved overnight, and without (-) or with (+) treatments with growth factors, were subjected to immunoprecipitation with anti-Flag (Flag) or control (Ctr) antibodies. Inputs (20 µg) and IPs were analyzed by immunoblotting with the indicated antibodies. pAKT was used for the positive control for growth factor stimulation. B) Structure function analysis. Equal amounts (50 nM) of recombinant purified IRSp53 full-length WT or the indicated fragments and mutant were spotted onto nitrocellulose and incubated with recombinant purified GST-RAB35S22N (50 nM). After washing, nitrocellulose membranes were immunoblotted with an anti-GST antibody. C) Indicated amounts of recombinant GST-RAB35S22N were spotted onto nitrocellulose and incubated with equal amounts (30 nM) of either recombinant purified IRSp53 WT or IRSp53 I-BAR* (K142E, K143E, K146E, K147E). After washing, nitrocellulose membranes were immunoblotted with an anti-IRSp53 antibody. BSA (100 nM) was used as the negative control. D) Cell lysates (2 mg) from MDCK cells transfected with GFP empty vector, GFP-IRSp53 wild-type (WT), GFP-IRSp53 I-BAR*, and RFP-RAB35S22N were subjected to immunoprecipitation with anti-GFP (GFP) or control (Ctr) antibodies. Inputs and IPs were immunoblotted with the indicated antibodies. E) *Top*: MDCK *IRSp53*-KD or *IRSp53*-KD cells were infected to express murine GFP-IRSp53 wild-typ (WT), I-BAR*, CRIB*, SH3* and PDZ*, and were seeded as single cells on a Matrigel layer and left to grow for 24/36 h. The cysts were fixed and stained with anti-β-catenin (green) or anti-PODXL (red) antibodies, and DAPI (blue), or processed for epifluorescence to visualize GFP-IRSp53 (green), and stained withan anti-PODXL antibody (red) and DAPI (blue). Asterisks, PODXL multi-foci. Scale bar, 10 µm. *Bottom left*: Quantification of multi-foci cysts. Data are means ±SD. Three/ four-cell stage cysts were analyzed, as at least 20 cysts/experiment in at least two independent experiments. *p < 0.05;**p < 0.01. *Bottom right*: Immunoblotting for expression levels of IRSp53, GFP-IRSp53 wild-type and mutants, and vinculin.

The direct interaction between IRSp53 and RAB35 suggested that these two proteins might function together in the regulation of PODXL localization during cystogenesis. Thus, perturbation of their association should affect PODXL trafficking to the AMIS and lumenogenesis. To test this possibility, structure–function rescue experiments were performed. We lentivirally transduced IRSp53 mutants in each of its critical domains (i.e., I-BAR, CRIB, SH3, PDZ-binding domains)^14^ into *IRSp53*-silenced MDCK cells and *IRSp53*-KO Caco-2 cells and perfomed cystogenesis assays to examine PODXL and aPKC localization. Expression of the I-BAR-4K mutant that does not bind to RAB35 failed to rescue the localization of PODXL to a single spot at the AMIS in these MDCK cells (Fig. 5E). This mutant was still present on the apical lumen, albeit its localization was more diffuse and less restricted than wild-type (WT) IRSp53 (see also below). Similar findings were obtained upon expression of I-BAR-4K in Caco-2 *IRSp53*-KO cells (Supplementary Fig. S5A). Furthermore, in the Caco-2 *IRSp53*-KO cells, analysis of the IRSp53 mutants showed that among the IRSp53 domains that were required for its activity, the I-BAR and SH3 domains were essential for correct IRSp53 localization at the apical lumen (Fig. 6A). Thus, IRSp53 required its functional I-BAR domain for its role in lumen formation in these epithelial cells. Molecularly, both its binding to negatively curved, phosphatidylinositol 4,5-bis phosphate (Ptdins(4,5)P_2_)-rich membranes (which is compromised by the mutations inserted^34^), and its interactions with RAB35 would appear to account for these findings. Consistent with this, *IRSp53* ablation impaired RAB35 localization at the nascent AMIS during the early phases of cystogenesis (Fig. 6B), which strengthened the concept that IRSp53 acts as an assembly platform that drives the correct localization of RAB35, and possibly its activation, at this site.

**Figure 6.**
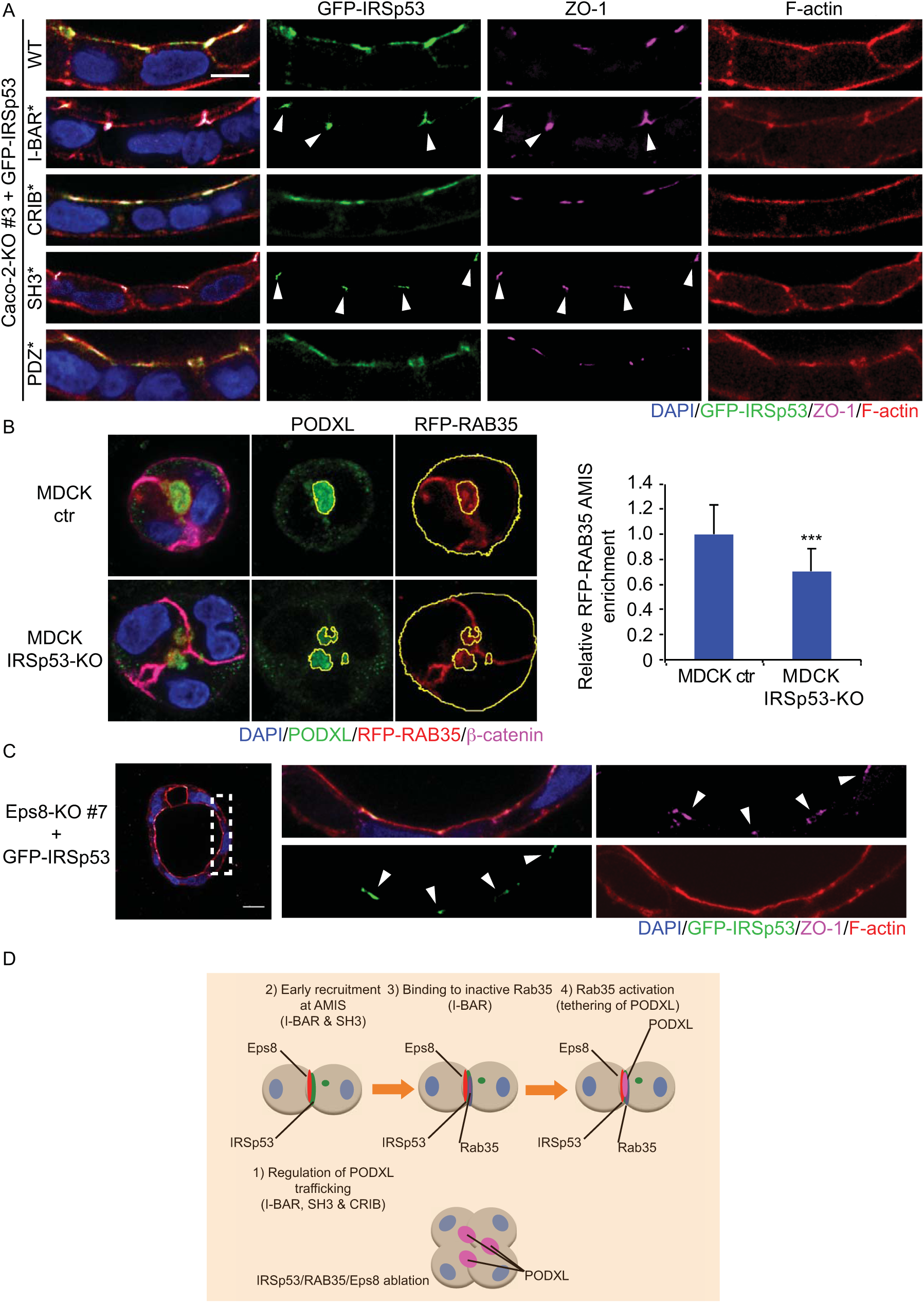
Membrane- and SH3-mediated interactions guide the correct IRSp53 localization and activity. A) Caco-2 *IRSp53*-KO #3 cells were stably infected with murine GFP-IRSp53 WT, I-BAR*, CRIB*, SH3*, PDZ*, and were embedded as single cells in Matrigel/ collagen matrix and fixed after 7 days. The cysts were processed for epifluorescence to visualize GFP-IRSp53 (green) and stained with an anti-ZO-1 antibody (magenta), rhodamine-phalloidin to detect F-actin (red), and DAPI (blue). Arrowheads, re-localization and enrichment of IRSp53 I-BAR* and SH3* at ZO-1–labeled cell–cell junctions. Scale bar, 10 µm. B) IRSp53 is required for correct RAB35 localization at the AMIS. *Left*: MDCK wild-type control (Ctr) or *IRSp53*-KO cells stably expressing RFP-RAB35 were seeded as single cells on a Matrigel layer and left to grow for 24 h. The cysts were fixed and processed for epifluorescence to visualize RFP-RAB35 (red) and stained with anti-PODXL (green) and anti-β-catenin (magenta) antibodies, and DAPI (blue). PODXL staining at the AMIS and RFP-RAB35 staining were analysed using masks (in yellow) to quantify RFP-RAB35 localization at the AMIS. *Right*: Quantification of recruitment of RFP-RAB35 at the AMIS, as the ratio of RFP-RAB35 signal at the AMIS/ RFP-RAB35 signal out of the AMIS. Data are means ±SD. At least 20 cysts/ experiment were analyzed in two independent experiments. ***p < 0.001. C) Caco-2 *Eps8*-KO #7 cells were stably infected with murine GFP-IRSp53 wild-type, embedded as single cells in Matrigel/ collagen matrix, and fixed after 7 days. The cysts were processed for epifluorescence to visualize GFP-IRSp53 (green), stained with an anti-ZO-1 antibody (magenta), rhodamine-phalloidin to detect F-actin (red), and DAPI (blue). Right panels represents 4× magnification of the area depicted by the dashed square (Left). Arrowheads, re-localization and enrichment of IRSp53 at the ZO-1–labeled cell–cell junctions. Scale bar, 20 µm. D) Schematic model of the functional interactions of IRSp53 with RAB35 and EPS8 at the AMIS during the early phases of cystogenesis, and the IRSp53 domains required for its localization and activity on PODXL trafficking.

The structure-function analysis further indicated the requirement of additional domains of IRSp53 for correct cyst morphogenesis. Indeed, IRSp53 mutations in either the CRIB-PP region (required for IRSp53 binding to active CDC42^12, 13, 18, 24^) or the SH3 domain (which mediates interactions with a variety of IRSp53 binding partners^12, 13, 17, 18, 21, 24, 35^) failed to rescue the defective distribution of PODXL in MDCK cells and lumen formation in Caco-2 cells, respectively, that arose from the IRSp53 ablation (Fig. 5E and Supplementary Fig. S5A). It is of note that impairing the function of the SH3 domain also altered the luminal localization of IRSp53 (Fig. 6A). This was particularly evident in the Caco-2 cysts. In this system, IRSp53 showed a restricted, although evenly distributed, localization along the apical, luminal side. Conversely, an IRSp53-SH3-defective mutant focally accumulated to tight junctions, where it colocalized with ZO-1, with its continuous apical distribution lost (Fig. 6A). These results indicate that in addition to the I-BAR interaction with RAB35, SH3 interactors are also likely to be important in coordination of IRSp53 localization and activity. Among these, the actin capping and bundling protein EPS8 was previously shown to form a stoichiometrically stable complex with IRSp53 *in vivo* and to have the highest affinity of association among the IRSp53 interactors^13^. Thus, we investigated the involvement of EPS8 together with IRSp53 in orchestration of the correct lumenogenesis program. Here, we show that: (a) EPS8, like IRSp53, is recruited early at the AMIS (Supplementary Fig. S5B) and has an apically restricted luminal localization in MDCK cells (Supplementary Fig. S5C) and Caco-2 cysts (Supplementary Fig S5D); b) loss of EPS8 in Caco-2 cells leads to the formation of cysts with multiple lumens (Supplementary Fig. S6A, B); c) EPS8 is required for the correct localization of IRSp53 (Fig. 6C).

### IRSp53 shapes and ensures the continuity of the plasma membrane at nascent lumens

Collectively, our findings argue that through concomitant regulation of polarized trafficking and by acting as a scaffold for the assembly of apically restricted RAB35 and EPS8 complexes, IRSp53 has an important role in the control of epithelial tissue polarity and lumenogenesis *in vitro* (Fig. 6D). However, IRSp53 is also expected to sense and control membrane curvature and local phospholipid composition through its I-BAR domain, which will ultimately impact on the ultrastructural organization and shape of the plasma membrane at the nascent AMIS. To investigate this possibility, we performed correlative light and electron microscopy analysis in the very early phases of cystogenesis. We focused on the two-cell stage of MDCK cells plated onto a Matrigel cushion. Comparison between the WT control and the *IRSp53*-KO cells showed that loss of IRSp53 delays or blocks the elimination of cytoplasmic bridges between newly formed apical domains at the AMIS (Fig. 7A, B). Also, albeit less frequently, this impacted on the intercellular luminal space (Fig. 7A), without any effects on the number and morphology of desmosomal junctional structures (data not shown), and resulted in the formation of ectopic lumens (cytoplasmic membrane vacuoles with features of apical domains) (Supplementary Fig. S7). Of particular significance, there was an increase in the number of ‘cytoplasmic bridges’ that interrupted the separation of the luminal (apical) plasma membrane of two adjacent *IRSp53*-KO cells (Fig. 7A, B). These were also visible in tomographic 3D reconstructions (Fig. 7B; Movie S2). Thus, IRSp53 is needed to ensure the integrity and structural organization of the plasma membrane at the AMIS. The increased number of membrane interconnections caused by the loss of IRSp53 might generate distinct ‘mini-lumens’ and plasma membrane targeting sites. These defects together with altered polarized trafficking of PODXL carriers might lead the accumulation of PODXL in multiple foci, which would eventually evolve into multiple lumens.

**Figure 7.**
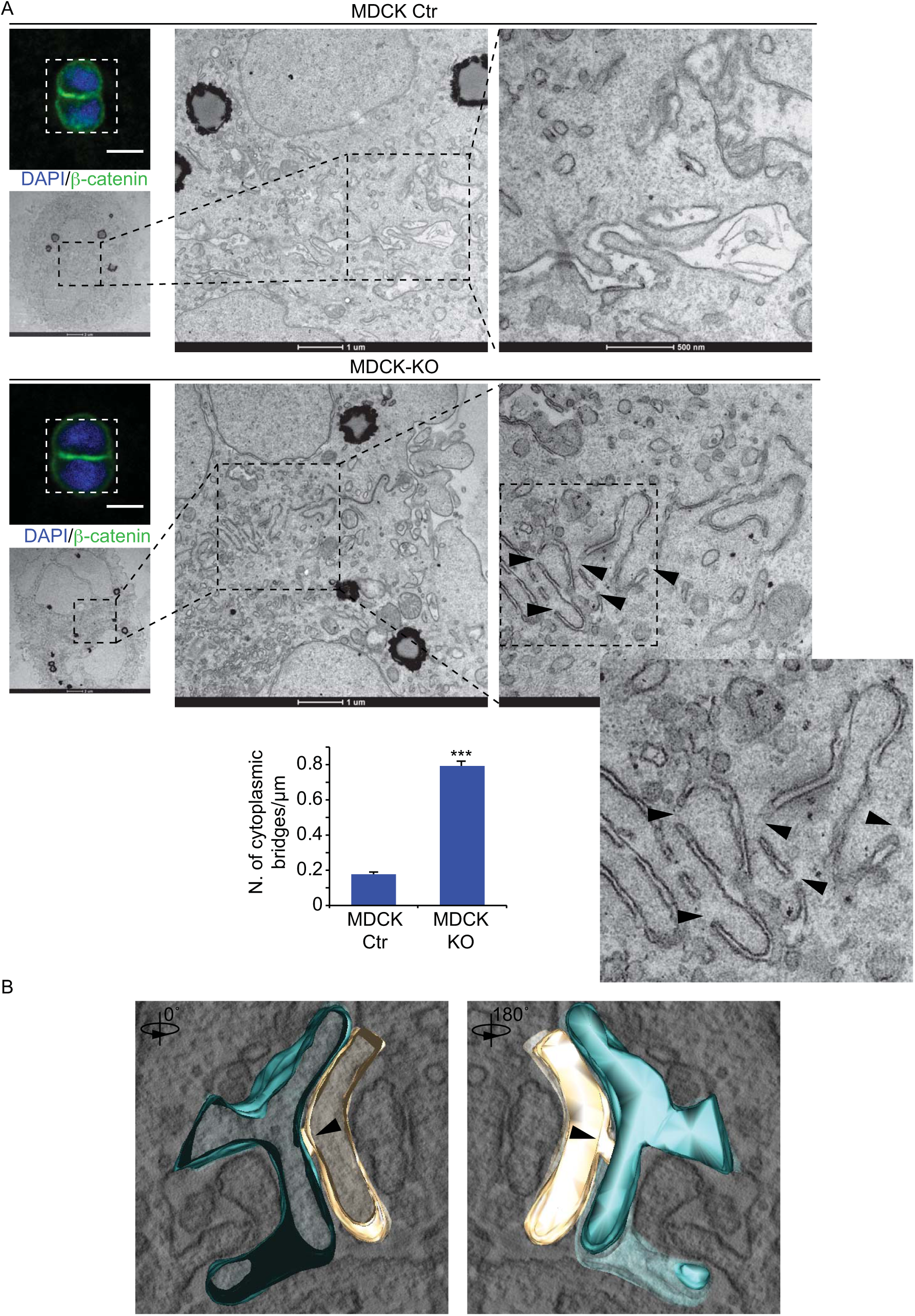
Loss of IRSp53 alters the opposing plasma membranes at the nascent AMIS. A) MDCK Ctr (top) and *IRSp53*-KO cells (middle) were seeded as single cells on Matrigel-coated gridded coverslips. The cysts were fixed 16 h after seeding, stained with anti-ß-catenin (green) and DAPI (blue). Two-cell stage cysts were initially identified on grids by confocal microscopy. Scale bar, 10 µm. Samples were then processed for electron microscopy, with images of the corresponding cells shown at the indicated magnifications. Arrowheads, inter-cytoplasmic bridges of the plasma membranes along the AMIS. *Bottom*: Quantification of the inter-cytoplasmic bridges/µm. Two-cell stage cysts were analyzed (Ctr, n = 6; KO, n = 6). In all, 383 (Ctr) and 310 (KO) different fields were counted from serial sections along the Z-axis. Data are means ±SEM. ***p < 0.001. B) Images of a three-dimensional tomographic reconstruction of an inter-cytoplasmic bridge at the opposing membranes of MDCK *IRSp53*-KO cells during early cystogenesis (see also Movie S2). Arrowheads, inter-cytoplasmic bridge.

### IRSp53 is required for kidney development and morphology

To determine whether the functional activity of IRSp53 is retained and physiologically relevant in living organisms, zebrafish lines were generated that carried a mutation in the fish paralog, *baiap2a* (b2a), and morpholinos were used to ablate *baiap2b* (b2b) (see below). Moreover, we analyzed tissue organization in *IRSp53*-KO mice.

In both of these models, IRSp53 showed prominent expression and peculiar luminal localization in the kidneys. In the renal tubules of adult mice, IRSp53 was prominently expressed and enriched at the luminal side (Fig. 1C). A similar, restricted pattern of luminal apical expression was seen in early stage pronephric ducts in developing fish embryos (Fig. 1D). The analysis was therefore focused on this organ, and specifically on its tubular architecture. In zebrafish, gene duplication has led to the generation of two distinct, but highly related, IRSp53 paralogs known as *baiap2a* (b2a) and *baiap2b* (b2b). These two gene products are expressed at variable levels in the developing embryo, with b2a as the more expressed (Supplementary Fig. S1B). A mutant zebrafish line was generated in which the gene product encoding for b2a was disrupted through introduction of a chemically mediated nonsense mutation (Supplementary Fig. S1C), with that for b2b targeted using a mixture of splice and translation blocking morpholinos (Supplementary Fig. S1C). We also characterized an antibody that can recognize both b2a and b2b, at least when ectopically expressed in mammalian cells (Supplementary Fig. S1D). Of note, the loss of b2a in the mutant line caused a large reduction in the expression of IRSp53 in the pronephric ducts, which was consistent with b2a being more expressed than b2b, albeit we cannot rule out that b2b was also present (Fig. 1D).

To visualize the pronephric duct during development, the WT and mutant b2a lines were crossed with Tg(CldnB::GFP). This transgenic line expresses EGFP under the control of the promoter of ClaudinB, which is localized in various epithelial structures and prominently labels the developing kidneys^39^. Individual depletion of the single *baiap2* genes, so in the b2a mutant line or by injection of the b2b-morpholino in WT embryos, did not interfere with normal pronephric duct development (Fig. 8A, Supplementary Fig. S8A, B and Fig. S9A). However, injection of b2b morpholinos in the b2a mutant lines caused alterations in the structure of the pronephric ducts. These ducts were less compact and dense (Supplementary Fig. S8A), with irregular or multiple lumens (Supplementary Fig. S8B) and frequently (in ∼ 67% of embryos) showed the formation of ectopic ClaudinB-positive structures emerging from their basal surfaces (Fig. 8A, Supplementary Fig. S8B, Fig. S9A and Movie S3). The multiple lumens and ectopic pseudo-lumenized structures characterized by irregular and diffused acetylated-tubulin and F-actin distributions (Supplementary Fig. S8B and Fig. S9A) were reminiscent of the defects observed in mammalian organoids devoid of IRSp53 caused by perturbation of the correct polarity program.

**Figure 8.**
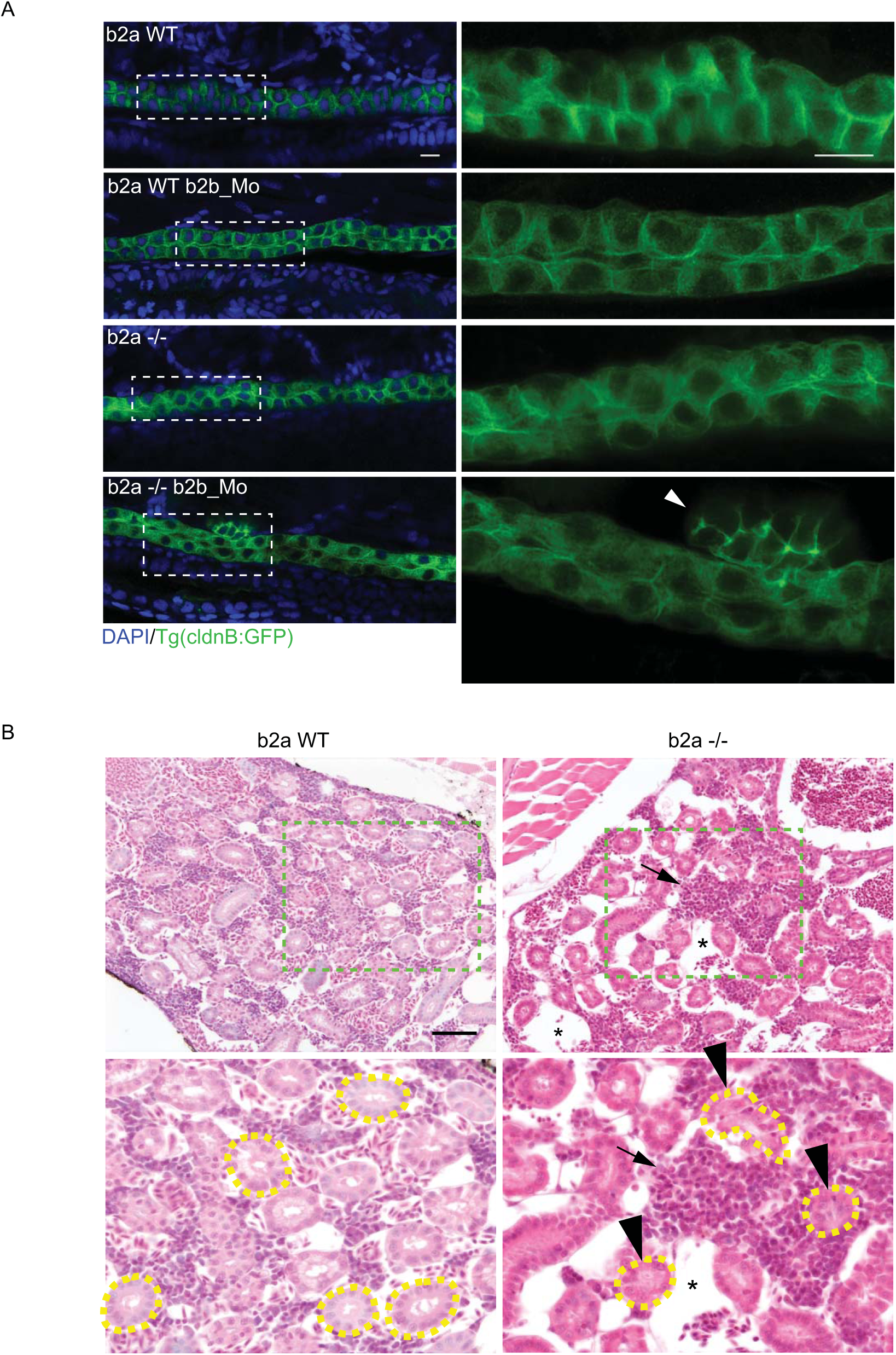
Genetic removal of IRSp53 leads to morphological defects in pro-nephric ducts and in adult kidneys in zebrafish. A) Digital light sheet confocal images of 72 hpf pronephric ducts. Embryos obtained from either wild-type (WT) or *baiap2a* mutant females in a Tg(CldnB:GFP) genetic background were treated with scrambled morpholino (b2a WT; b2a -/-) or with spliced and translation blocking morpholinos (b2a WT b2b_Mo; b2a -/- b2b_Mo), and fixed and mounted in agarose. Samples were stained with an anti-GFP antibody (green) and DAPI (blue). Z-stack projections (magnification, 3.4×) are shown on the right for the dashed, rectangular boxes on the left. Arrowhead, extra-duct structure in the b2a -/- b2b_Mo embryo. Scale bars, 10 µm. Two-thirds of the b2a -/- b2b_Mo embryos showed such defects (n = 8/12). B) Hematoxylin and eosin staining of adult zebrafish kidneys. Adult wild-type (b2a WT) and b2a -/- strains were fixed, decalcified, and paraffin embedded. *Bottom*: Magnification (2×) of the light green dashed boxes from the upper images. Renal tubules are surrounded by yellow dashed lines. Arrowheads, abnormal disarrayed and not patent tubules in the b2a -/- sample, characterized by the irregular distribution and shape of the nuclei; asterisks, vascular lacunae; arrows, infiltrating cells. In all, 70% of the b2a-/- samples show such defects (n = 7/10) *versus* ∼10% of the b2a wild-type (n = 1/9). Scale bar, 50 µm.

Despite the absence of an overt phenotypic alteration in the pronephric ducts of singly mutated b2a embryos, we reasoned that in adult life, aging-dependence or environmental stresses might combine with the genetic alterations to cause phenotypic perturbations that are not visible during development. Consistent with this, histological analysis of adult zebrafish kidneys at various ages (e.g., 4, 8 months) showed several morphological defects in the kidneys of the adult b2a mutant line (Fig. 8B). The general architecture of the organs of the b2a mutants was altered, with a decrease in the density of the tubules and a relative increase in the epithelial-to-hematopoietic cell ratio, compared to those of age-matched WT animals (Fig. 8B). This was accompanied by vascular lacunae and defects in the architectural organization of the renal tubules, which were frequently coarsed or lacked an overt lumen (Fig. 8B).

Prompted by these observations, we examined whether there might also be similar renal architectural alterations in the *IRSp53*-KO mice. *IRSp53*-KO mice have a partially penetrant mid-gestation lethality that is due to defects in placental morphogenesis, as previously reported (data not shown; and ^37^). Some 20% to 25% of these mice, however, are born fertile and devoid of gross tissue dysmorphology^12, 38, 39^. Histopathological and immunohistochemical analyses of renal tissues of the *IRSP53*-KO mice, however, showed some relatively mild, but evident and recurrent, alterations that were reminiscent of those in the kidneys and tubules of the adult B2a mutant zebrafish (Fig. 9, Supplementary Fig. S9B). The overall morphology of the *IRSp53*-KO mouse kidneys was defective, with thinning of the cortex despite preserved glomerular density. The renal proximal tubules showed irregular contours that resulted in ill-defined lumina lined by irregularly spaced epithelial cells with prominent nuclear dysmorphism (Fig. 9A). Similar architectural alterations were seen in the distal tubules. Focal periodic acid Schiff–positive urinary casts were detected in the *IRSp53*-KO mouse kidney parenchyma (Supplementary Fig. S10A). Additionally, the glomeruli of these *IRSp53*-KO mice showed alterations in the capillary tufts, because of multifocal pseudo-aneurismatic vascular dilations and/or mesangial cell proliferation (Fig. 9A). The severity of these phenotypes was scored using a histological alteration score (Fig. 9B, Supplementary Fig. S9B). Immunohistochemistry analysis of both apical and basal epithelial renal markers further highlighted defects in the epithelial tubular cell apical–basal polarity, as shown by the following findings (Supplementary Fig. S10B-D): (i) ZO-1, which labeled tight apical junctions in most tubules of the control kidney, was diffuse or reduced in *IRSp53*-KO kidney tubules (Supplementary Fig. S10B); (ii) similarly, the apical label aPKCζ was diminished, and was sometimes absent from the lumen of mutant animals (Supplementary Fig. S10C); (iii) endomucin, which localizes to the basal side of tubules in the controls, remained largely basal when expressed, but was irregularly distributed and sometimes absent in the *IRSp53*-KO tubules (Supplementary Fig. S10D).

**Figure 9.**
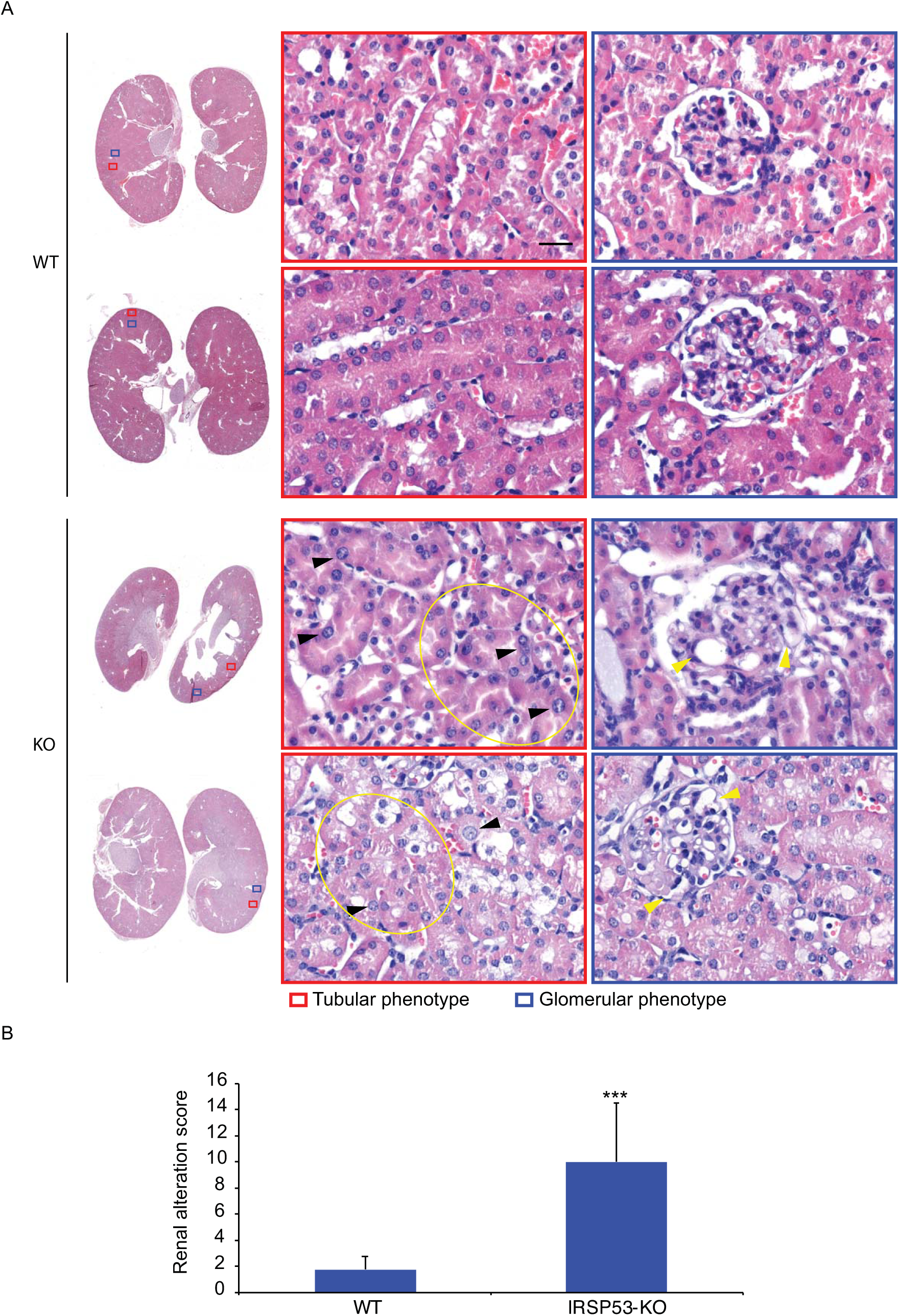
Genetic removal of IRSp53 leads to defects in murine kidney morphology. A) Hematoxylin and eosin staining of kidneys explanted from wild-type (WT) and *IRSp53*-KO adult mice were fixed, processed and paraffin embedded. *Left*: Low magnification images. *Right*: magnified images of the red (tubular phenotype) and blue (glomerular phenotype) boxes on the left. Black arrowheads, abnormal irregular distributions and shapes of the nuclei in the *IRSp53*-KO kidneys; yellow ellipses, areas with disarrayed and convoluted tubules; yellow arrowheads, glomerular tuft microcystic alterations. Scale bar, 50 µm. B) Quantification of the renal alteration score. Kidneys from wild-type (WT) and *IRSp53*-KO mice were analyzed and scored for defects in renal morphology (see also Figure S9B and Methods). Data are means ±SD. WT, n = 8 (4M, 4F); KO, n = 16 (6M, 10F). *** p < 0.001.

Altogether these data indicate that IRSp53 loss results in a set of subclinical morphological and histological aberrancies in the developing kidney and in adult renal tubules in zebrafish and mice, by impinging on, at least in part, the correct establishment or maintenance of epithelial polarity.

## Discussion

During the development of tubular or glandular structures of epithelial tissues, and specifically of kidneys, *de-novo* formation of a polarized central lumen is a key step that requires spatiotemporally coordinated interplay between actin cytoskeleton dynamics and targeted exocytosis of apical cargos, such as PODXL^40^. Although the molecular players that coordinate these events have been intensely studied^41^, we still do not have a complete mechanistic understanding of this process. Additionally, whether the knowledge gleaned from the analysis of *in-vitro* morphogenesis of multicellular systems, such as cysts derived from MDCK or Caco-2 cells^1, 28, 42–47^, is physiologically relevant *in vivo* for the development of complex organs has not been systematically tested.

Here, we show that IRSp53, a CDC42 effector that can sense Ptdins(4,5)P_2_-rich negative membrane curvature^11, 12, 14, 24, 26, 27^, acts as a platform for the assembly, integration, and spatial restriction of multiple complexes that are needed for correct formation of the polarized single lumen in various epithelial systems *in vitro*. We further show that IRSp53 is critical in the shaping of the plasma membrane, and to ensure its integrity and continuity in the nascent lumen (see Supplemntary Fig. S11). These functional roles in *in-vitro* systems are mirrored by the requirement for IRSp53 for correct morphogenesis and structural organization of kidney tubules in mice and zebrafish, where it controls the appropriate distribution of different polarity determinants.

Topologically, IRSp53 is apically restricted at the luminal membrane side of tubular and glandular epithelial tissues in human, murine, and zebrafish, and in 3D epithelial acini *in vitro*. This apically restricted localization is particularly prominent in adult murine and human kidney tubules, and in the pronephric ducts of zebrafish during development. Depletion of IRSp53 disrupts cystogenesis *in vitro*, which leads to multi-lumen formation and inversion of polarity. A remarkably similar set of alterations, which includes formation of multiple lumens and of ectopic structures that have lost their stereotypical apical–basal morphology, are also observed in the pronephric duct after genetic removal of the two zebrafish paralogs of mammalian IRSp53. Renal tubules in *IRSp53*-KO mice are convoluted, not patent, and frequently display an aberrant distribution of the epithelial cells that normally surround the lumen in an orderly and stereotypical pattern. The latter phenotypes correlate with incorrect distribution of various apical determinants, including ZO-1 and aPKC. Collectively, these findings support the crucial involvement of IRSp53 in renal lumenogenesis and tubule morphogenesis across different species.

The relatively mild tubular defects in adult organisms appears to reflect the plasticity of the epithelial morphogenesis processes, which can use multiple, and often redundant, molecular pathways to ensure maintenance of the correctly organized structure and the functional activity of the tissues. In keeping with this concept, the genetic loss of another member of the I-BAR family proteins, known as ‘missing-in metastasis’ (MIM, also known as MTSS1), was shown to impact on adult kidney architecture through regulation of intercellular junctions. MIM^-/-^ mice show a progressive kidney pathological diseased state with dilated tubules, glomerular degeneration, and fibrosis. This set of alterations is accompanied by polyuria at the systemic level, whereas increased intercellular junctional spaces are detected by ultrastructural electrom microscopy analysis^48^. Thus, different members of the I-BAR family proteins contribute to the proper structural organization of adult murine kidney, with these partly acting in a redundant fashion^48^. It is also of note that some of the alterations due to loss of IRSp53 are observed in mice devoid of key components of the molecular network of the IRSp53 interactors. A case in point is CDC42, which has a pivotal role in virtually all polarized cell processes in diverse epithelial and non-epithelial tissues by acting as a central signaling hub that connects actin dynamics, vesicular trafficking, and membrane remodeling^1, 49, 50^. Indeed, genetic loss of CDC42 in murine kidney tubules during development leads to abnormal tubulogenesis with profound defects in polarity, lumen formation, and the actin cytoskeleton, which eventually result in premature death of these animals^51–53^. These alterations are significantly more severe than those detected in *IRSp53*-KO mice, which is in keeping with the concept that CDC42 is a central hub that coordinates and uses multiple effectors and pathways to control polarity. Among these, IRSp53 appears to contribute to interconnecting of the CDC42 axis with RAB35/PODXL trafficking and EPS8-based actin dynamics. However, other CDC42-dependent pathways, such as Ptdins(4,5)P2-Annexin-CDC42^54^, CDC42-PAR3-aPKC^55^, and CDC42/YAP1^53^ are expected to remain functional following disruption of IRSp53, and might, therefore, compensate for the lack of IRSp53.

Redundancy and compensation mechanisms of IRSp53 functions might also be at play in processes and pathways that control endocytic internalization of PODXL, inversion of polarity and Robo2-p53 axis, which, as described in the supplementary discussion, have been implicated in the correct lumenogenesis and renal tubule formation.

At the cellular level, during 3D cystogenesis, the loss of IRSp53 leads to formation of multiple lumens and reversal of polarity, albeit less frequently. The formation of multiple lumens occurs frequently following disruption of a variety of trafficking, polarity, and cytokinetic regulators. This indicates that none of these processes is essential for establishment of apical–basal epithelial polarity; rather, they all contribute to ensure the formation of the single AMIS that will eventually generate a single lumen. In the case of IRSp53, its loss is likely to impact on the optimal functionality of many of these critical cell-biology processes. Indeed, IRSp53 is recruited very early on, at the onset of AMIS formation, which appears to be as a consequence of its I-BAR domain sensing of Ptdins(4,5)P_2_-rich, negative membrane curvature that is a distinguishing early feature during *de-novo* formation of the AMIS around the midbody immediately after completion of the first cytokinesis^6, 54^. At this site, and similar to what is seen in extending protrusions of migratory cells^16, 56–60^, IRSp53 accumulates together with its interactor, the actin capping protein EPS8. This protein is expected to regulate not only local cortical actin dynamics, but also to maintain IRSp53 in an ‘open conformation’, opposing the inhibitory actions of 14-3-3^24, 61^. This might facilitate the association of IRSp53 with Ptdins(4,5)P_2_-rich membranes, and also with activated CDC42 and inactive RAB35. In keeping with this concept, removal of EPS8 not only phenocopies the loss of IRSp53 in cystogenesis assays, but also results in redistribution of IRSp53, which loses its continuous apical pattern, and becomes concentrated at tight junctions in association with ZO-1. The importance of the interaction between IRSp53 and activated CDC42 is underscored by the observation that a single-point mutation in the CDC42 binding site of IRSp53 fails to rescue the phenotype in *IRSp53*-KO MDCK and Caco-2 cells. Additionally, *in vitro*, IRSp53 loss mimics the removal of CDC42^28, 62, 63^, which leads to the formation of multiple lumens. Of note, the loss of CDC42 also disrupts the orientation of the spindle in dividing cells during cystogenesis^28, 62, 63^ and controls the trafficking of polarized determinants. Both of these altered processes are thought to be responsible for the aberrant formation of multiple lumens. Within this context, IRSp53, which directly binds to CDC42, might mediate some of these specific functions. Consistent with this, trafficking of PODXL was perturbed and IRSp53 localized to PODXL-positive recycling endosomes. Further, the loss of IRSp53 significantly altered the spindle orientation during Caco-2 cystogenesis, to an extent and in a fashion not dissimilar from those seen after the loss of CDC42 (data not shown). While the precise mechanisms through which IRSp53 applies this multiple-level regulation remains to be defined, the most parsimonious interpretation of our findings is that IRSp53 acts as an important CDC42 effector, during both spindle orientation and polarized membrane trafficking.

In addition to this role, the localization of IRSp53 at the AMIS and its interaction with RAB35 suggests that it is also part of a set of molecular cues that define the identity of, and ensure the structural integrity of, the future apical membrane. Here, IRSp53 might act as a stabilizing factor for RAB35, either directly through protein:protein interactions, or indirectly through sensing and enhancing the concentration of PtdIns(4,5)P2^22^, to which RAB35 is known to associate^64^. In this latter respect, it is worth noting that: (i) As for the loss of RAB35 that was recently demonstrated to function as a PODXL tethering factor at the AMIS^10^, also the loss of IRSp53 disrupts the correct apical distribution of PODXL; (ii) the binding to RAB35 and Ptdins(4,5)P2-rich membrane is mediated by the I-BAR domain. Additionally, a mutant in the key positively charged patches of this domain that disrupts the interaction of IRSp53 with Ptdins(4,5)P_2_-rich lipid bilayers^36^ as well as the binding to RAB35 failed to correctly localize apically and to rescue the multi-lumen phenotype in the *IRSp53*-KO cysts; (iii) *IRSp53* ablation impairs RAB35 localization at the nascent AMIS, which demonstrates that IRSp53 drives the correct localization of RAB35. An equally plausible, although not mutually exclusive, possibility is that in addition to binding to inactive RAB35 along the AMIS, IRSp53 might associate with regulators of the RAB35 GTPase, including through its GEFs or GAPs, to thus contribute to the regulation of its activity. Our proteomic studies using IRSp53–BirA approaches that can detect transient interactions^65^, however, failed to identify any of these proteins (deposited at http://www.peptideatlas.org/PASS/PASS01464 for details see Materials and Methods), which would argue against this mode of action.

Ultimately, the multiple functional roles of IRSp53 are likely to be integrated with its sensing and shaping of the flat plasma membrane at the nascent lumen via its I-BAR domain. Here, IRSp53 appears to be crucial in the control of the integrity, continuity, and shape of the opposing plasma membrane. Indeed, its loss leads to the formation (or preservation) of bridges between the apical domains of the plasma membrane, which interrupt the continuity of the nascent lumen (Supplementary Fig. S11). It appears reasonable to propose that it is exactly this structural, membrane role in combination with its control of the trafficking of key polarity determinants that accounts for its involvement in lumen formation and tubule morphology.

Whatever the case, all in all, our findings are consistent with a multifaceted role for IRSp53 that is crucial to ensure the structural integrity and shape of the plasma membrane, and to coordinate the trafficking of apical proteins to the AMIS. They further support the concept that IRSp53 acts as a platform where trafficking determinants (e.g., RAB35), actin regulatory proteins (e.g., EPS8), and the small GTPase that is critical for polarity (i.e., CDC42) are assembled for the correct coordination of diverse processes during tubule morphogenesis and the establishment of renal tubular architecture in different species.

## Acknowledgments

We thank: Fernando Martin-Belmonte and Gregory Emery for critically reading the manuscript. for Chiara Luise and Giovanna Jodice for technical assistance with immunohistochemistry experiments; Ilaria Costa and the IFOM Imaging Facility for technical assistance in the design and performing of the confocal imaging; Ghazaleh Saberamoli for assistance with graphic design. This study was supported by: the Associazione Italiana per la Ricerca sul Cancro (AIRC-IG#18621 to GS, AIRC-0IG#22145 to CT, and 5XMille #22759 to GS and CT); the Italian Ministry of University and Scientific Research (MIUR) to GS (PRIN: Progetti di Ricerca di Rilevante Interese Nazionale – Bando 2017#2017HWTP2K); the Italian Ministry of Health (RF-2013-02358446) to GS. SB was supported by Fellowships from AIRC.

## Materials and Methods

A detailed description of the Materials and Methods is included as the Supplementary Text in the Supplementary Information section.

## Author contributions

SB, AR, SM, AD designed and performed all the experiments and edited the manuscript; DC aid in generating cell lines and in the analysis of immunofluorescence; AD, and GvB and AM perfomed all of the electron microscopy studies, IF, GB, FP, SP, GV, and CT collected the specimens, and performed and interpreted all the immunohistoichemical studies in human murine and zebrafish tissues: AC, AB performed and analyzed the mass spectrometry with SB. AO performed and analyzed the light imaging data; AD, GS conceived the whole study, wrote the manuscript, and supervised the whole project.

## Data Availability Statement

The authors declare that all of the data that support the findings of this study are available within the paper and its Supplementary Information files, and from the corresponding authors upon reasonable request.

Proteomic data have been deposited and are available at the PeptideAtlas repository (http://www.peptideatlas.org/PASS/PASS01464)

