## Supplementay Text and Figures for "IRSp53 shapes the plasma membrane and controls polarized transport at the nascent lumen during epithelial morphogenesis"

### SUPPLEMENTARY INFORMATION

**- Supplementary Discussion**

**- Materials and Methods**

**- Supplementary Figures and Supplementary Movies legends**

**- Original uncropped immunoblots and gels for all figures**

#### SUPPLEMENTARY DISCUSSION

##### On IRSp53 and RAB35 endocytosis and trafficking in kidney development

IRSp53 loss in addition to perturb endocytic trafficking, also specifically impacts on PODXL internalization into the vacuolar apical compartment after $Ca^{++}$ removal. Little is known as to the routes that mediate PODXL endocytosis under these conditions. The recently reported involvement of RAB35 in controlling PODXL trafficking in cystogenesis[1,2] and the stablished function of this GTPase in Clathrin-mediated endocytosis (CME)[3-5] suggest that PODXL may enter, at least in part, through this internalization pathway. These findings further point to the concept that disruption of the RAB35- CME pathway may affect the proper development of epithelial polarity in vitro as well as in in vivo. Notably, mutations in Lowe oculocerebrorenal syndrome protein (OCRL), a direct interactor/ effector of RAB35[6] implicated in the control of phosphoinositide switch during Clathrin-mediated endocytosis , also impacts on proximal kidney tubule architectural organization [7]. OCRL is an inositol polyphosphate 5-phosphatase that regulates phosphoinositide homeostasis along the endo-lysosomal pathway [8]. Loss of function of OCRL in conjunction with tissue-specific ablation in the renal tubules of its paralog type 2 inositol polyphosphate-5-phosphatase (INPP5B) results in profound dysfunction of the cells lining the kidney proximal tubules[7]. This is primarily due to defective re-absorption of the majority of the daily filtered load, as a consequence of impaired clathrin-dependent endocytosis of solutes, as well as defective transporters and channels. The tubular defects in *IRSp53*-KO mice, however, appear to be linked to disrupted tubular architecture, rather than perturbations in clathrin-dependent endocytosis, which is not affected in the *IRSp53*-KO epithelial cells (data not shown),. Importantly, however, recent work implicated this protein in non-clathrin-dependent internalization pathways [9] suggesting that a more complex interplay among different endocytic routes is at play during lumenogenesis.

**On IRSp53 and inverted polarity**

One striking albeit quantitatively limited phenotype due to IRSp53 loss is the inversion of polarity. This phenotype has previously been reported in cysts that expressed a dominant-negative form of RAC1[10], cysts with depletion of the small GTPases ARF6[11] or RAB35 [2], and cysts grown in the presence of functional-blocking antibody against β1-integrin[12]. All of these proteins are connected with signaling pathways from the extracellular matrix to the cell interior, starting from the binding of matrix components to integrin dimers, which leads to ARF6 and RAC1 activation and further signal transduction [11,12]. As IRSp53 is an interactor of RAC1[13] and a component of the adhesome[14], and as its depletion reduces cell spreading and focal adhesion formation[15], it is possible that the inverted phenotype seen in these *IRSp53*-KO cysts is the consequence of incorrect β1-integrin recycling, that might be caused by dysregulation of the interplay between RAB35, CDC42, and RAC1.

**On Robo2-IRSp53 signaling**

IRSp53 has also been reported to interact with Robo2 and to be implicated in the aberrant formation of renal cysts[16]. Robo2 is a cell surface receptor for the for the repulsive guidance cue Slit that in central nervous system acts in the control of axonal guidance and migration[17,18]. During renal tubule development, Robo2 deficiency results in cystic kidneys, and the cyst cells showed defective cilia and polarity defects in tubular epithelium[16]. This, in part, is the result of failure of the differentiation program of cysts cells, which continue to proliferate[16]. Molecularly, Robo2 was reported to function by regulating the levels of p53 and ciliary polarity complexes, including PAR3, aPKC, and ZO-1[16]. It was also shown that Robo2 binds to IRSp53 via the I-BAR domain[16]. This interaction appears essential for the phosphorylation of the p53-specific, E3 ligase MDM2, and to maintain p53 homeostatic levels[16]. Consistently, ablation of Robo2 or IRSp53 caused a reduction in MDM2 phosphorylation, the aberrant elevation of p53 that, in turn, triggered a DNA-damage-mediated, p53-dependent senescent program, leading to defect in epithelial development, including altered ciliogenesis, polarity and differentiation[16]. Using our IRSp53 CRISPR MDCK cells or renal tissues derived from IRSp53-KO mice, however, we failed to detect any alterations in the levels of DNA damage, of total and phosphorylated p53 or MDM2 (not shown). Moreover, while some micro-cysts were detected within the glomerular tufts, IRSp53-KO kidneys did not display any signs of an overt cystic disease as observed in Robo2 null mice. These results indicate that IRSp53 is not strictly essential for the functionality of the Robo2-p53 axis. It is possible that other I-BAR containing proteins may compensate for the loss of IRSp53 in this process and further experiments are required to address this issue.

#### MATERIALS AND METHODS

All animal experiments were conducted in accordance with national guidelines and were approved by the ethics committee of the Animal Welfare Office of the Italian Health Ministry and conformed to the legal mandates and Italian guidelines for the care and maintenance of laboratory animals (autorizzazione n. 598/2015-PR del 26/06/2015).

##### 1.1 Antibodies, plasmid and reagents

The antibodies used were: anti-IRSp53 mouse (IF and WB on MDCK and Caco-2, IHC on human tissue samples; IFOM); anti-IRSp53 rabbit (IHC on mouse tissue samples; SIGMA HPA023310); anti-IRSp53 rabbit (IF and WB on zebrafish Baiap2a and Baiap2b, IHC on zebrafish embryo; gift from Hans-Jürgen Kreienkamp, University of Hamburg); anti-vinculin (Sigma, V9131); anti-GFP rabbit for WB (Sigma, G1544); anti-GP135/podocalyxin (DSHB, Iowa University, 3F2/D8) ; anti

Eps8 (BD-Transduction Lab, 610144); anti-RAB7 (Cell Signaling, 9367); antiRAB11 (Cell Signaling, 5589); anti-ZO1 rabbit (Invitrogen, 402200); anti-aPKC rabbit (SCBT sc-216); anti-ß-catenin (Sigma, PLA2030); anti-Myc 9E10 mouse (IFOM); anti-EEA1 (SCBT, sc-6415); anti-LAMP1 (Sigma, L1418); anti-Giantin (BioLegend, 924302); anti-pERM (Cell Signaling, 3141L); anti-Flag (Sigma, F3165); anti-acetyl-α-tubulin (IF on Zebrafish embryos; Abcam, Ab24610); anti-Endomucin (Abcam, clone V.7C7.1 ab106100); anti-acetyl-α-tubulin (IHC on Tissue; Cell Signaling, #5335); anti-mouse ZO-1 (IHC; Genetex, GTX108592).

pFUW-GFP, pFUW-GFP-IRSp53 WT and mutants were a gift from Hans-Jürgen Kreienkamp, University of Hamburg; pRK5 Flag-IRSp53 was a gift from Sonia Krugmann; pRRL-GFP-Eps8 was generated by sub-cloning GFP-Eps8 in pRRL lentiviral vector[19]; mRFP-Rab7 was a gift from Ari Helenius (Addgene plasmid # 14436); GFP- and RFP-RAB35 WT, -RAB35 S22N and -RAB35 Q67L were kind gifts from Cécile Gauthier-Rouviere, Centre de Recherche en Biologie cellulaire de Montpellier; pGEX-RAB35-S22N, RAB35-Q67L were generated by sub-cloning GFP-RAB35-Q67L, -RAB35-S22N in pGEX6P1 vector; RFP-PODXL was a gift from Fernando Martín-Belmonte, Centro de Biología Molecular Severo Ochoa, Madrid.

MDCK IRSp53 shRNAmir inducible cell line were a gift from Ann Musch and were described elsewhere[15].

#### 1.2 Cell culture

##### 1.2.1 Cells culture

MDCK cells were grown in Dulbecco's Modified Eagle Medium (DMEM, Lonza) supplemented with 5% South American serum (EuroClone) and 2mM L-Glutamine (EuroClone). MDCK TetOFF cells were grown in Dulbecco's Modified Eagle Medium (DMEM, Lonza) supplemented with 5% South American serum (EuroClone), 2mM L-Glutamine (EuroClone); controls were maintained in the presence of 1 μg/ml tetracycline (Sigma) to repress IRSp53 RNAi. Caco-2 cells were grown in Modified Eagle Medium (MEM , Biowest) supplemented with 20% South American serum, 2mM L-

Glutamine and 0.1mM NEAA. HEK 293-T cells were grown in Dulbecco's modified Eagle's medium (DMEM, Lonza) supplemented with 10% South American serum (EuroClone) and 2mM L-Glutamine (EuroClone). Cells were grown at 37°C in 5% $CO_2$. HeLa cells were grown in Minimum Essential Medium (MEM Invitrogen) supplemented with 10% South American serum (EuroClone), 1% non-essential amino acids and 1% Sodium Pyruvate.

##### 1.2.2 Transfection

Transfections were performed using the calcium phosphate method or liposoluble agents JETPrime (POLYPlus), INTERFERin (POLYPlus) and FugeneHD (Promega).

293T cells were transfected using the calcium phosphate procedure. In this case DNA (10 µg for a 10 cm plate) was diluted in 439 µl of $ddH_2O$ and 61 µl of 2M $CaCl_2$ were added. This solution was added, drop-wise, to 500µl of HBS 2x (50 mM Hepes pH 7.5, 10 mM KCl, 12 mM dextrose, 280 mM NaCl, 1.5 mM $Na_2HPO_4$). Then, the precipitate was added to the cells and the medium replaced after 12-16 hours.

JETPrime (POLYPlus) was used, according to manufacturer's instruction, to transfect MDCK cells with fluorescent tagged CRISPR/CAS9 plasmid carrying sgRNA sequences, and fluorescent or tagged proteins for IF or IP experiments.

INTERFERin (POLYPlus) was used, according to manufacturer's instruction, to transfect MDCK cells for transient RNAi interference experiments.

FugeneHD (Promega) was used, according to manufacturer's instruction, to transfect CACO2 with fluorescence-tagged CRISPR/CAS9 plasmid carrying sgRNA sequences.

##### 1.2.3 CRISPR/CAS9 genome editing

MDCK and Caco-2 transiently transfected with fluorescent tagged CRISPR/CAS9 plasmid carrying sgRNA sequences (see below for sgRNA sequences and analysis of the lesions introduced)); 48 hours after transfection cells were sorted by FACS and re-cultured. Mass population were subjected to WB and IF analysis to verify the extent of KO. Clones were then derived by serial dilution. Analysis of the lesions introduced was performed by PCR.

MDCK

sgRNA:

WT 5’-GGCAGCAGGCCTGCGCCGACCCCAACAAGATCCCAGACCGCGCGGTGCAG-3’

Sequence analysis:

Allele1 5’-GGCAGCAGGCCTGCG**T**CCGACCCCAACAAGATCCCAGACCGCGCGGTGCAG-3’

Allele2 5’-GGCAGCAGGCCTGC - - CGACCCCAACAAGATCCCAGACCGCGCGGTGCAG-3’

Both mutations cause the premature termination of the protein at the end of the I-BAR domain.

Caco-2

sgRNA:

5’-AAGCAGGGCGAGCTGGAGAATTACGTGTCCGACGGCTACAAGACCGCACT-3’

Clone #3:

Allele1 5’-AAGCAGGGCGAGCTGGAGAATTACG - - - - - - ACGGCTACAAGACCGCACT-3’

Allele2 5’-AAGCAGGGCGAGCTGGAGAATTACG - - - - - - A - - GCTACAAGACCGCACT-3’

Allele3 5’-AAGCAGGGCGAGCTGGAGAATTACGTGT - - - - - - - CTACAAGACCGCACT-3’

Deletion in Allele 1 causes the translation of a protein lacking 2aa within the I-BAR (179-180). Deletion in Allele 2 causes premature termination of the protein at the end of the I-BAR domain (aa 311). Deletion in Allele 3 causes premature termination of the protein within the I-BAR domain (aa 184).

Clone #12

Allele1 5’-AAGCAGGGCGAGCTGGAGAATTACG - - - - - - - - - GCTACAAGACCGCACT-3’

Allele2 5’-AAGCAGGGCGAGCTGGAGAATTACGTGTC - GACGGCTACAAGACCGCACT-3’

Allele3 5’-AAGCAGGGCGAGCTGGAGAATTACG - - - - - - ACGGCTACAAGACCGCACT-3’

Deletion in Allele 1 causes the translation of a protein lacking 3aa (179-180-181). Deletion in Allele 2 causes premature termination of the protein within the I-BAR domain (aa 186). Deletion in Allele 3 causes the translation of a protein lacking 2aa within the I-BAR (179-180).

PAM sequence; gRNA sequence.

##### 1.2.4 Short interfering RNA (siRNA) experiments

siRNAs (small interfering RNAs) delivery was achieved by mixing from 5 to 50 nM of specific siRNAs with INTERFERin Transfection Reagent (POLYPlus) according to manufacturer's instruction. For each RNA interference experiment, negative control was performed with the same amounts of scrambled siRNAs (si CTR). Knocking down efficiency was controlled by western blot. Oligos details as follows:

IRS 302 CCAAGGAACUCGGAGACGUUCUCUU (Invitrogen)

IRS 749 CAUAGUGGCAGUUUCUGUGCCAGCA (Invitrogen)

RAB35-1 AGAAGAUGCCUACAAAUUCUU (Invitrogen)

RAB35-2 GCUCACGAAGAACAGUAAAUU (Invitrogen)

##### 1.2.5 Lentiviral infections

GFP and GFP-IRSp53 WT or mutants were transduced into MDCK, Caco-2, Huvec and mEC cells by lentiviral infection using pFUW-based plasmids. Lentivirus was produced transfecting HEK 293T cells with 25 μg of pFUW- or pLLRsin-based constructs, 9 μg ENV, 16.25 μg pMDL, 6.25 μg REV (for virus packaging). 48 and 72 hours after transfection, supernatants were collected and passed through a 0.45 μm filter. Supernatants collected at 48h and 72h were added to cells and medium was changed after 6 hours. GFP and GFP-IRSp53 WT and mutants and GFP Eps8 expression was controlled by WB and IF.

##### 1.2.6 Cyst Matrigel Culture

MDCK cells were trypsinized to a single cell suspension at $4 \times 10^4$ cells/ml in complete medium containing 2% Matrigel (BD Biosciences). Cell-medium-Matrigel suspensions (250 μl) were plated into μ-Slide 8 well ibiTreat chambers (Ibidi, 80826), precoated with 15 μl of Matrigel (10mg/ml).

For long cultures, medium was replaced every 2 days. Cells were grown at different time points before fixation in 4%paraformaldehyde (PFA).

##### 1.2.7 Cyst Matrigel/Collagen Culture

Caco-2 cells were trypsinized to a single cell suspension at $6 \times 10^4$ cells/ml and mixed with 20 mM HEPES pH7.5, 1 mg/ml Collagen I (Corning®), 50% Matrigel (BD Biosciences). Final volume was adjusted with complete medium containing. Cell-Matrix-medium suspensions (250 µl) were plated into µ-Slide 8 well ibiTreat chambers (Ibidi, 80826), and left in the incubator at 37 ˚C for at least 30 minutes to allow the mix to solidify through polymerization of Collagen I and Matrigel. The solidified mixture was then supplemented with 250 µl of complete medium. Cells were grown at different time points before fixation in 2%paraformaldehyde (PFA). For long cultures, medium was replaced every 2 days. To speed-up lumen formation in mature cysts, Cholera Toxin (0.1 µg/ml; ) was added at day 6 and cysts culture fixed at day 7.

#### 1.3 Imaging techniques

##### 1.3.1 Immunofluorescence

Cells were plated on glass coverslips (pre-incubated with 0.5% gelatin in PBS at 37˚C for 30 minutes). Cells were processed for epifluorescence or indirect immunofluorescence microscopy as follow. Cells were fixed in 4% paraformaldehyde for 10’, washed with PBS and permeabilized in PBS 0,1% Triton X-100 for 10 minutes at room temperature (RT). To prevent non-specific binding of the antibodies, cells were then incubated with PBS in the presence of 1 % BSA for 10 minutes.

The coverslips were then gently deposited, face down, on 50 µl of primary antibody diluted in PBS 1% BSA, spotted on Parafilm. After 40 minutes of incubation at RT, coverslips were transferred into 12 well plates and washed three times with PBS. Cells were then incubated for 40 minutes at RT with the appropriate secondary antibody. F-actin was detected by staining with FITC or TRITC-conjugated phalloidin (Molecular Probes) at a concentration of 6.7 U/mL. After three washes in PBS, coverslips were transferred into 12 well plates and incubated in PBS containing DAPI (1:3000) for 5 minutes at RT. Coverslips were washed three times in PBS coverslips and mounted in Mowiol (20% Mowiol (Sigma), 5% Glycerol, 2.5% DABCO (Molecular Probes), 0.02% NaN3 in PBS) and examined by fluorescent optical microscopy or in a 90% glycerol solution containing diazabicyclo-(2.2.2) octane antifade (Sigma) and examined under confocal microscope (see below for further information regarding confocal microscopes). Confocal image acquisition was performed in sequential mode to limit channel cross-talk and corrected for residual fluorescence bleed through. Images were further processed with the Image J software (Adobe).

##### 1.3.2 MDCK cysts

Cysts seeded on Matrigel into µ-Slide 8 well ibiTreat chambers were fixed at different time point after seeding in PBS 2% PFA for 20 minutes, washed 3 times in PBS 1% Glycine (quenching) and left 2 days in PBS 1x at 4 ˚C. Cysts were permeabilized in PBS + 0.5% Triton X-100 and quickly washed in IF-buffer (PBS, 0.2% Triton X-100, 0.1% BSA, 0.05% Tween). Samples were then incubated in blocking solution (IF-buffer + 5% BSA) for 1 hour and 30 minutes. After 1 wash in IF-buffer, cysts were incubated with primary antibodies in IF-buffer, overnight at 4 ˚C. Samples were washed 3 times in IF-buffer (5 minutes each) and incubated with the appropriate secondary antibody and/or phalloidin for 1 hour and 30 minutes in the dark. After 1 wash in IF-buffer (5 minutes) and 2 washes in PBS samples were incubated with Dapi (5 minutes). Samples were washed 2 times in PBS and them left in PBS for confocal analysis. Images were acquired with confocal microscopes of the Leica SP series (see below for the detailed information) equipped with a HCX PL APO 40X/0,75-1,25 oil immersion objective.

##### 1.3.3 Caco-2 cysts

Cysts embedded in Matrigel/Collagen I into µ-Slide 8 well ibiTreat chambers were fixed at different time point after seeding in PBS 2% PFA for 20 minutes, washed 3 times in PBS 1% Glycine (5 minute each), 1 time in PBS and left 2 days in PBS, 0.03 % $NaN_3$ at RT. Cysts were permeabilized in PBS + 0.5% Triton X-100 and rinsed 3 times in PBS (5 minutes each). Blocking was performed in IF-buffer (PBS 0.2% Triton X-100, 0.05% Tween 20, 0.1% BSA, 0.03% $NaN_3$) + 10% Goat serum for 1 hour. Cysts were incubated with primary antibodies in IF-buffer, overnight at 4 ˚C. Samples were washed 3 times in IF-buffer (5 minutes each) and incubated with the appropriate secondary antibody and/or phalloidin in IF-buffer for 1 hour in the dark. Samples were washed 3 times in IF-buffer (5 minutes each) in the dark, incubated with Dapi (5 minutes) and rinsed once with PBS for 5 minutes in the dark. To reduce the diffusion of antibodies, samples were fixed with 2% PFA for 9 minutes in the dark at RT. After 3 washes (5 minutes each) with PBS/glycine in the dark, samples were rinsed with PBS (5 minutes) in the dark and stored in PBS in the dark for confocal analysis. Images were acquired with confocal microscopes of the Leica SP series (see below for the detailed information) equipped with a HCX PL APO 40X/0,75-1,25 oil immersion objective.

##### 1.3.4 Podocalyxin trafficking assays

**PODXL re-localization at VAC (Vacuolar Apical Compartment)**[20,21]. MDCK cells were grown in normal medium till confluency. Monolayer were then wash 3 times in PBS 1x (w/o Ca2+) and cultured in medium without Ca2+ for 2, 5, 8 hours. Cells were then fixed in 4% PFA, permeabilized and subjected to immunofluorescence analysis. In control experiments to test the anti-PODXL antibody that recognize the N-Terminal extracellular domain of the protein (Figure SI3B), MDCK cells monolayers were pre-incubated with anti-PODXL antibody for 1h. After incubation cells were washed 3 time in cold PBS 1x and fixed in 4% PAF for 10min without permeabilization (-S -P), subjected to surface stripping with 0.5% acetic acid pH3 + 0.5 M NaCl, washed 3 time in cold PBS 1x and fixed in 4% PAF for 10min without permeabilization (+S -P) or subjected to surface stripping with 0.5% acetic acid pH3 + 0.5 M NaCl, washed 3 time in cold PBS 1x, fixed in 4% PAF for 10min, and then permeabilized (PBS 1X, 0,1% Triton X-100, 10min) (+S +P).

**1.3.5. RFP-RAB35 localization at AMIS**

MDCK cysts WT control or IRSp53-KO, stably expressing RFP-RAB35, were fixed and processed for epifluorescence to visualize RFP-RAB35 (red) and stained with anti-PODXL (green), anti-β-catenin (magenta) and Dapi (blue) and images were acquired with a Leica SP8 confocal microscope equipped with a HCX PL APO 40X/0,75-1,25 oil immersion objective. PODXL staining at AMIS and RFP-RAB35 staining were adopted to design, using ImageJ software, masks contouring the AMIS and the whole cell body (in yellow). The area of the AMIS was subtracted to the area of the well cell bodies to obtain the "non AMIS" cell area. RFP signal (Integrated density) was determined for each cell using ImageJ software in the AMIS vs "non AMIS" area. RFP-RAB35 ratio was calculated by dividing the signal at the AMIS over the signal of the "non AMIS" area.

**1.3.6 Confocal spinning-disc time-lapse microscopy.**

MDCK cells were seeded as single cells on Matrigel coated coverslips in complete medium containing 2% Matrigel (BD Biosciences). 6 hours after seeding, images were acquired (15 hours, 5 min time interval) with the UltraVIEW VoX (Perkin Elmer) spinning disk confocal unit. Images were acquired with a 60X oil immersion objective (Nikon PLANApo VC 1.4 NA). The experiments were performed using an environmental microscope incubator (OKOLab) set to 37°C and 5% $CO_2$ perfusion.

**1.3.7 Correlative light and electron microscopy (CLEM)**

MDCK cells were trypsinized to a single cell suspension at $4 \times 10^4$ cells/ml in complete medium containing 2% Matrigel (BD Biosciences). Cell-medium-Matrigel suspensions (350 µl) were plated into 35 mm dish with gridded coverslips (MatTek Corporation, P35G-1.5-14-CGRD), pre-coated with 20 µl of Matrigel (10mg/ml). 2% Matrigel complete medium was added to 2 ml. After 16-18 hours cells were fixed with 150 mM Hepes pH 7.5, 4% PFA, 0.05% Glutaraldehyde (5 minutes RT) and 150 mM Hepes pH 7.5, 4% PFA (10 minutes RT - 3 times). Cells were then washed with PBS

1X (5 minutes RT - 3 times), Blocked & Permeabilized in PBS 1x, 1% BSA, 0.1% Saponin (1 hour 30 minutes RT). Primary antibody was then added in PBS 1X, 1% BSA, 0.1% Saponin (4 hours RT). After 3 washes in PBS 1X cells were incubated with secondary antibody (+ Dapi) in PBS 1X, 1% BSA, 0.1% Saponin (2 hours RT) then washed in PBS 1X (3 times) and kept in PBS.

Samples were analyzed with SP8 confocal microscope (20X dry 0.55 NA objective) to identify cells of interests on the gridded coverslips. After selection samples were fixed with of 4% paraformaldehyde and 2,5% glutaraldehyde (EMS, USA) mixture in 0.2 M sodium Cacodylate pH 7.2 for 2h at RT, followed by 6 washes in 0.2 Na Cacodylate pH 7.2 at RT. Then cells were incubated in 1:1 mixture of 2% osmium tetra oxide and 3% potassium ferrocyanide for 1h at RT followed by 6 times rinsing in Cacodylate buffer. Then the samples were sequentially treated with 0.3% Thiocarbohydrazide in 0.2 M Cacodylate buffer for 10 min and 1% OsO4 in 0.2 M Cacodylate buffer (pH 6,9) for 30 min. Samples were rinsed with 0.1 M sodium Cacodylate (pH 6.9) buffer until all traces of the yellow osmium fixative have been removed, washed in de-ionized water, treated with 1% uranyl acetate in water for 1 h and washed in water again [22]. The samples were subsequently subjected to de-hydration in ethanol, and embedded in Epoxy resin at RT and polymerized for at least 72 h in a 60 °C oven.

**Immunolabeling with gold particles.** Samples, analyzed with SP8 confocal microscope (20X dry 0.55 NA objective) to identify cells of interests on the gridded coverslips (as described above), were fixed with a mixture of 4% paraformaldehyde and 0.05% glutaraldehyde in 0.15M Hepes for 5 min at RT and then replaced with 4% paraformaldehyde in 0.15M Hepes for 30 min. Afterwards, the cells were washed 3 times in PBS and incubated with blocking solution for 30 min at RT. Then cells were incubated with primary antibody diluted in blocking solution overnight at 4°C. On the following day, the cells were washed 3 times with PBS and incubated with goat anti-rabbit or anti-mouse Fab' fragments coupled to 1.4nm gold particles (diluted in blocking solution 1:100) for 2h and washed 6 times with PBS. Meanwhile, the activated GoldEnhanceTM-EM was prepared according to the manufacturer's instructions and 100 µl were added into each sample well. The reaction was monitored by a conventional light microscope and was stopped after 5-10 min when the cells had turned “dark enough” by washing several times with PBS. Then cells were fixed with 4% paraformaldehyde and 2,5% glutaraldehyde (EMS, USA) mixture in 0.2 M sodium cacodylate pH 7.2 for 2 h at RT, followed by 6 washes in 0.2 sodium cacodylate pH 7.2 at RT. Then cells were incubated in 1:1 mixture of 2% osmium tetra oxide and 3% potassium ferrocyanide for 1 h at RT followed by 6 times rinsing in cacodylate buffer. Samples were sequentially treated with 0.3% Thiocarbohydrazide in 0.2 M cacodylate buffer for 10 min and 1% $OsO_4$ in 0.2 M cacodylate buffer (pH 6.9) for 30 min, rinsed with 0.1 M sodium cacodylate (pH 6.9) buffer until all traces of the yellow osmium fixative were removed, washed in de-ionized water, treated with 1% uranyl acetate in water for 1 h and washed in water again. The samples were subsequently subjected to dehydration in ethanol, and embedded in Epoxy resin at RT and polymerized for at least 72 h in a 60 °C oven[23].

**Sectioning.** In order to get EM images of the cell of interest, cells were grown in three-dimensional Matrigel (collagen) matrix, which was prepared in such a way to get the minimal thickness of the matrix layer. The cell of interest and in the phase of interest was selected during the analysis of the MatTek and then when the cell was selected and the optical sectioning and the Z-stacking was performed using confocal microscope. During Z-stacking the distance between the bottom and the surface of the cell was estimated. Z-stack images were printed and during trimming of the pyramid and its sharpening these images were constantly used. Embedded samples were then sectioned with diamond knife (Diatome, Switzerland) using Leica EM UC7 ultra microtome. In order to start sectioning exactly when the surface of sectioning reached the cell surface, we used display of the ultramicrotome section counter, which shows the total advance and the total number of sections cut from the moment of CLEAR setting. For the trimming and advance to ROI we used the Diatome diamond trimming blades (trimtool 45, and histo) (Diatome, Switzerland). In order to avoid mistakes we took into consideration the possible "shrinkage" of the matrix during its dehydration and embedding into Epon. Thus, when only 3 microns were left before the beginning of the cell surface we replaced the trimming histo-knife with the Ultra 35 knife (Diatome, Switzerland), and then cut two 200-nm sections and then a small series of 70 nm sections. This series then was interrupted by two 200-nm sections and then the same alternating procedure was repeated several times until the moment when according to our estimation of the deepness of the cell, the cell should be finished. Sections were analyzed with a Tecnai 20 High Voltage EM (FEI, now Thermo Fisher Scientific; The Netherlands) operating at 200 kV. Tilt series were recorded at a magnification of 9,600x, 11,500x, 14,500x, 19,000x or 25,000x using software supplied with the instrument.

**1.3.7.1. Tomography**: Two-step CLEM based on the analysis of tomographic reconstructions acquired under low magnification with consecutive reacquisition of EM tomo box under high (11,500-25,000x) magnification and its re-examination was used exactly as described [23]. Briefly, an ultramicrotome (Leica EM UC7; Leica Microsystems, Vienna) was used to cut 60 nm serial thin sections and 200 nm serial semi-thick sections. Sections were collected onto 1 % Formvar films adhered to slot grids. Both sides of the grids were labelled with fiduciary 10 nm gold (PAG10, CMC, Utrecht, the Netherlands). Tilt-series were collected from the samples from ±65° with 1° increments at 200 kV in Tecnai 20 electron microscopes (FEI, now Thermo Fisher Scientific, Eindhoven, the Netherlands). Tilt series were recorded at a magnification of 9,600x, 11,500x, 14,500x, 19,000x or 25,000x using software supplied with the instrument. The nominal resolution in our tomograms was 4 nm, based upon section thickness, the number of tilts, tilt increments, and tilt angle range. The IMOD package and its newest viewer, 3DMOD 4.0.11, were used to construct individual tomograms and for the assignment of the outer leaflet of organelle membrane contours, CLEM was performed exactly as described[24].

##### 1.4 Immunohistochemistry

Mouse tissues and organs were dissected just after the mice were sacrificed by $CO_2$ asphyxiation or cervical dislocation.

Human tissue samples were obtained from: the University of Palermo, School of Medicine, Istituto di Patologia Generale, Palermo; the European Institute of Oncology (IEO), Milano.

Four-micrometers-thick sections obtained from paraffin embedded tissues were stained with H&E for histomorphological evaluation. For immunostaining, tissue sections were dewaxed and rehydrated. The antigen unmasking technique was performed using Novocastra Epitope Retrieval Solutions, pH6 citrate-based buffer and pH9 EDTA-based buffer in thermostatic bath at 98°C for 30 minutes. After the sections were brought at room temperature, the neutralization of the endogenous peroxidase with 3% H2O2 and protein blocking by a specific protein block were performed. For immunostaining on Human kidney, colon, gastric mucosa, prostate, salivary gland and breast samples, Mouse kidney samples and Zebrafish kidney samples from wild type IRSp53 and IRSp53 -/- phenotypes, the following primary antibodies were used: Mouse anti-human IRSp53 (1,5 hours at room temperature, dilution 1:200 pH6), Rabbit anti-mouse IRSp53 (overnight at 4°C, dilution 1:50 pH9, HPA023310 Sigma), Rabbit anti-zebrafish IRSp53 (overnight at 4°C, dilution 1:30, pH9), Rat anti-mouse Endomucin, (overnight at 4°C, dilution 1:100); Rabbit anti-mouse acetyl-α-tubulin (overnight at 4°C, dilution 1:200, clone D20G3 #5335 Cell Signaling); Rabbit anti-mouse PKC ζ (overnight at 4°C , dilution 1:100 pH9, sc-216 Santa Cruz); Rabbit anti-mouse ZO-1 (1 hour at room temperature, dilution 1:100 pH9, GTX108592 Genetex). The immunostaining was revealed by either a polymer detection method (Novolink Polymer Detection Systems Novocastra Leica Biosystems Newcastle Ltd Product No: RE7280-K) and following specific secondary antibodies: donkey anti-rabbit IgG (H&L) specific secondary antibody 1:500 (Novex by Life Technologies) and goat anti-rat IgG (H&L) specific secondary antibody 1:500 (Novex by Life Technologies) and AEC (3-Amino-9-Ethylcarbazole) substrate was used as chromogen. The slides were counterstained with Harris hematoxylin (Novocastra, Ltd). In double-marker immunofluorescence (IF) staining, tissue sections were incubated with the following primary antibodies: Mouse anti-human IRSp53, (1,5 hours at room temperature, dilution 1:200), Rabbit anti-human ZO-1 (overnight at 4°C, dilution 1:100 pH9, GTX108592 Genetex) and Rabbit anti-human Laminin (overnight at 4°C, dilution 1:25 pH6, ab11575

Abcam). The following secondary antibodies were used: Alexa fluor 568-conjugated goat anti-mouse, Alexa fluor 488-conjugated goat anti-rabbit (Life Technologies). Nuclei were counterstained with DAPI (4',6-diamidin-2-fenilindolo). Sections were analyzed with: 1) Zeiss Axio Scope A1 optical microscope (Zeiss, Germany) and microphotographs were collected using an Axiocam 503 Color digital camera with the ZEN2 imaging software (Zeiss Germany); 2) Olympus BX63 Upright microscope equipped with a motorized stage Black and white camera: Hamamatsu Orca AG (12 bit, 6.45 um pixel size) and Color Camera: Leica DFC450C (36 bit, 3.4 um pixel size); 3) Scan Scope XT device and the Aperio Digital pathology system software (Aperio; Leica).

#### 1.5 Biochemical procedures

##### 1.5.1 Standard Cell lysis for WB analysis

After washing with PBS 1x, cells were lysed in JS buffer directly on the plates using a cell-scraper. About 250 μl of JS buffer/10 cm plates were used. Lysates were incubated on ice for 10 minutes and spun at 13200 rpm for 10 min at 4ºC. The supernatant was transferred into a new eppendorf and protein concentration was measured by the Bradford assay (Biorad), following manufacturer's instructions.

##### 1.5.2 SDS polyacrylamide gel electrophoresis (SDS PAGE)

Pre-casted acrylamide gels (BIORAD or Thermo Fisher Scientific) were employed for resolution of proteins.

##### 1.5.3 Western blot analysis

Desired amounts of proteins were loaded onto 0.75-1.5 mm thick polyacrylamide gels for electrophoresis (Biorad). Proteins were transferred in Western transfer tanks (Biorad) to nitrocellulose (Schleicher and Schuell) in Western Transfer buffer 1x (diluted in 20% methanol) at constant Voltage (100V for 1 hour or 30V overnight). PonceauS colouring was used to reveal roughly the amount of proteins transferred on the filters. Filters were blocked 1 hour (or overnight) in 5% milk or 5% BSA in TBS 0.1% Tween (TBS-T).

After blocking, filters were incubated with the primary antibody, diluted in TBS-T 5% milk, for one hour at room temperature, or overnight at 4˚C, followed by three washes of five minutes each in TBS-T and then incubated with the appropriate peroxidase-conjugated secondary antibody diluted in TBS-T for 1 hour. After the incubation with the secondary antibody, the filter was washed three times in TBS-T and the bound secondary antibody was revealed using the ECL (Enhanced Chemiluminescence) method (Amersham).

##### 1.5.4 Co-immunoprecipitation assay

Lysates prepared in IP buffer (40mM Hepes pH7.5, 150mM NaCl, 10mM MgCl2, 2mM EDTA, 0.3% CHAPS, 10mM NaPyr, 50mM NaF, 10mM NaVan, 2mM PMSF, 1mM DTT, PIC) were incubated in the presence the anti-FLAG M2 affinity gel (SIGMA; Figure 5A), anti-GFP mAb agarose (MBL; Figure 5D) or control antibodies (Ctr) for two cycles of 1 hour each at 4°C with rocking. Immunoprecipitates were washed 3 times in IP buffer. After washing, beads were resuspended in 1:1 volume of 2x SDS-PAGE Sample Buffer, boiled for 10 min at 95°C, centrifuged for 1 minute and then loaded onto polyacrylamide gels.

##### 1.5.5 Overlay assay

Nitrocellulose membranes were incubated in TBST 0.1% Triton X-100 buffer and let do dry. Equal or increasing amounts of recombinant-purified proteins were spotted on membranes and let to dry. Membranes, previously blocked in TBST 0.1% Triton X-100 5% Milk, were then incubated with the recombinant protein of interest, resuspended in TBST 0.1% Triton X-100 5% Milk, 1-2 hours at 4

˚C. After extensive washing in TBST 0.1% Triton X-100, membranes were subjected to western blot analysis with the desired antibodies.

##### 1.5.6 GST-fusion and His-fusion proteins production

All the GST and His fusion proteins used were product in bacteria using *E. coli* BL21 Rosetta (DE3) competent cells transformed with the pGEX6P1 or pTRC-His vector in which the desired construct had been cloned.

###### 1.5.6.1 Bacterial culture

*E. coli* BL21 Rosetta (DE3) cells picked from individual colonies, transformed with the indicated GST-fusion, were used to inoculate 200 mL of LB medium (containing ampicillin at 50 µg/mL) and were grown overnight at 37°C. Between 10 and 100 mL of the overnight culture was diluted in 1 litre of LB and was grown at 37°C (240 rpm shaking) till it reached approximately OD=0.4-0.6.
IPTG (1mM) was then added used to induce the protein production. After the induction cells were pelleted down at 6000 rpm for 15 minutes at 4°C and pellets were used immediately or conserved at -80°C after washing in PBS 1X.

###### 1.5.6.2 Protein production

**GST-fusion protein**

Pellets were suspended in GST-lysis buffer (15mL for 1L culture). Samples were sonicated 3 times for 30 seconds/each on ice and were pelleted down at 13200 rpm for 30 minutes at 4°C using a JA 20 Beckman rotor or at 40000 rpm for 45 minutes at 4°C using a 55.2 Ti Beckman rotor. 1 mL of glutathione-sepharose beads (Amersham), previously washed 3 times with GST-lysis buffer, was added to the supernatant and samples were incubated 1-2 hour at 4°C while rocking. Beads were then washed 3 times (with 5 minutes of incubation at 4°C each) in the GST lysis solution. GST-proteins were resuspendend 50% slurry in the GST-lysis solution. The quantification was achieved in an SDS PAGE gel using a titration curve with BSA.

**GST-lysis buffer**

2x TBS

0.5 mM EDTA

10% Glycerol

protease inhibitor cocktail (Roche, Basel, Switzerland) (freshly added)

1 mM DTT (freshly added)

**His-fusion protein**

Pellets were resuspended in His-lysis buffer (10ml every liter of culture); samples were sonicated 3 times for 30 seconds/each on ice and were pelleted down at 13200 rpm for 30 minutes at 4°C using a JA 20 Beckman rotor or at 40000 rpm for 45 minutes at 4 ˚C using a 55.2 Ti Beckman rotor. 600 µL of NiNTA beads (Qiagen), previously washed 3 times with His-lysis buffer, was added to the supernatant and samples were incubated 1-2 hour at 4 ˚C while rocking. Beads were then washed 2 times in washing buffer 1 and 1 time in washing buffer 2 (5 min, 4 ˚C).

**Elution**

Beads were packed in Poly-Prep® Chromatography Columns (BioRad) and eluted with Elution buffer (500µl fractions). Fractions, evaluated by Bradford assay and SDS-PAGE, were pooled together and buffer exchange with Exchange buffer, using PD10 columns (GE Healtcare). Samples were diluted 1:2 in Dilution buffer and loaded on RESOURCE S cation exchange chromatography column (GE Healtcare) (settings for ResS run: 1CV buffer A, than gradient 0→50%B 60CV, fractions: 1.2ml). Fractions were pulled, concentrated in S200 10/30 column equilibrated with Modified storage buffer, flash frozen and stored at -80 ˚C.

**His-lysis buffer**

50 mM Tris pH8

300 mM NaCl

10 mM imidazole

1mM ß-mercaptoethanol

proteases inhibitors

10% glycerol

**Washing buffer 1**

20mM imidazole

600mM NaCl

50mM Tris pH8

1mM ß-mercaptoethanol

10% glycerol

**Washing buffer 2**

40mM imidazole

300 mM NaCl

50mM tris pH 8

1 mM ß-mercaptoethanol

10% glycerol

**Elution buffer**

200m M imidazole

50 mM Tris pH8

200mM NaCl

1mM ß-mercaptoethanol

10% glycerol

**Exchange buffer**

50mM Tris pH6.8

100mM NaCl

5% glycerol

1mM DTT

0.5mM EDTA

**Dilution buffer**

50mM Tris pH 6.8

5% glycerol

0.5mM EDTA

**ResS buffer A**

50mM NaCl

50 mM Tris pH 6.8

5% glycerol

1mM EDTA

1mM DTT

**ResS buffer B**

1 M NaCl

50 mM Tris pH 6.8

5% glycerol

1mM EDTA

1mM DTT

**Modified storage buffer**

50mM Tris 7.5

200mM NaCl

1mM DTT

10% glycerol

##### 1.5.7 SILAC BioID of IRSp53 interactors

BirA* expressing vector was purchased from Addgene (plasmid # 35700). BirA* was cloned upstream of IRSp53 or GFP, as control, in lentiviral pLVX vector (Clonetech). HeLa cells were infected with lentiviral particles generated from pLVX BirA*-IRSp53 or pLVX-BirA*-GFP. The two populations have been treated with heavy (BirA*-GFP) and light (BirA*-IRSp53) labelled DMEM medium for SILAC (Thermo Fisher Scientific) (supplemented with dialyzed fetal bovine serum) for 6 passages; swap of labelling was performed in the replicate. Cells have been treated with 50 µM biotin for 24 hours and lysed in JS buffer, then biotinylated proteins were fished out with streptavidin-conjugated magnetic beads (Thermo Fisher Scientific). Beads were washed out from aspecific binders using 50 mM Tris pH 7.5, 1% SDS and protease inhibitors; elution was performed using SDS-PAGE Sample buffer 2X, coupled with 95°C boiling. Eluates were then subjected to SDS-PAGE; after colloidal blue staining, bands were excised and reduced with 10 mM dithiothreitol, alkylated with 55 mM iodoacetamide and finally digested overnight with 12.5 ng/µl of trypsin (Roche). After acidification, peptide mixtures were concentrated and desalted, dried in a Speed- Vac and resuspended in 12-15 µL of solvent A (2% acetonitrile, 0.1% formic acid). 5µl of each digested sample from the forward and reverse experiments were loaded on a LC (liquid chromatography)–ESI–MS/MS quadrupole Orbitrap QExactive mass spectrometer (Thermo Fisher Scientific). Peptides were separated on a linear gradient from 95% solvent A to 40% solvent B (80% acetonitrile, 0.1% formic acid) over 30 min and from 40 to 100% solvent B in 3 min at a constant flow rate of 0.25 µl/min on UHPLC Easy-nLC 1000 (Thermo Scientific), where the LC system was connected to a 25-cm fused-silica emitter of 75 µm inner diameter (New Objective, Inc. Woburn, MA, USA), packed in-house with ReproSil-Pur C18-AQ 1.9 µm beads (Dr Maisch Gmbh, Ammerbuch, Germany). MS data were acquired using a data- dependent top 12 method for HCD fragmentation. Survey full scan MS spectra (300– 1650 Th) were acquired in the Orbitrap with 70000 resolution, AGC target 3e6, IT 60 ms. For HCD spectra, resolution was set to 17 500 at m/z 200, AGC target 1e5, IT 120 ms;

Normalized Collision energy 25% and isolation with 2.0 m/z. Technical replicates were conducted on the LC–MS-MS part of the analysis. Raw data were processed with MaxQuant ver. 1.4.1.2. Peptides were identified from the MS/MS spectra searched against the UniProt_Human_2014_10 database using the Andromeda search engine in which trypsin specificity was set up with a maximum of two missed cleavages. Cysteine carbamidomethylation was used as fixed modification, methionine oxidation and protein N-terminal acetylation as variable modifications. Mass deviation for MS/MS peaks was set at 20 ppm. The peptides and protein false discovery rates (FDR) were set to 0.01; the minimal length required for a peptide was six amino acids; a minimum of two peptides and at least one unique peptide was required for high-confidence protein identification. The lists of identified proteins were filtered to eliminate reverse hits and known contaminants. For quantitative analysis, "re-quantify" and "second peptide" options were selected. The statistical program Perseus (ver. 1.5.1.6) was used to quantify significantly up and down regulated proteins following the criteria: (i) significance A with a Benjamini-Hochberg FDR<0.05; (ii) ratio normalized values concordant in both forward and reverse experiments; (iii) minimum H/L ratio counts = 2. Proteomic data are available at the PeptideAtlas repository (http://www.peptideatlas.org/PASS/PASS01464).

##### 1.6 Zebrafish strains

Adult zebrafish were maintained in a multi-rack system (from *Aquatic Habitats*) at a water temperature of 28 °C, pH 7 and conductivity 600 µS. Zebrafish embryos and larvae not older than 5 dpf were maintained at 28.5 °C in E3 water (50 mM NaCl, 0.17 mM KCl, 0.33 mM CaCl, 0.33 mM MgSO4, 0.05% methylene blue). Zebrafish strains used in this study are *AB* (referred to as wild type) *sa11319* obtained from European Zebrafish Resource Center (EZIRC) referred to as *baiap2a*$^{C201*}$ and *Tg*(*cldnB:GFP)*. All the strains were maintained and bred according to the national guidelines (Italian decree "4 March 2014, n.26"). All experimental procedures were approved by the FIRC Institute of Molecular Oncology Institutional Animal Care and Use Committee and Italian Ministry of Health.

##### 1.6.1 Genomic DNA extraction from zebrafish embryos

Caudal fin biopsies (fin clip) were incubated 10 min. at at 98 °C in 50 µl of lysis buffer (Tris-HCl 10 mM pH 8.0, EDTA 1 mM, 0.3% Tween, 0.3% NP40) followed by the ice cooling. 5 µl of Proteinase K 10 mg/ml (Sigma-Aldrich) were added and samples were incubated at 55 °C O/N. The second day, 145 µl of sterile water were added, followed by 20 µl of Sodium Acetate and 200 µl of Phenol. Samples were mixed by inverting them and centrifuged at 13000 rpm for 1 minute. Supernatant was collected and precipitated O/N with 100% ethanol at −20 °C. The third day, samples were centrifuged for 30 minutes at 4 °C and recovered pellets were washed with 75% ethanol, centrifuged again for 5 minutes and resuspended in 20 µl of DNAse-free water.

***sa11319* genotyping**

Genomic DNA (gDNA) was extracted from the caudal fin biopsies of adult *sa13359* fish (generated with ENU at Sanger Institute) and a fragment of 420 bp containing *baiap2a* mutation was amplified by PCR with the specific primers forward 5'-TGTTGAGGCCATCAGCAGTA-3' and reverse 5'-CAAACTGTGCCCAATGGAG-3' and sequenced (Cogentech Sequencing Facility). Only heterozygous fish were maintained and in-crossed to obtain omozygous embryos and adult fish.

##### 1.6.2 RNA extraction from zebrafish, cDNA synthesis and RT-PCR

Wild type zebrafish larvae (AB strain) were collected at 2, 6, 24 and 48 hpf and RNA was extracted using TRIZOL Reagent (Invitrogen) and RNAse Mini kit (QIAGEN). To avoid genomic DNA contamination, samples were digested with RQ1 RNase-Free DNase (Promega). The cDNA was retrotranscribed from 1 µg of RNA using SuperScript VILO cDNA Synthesis kit (Invitrogen), according to manufacturer instructions. 500 ng of cDNA were used as template for a semi quantitative RT-PCR using the following primers: 5'-ACGGAGTGTCTCAGGGAAGA-3' and 5'-

TTCTCTCCATAGTGCCAGCC-3’ for *baiap2a*, 5’-TTGGAGAGAAATGGAC-3’ and 5’-CGTGTAGGAGAACGGGAACCA-3’ for *baiap2b*. β-actin was used as housekeeping.

##### 1.6.3 Hematoxylin and eosin staining and immunostaining on paraffin sections

Larvae were fixed O/N at 4 °C in 4% PFA diluted in PBS and positioned in a 7 × 7 × 6 mm plastic base-molds (Kaltek) containing 1.2% low-melting agarose in PBS. Before agarose solidification, larvae were correctly oriented. After agarose block solidification, larvae were removed from the base mold and immersed in 70% ethanol. After dehydration, agarose blocks were subjected to paraffin embedding by Leica ASP300 S Fully Enclosed Tissue Processor and 5 μm thick sections were cut using a manual rotatory microtome (Leica). Sections were stained with Harris hematoxilin solution for 2 minutes, washed in running water for 5 minutes, counterstained with Eosin-Y solution for 7 seconds and washed in running tap water for 5 minutes. Sections were dehydrated with 95% ethanol and 100% ethanol for 5 minutes two times. Then, they were cleared two times with xylene for 5 minutes and mounted on a glass slide. Sections were finally imaged using a Nikon Eclipse 90i microscope, respectively with 20X and 100X objectives. For immunostaining, sections were incubated in sodium citrate buffer (2.94 mg/ml tri-sodium citrate pH 6, 0.05% Tween 20) at 95 °C for 45 minutes and cooled at RT for 1 hour under chemical hood. Sections were then incubated in blocking solution (2% fetal bovine serum, 2 g bovine serum albumin, 0.05% Tween 20 in PBS 1X adjusted at 7.2 pH) for 1 hour at RT followed by IRSp53 primary antibody diluted in blocking solution O/N. Samples were rinsed in PBS 1X three times for 5 minutes. The antibody signal was revealed with DAB and haematoxylin staining was performed, then sections were dehydrated and mounted with Eukitt-mounting medium.

##### 1.6.4 Zebrafish whole-mount IF and sections

3 days post fertilization larvae were fixed O/N at 4 °C with 4% PFA diluted in PBS 1X and rinsed 3 times with PBS 1X. Embryos older than 24 hpf were treated with 0.25% trypsin (Sigma-Aldrich) at

RT for a range of time between 2 minutes (for 24 hpf embryos) up to 60 minute (for 5 dpf larvae). Samples were then rinsed 3 times for 5 minutes with washing buffer (1% Triton-X100, 0.2% DMSO in PBS 1X) and incubated for at least 1 hour in blocking buffer (0.1% Triton X-100, 1% DMSO, 5% normal goat serum in PBS 1X) on a shaker. Subsequently, embryos were incubated with primary antibodies diluted in blocking buffer O/N at 4 °C. The following day, samples were rinsed rapidly twice with washing buffer and at least 3 washes of 1–2 hours each with washing buffer were performed. Samples were incubated in blocking buffer for 30 minutes followed by secondary antibodies diluted in blocking buffer O/N at 4 °C. The final day, samples were rapidly rinsed 2 times with washing buffer and two washes of 5 minutes each with PBS 1X were performed. Samples were incubated 10 minutes with DAPI, rapidly rinsed with PBS 1X and mounted on a glass slide in 85% glycerol. The following primary antibodies were used: chicken anti-GFP 1:1000 (Abcam) and H3S10ph 1:800 (Upstate). Alexa fluor-488 and -543 1:400 (Invitrogen) were used as secondary antibodies. Pronephric ducts were first acquired using a Leica SP8 confocal microscope equipped with 63X immersion oil objective, deconvolved with a Huygens software (Fig. 8A and Supplementary Movie S2) and then elaborated (Supplementary Movie S2) with a 3D rendering processing (Scientific Volume Imaging).

**Vibratome section**

To prepare 80 μm transversal sections, 3 dpf embryos previously stained were cutted in PBS with a vibratome after inclusion in 5% low-melting agarose. Sections obtained were equilibrated and mounted in glycerol 85% in PBS on glass slides and observed under a Leica TC-SP2 confocal microscope.

**Paraffin section**

3 dpf embryos were fixed, included and processed as already described. 4μm sections were dewaxed and rehydrated. The antigen unmasking technique was performed using Novocastra Epitope Retrieval

Solutions, pH6 citrate-based buffer in thermostatic bath at 98°C for 30 minutes. After the sections were brought at room temperature and stained with chicken anti-GFP 1:1000 (Abcam) and anti-acetyl-α-tubulin 1:500 (Abcam). Alexa fluor-488 and -561 1:400 (Invitrogen) were used as secondary antibodies and Dapi to counterstain nuclei. Images were acquired using a Leica SP8 confocal microscope equipped with 63X immersion oil objective.

##### 1.6.5 Morpholino injections

Zebrafish embryos were microinjected at 1 cell stage with an Olympus SZX9 and a Picospritzer III microinjector (Parker Instrumentation). Injection mixes were composed of Danieau solution 1X (NaCl 58 mM, KCl 0.7 mM, $MgSO_4$ 0.4 mM, $Ca(NO_3)_2$ 0.6 mM, HEPES 5.0 mM, pH 7.6), Phenol red 0.1% and Morpholino (MO) antisense oligos (Gene Tools). Each embryo was injected respectively with 1.7 ng of splice-blocking *baiap2b* MO, 5'-TTCGGGCACTACATGAGTGACCTT-3', and 1.7 ng of 5'-UTR *baiap2b* MO, 5'-AAAGGTCACTCATGTAGTCGCCGAA-3', together.

##### 1.6.6 Western blot analysis

Larvae were lysed in sample buffer; 40 µg of total extracts were resolved by SDS-PAGE, transferred to nitrocellulose and tested with the IRSp53 (1:50) antibody. Band intensities were quantified using ImageJ.

#### 1.7 Microscopes

**Widefield**

Upright Olympus BX51 FL, equipped with 60× UPlanApo 1.35 NA oil and a Photometrics Cool SnapEZ camera.

Software: Metamorph

**Confocal**

**Spinning disk**

UltraVIEW VoX (Perkin Elmer) spinning disk confocal unit, equipped with an EclipseTi inverted microscope (Nikon), a C9100-50 EMCCD camera (Hamamatsu) and driven by Volocity software (Improvision, Perkin Elmer). Environmental microscope incubator (OKOLab) set to 37°C and 5% $CO_2$ perfusion. Nikon PLANApo VC 1.4 NA60X oil immersion objective.

**SP5inv**

Leica TCS SP5 laser confocal scanner mounted on a Leica DMI 6000B inverted microscope equipped with motorized stage, HC PL Fluotar 10X/0.30NA dry objective, HC PL FLUOTAR 20X/0.5NA dry objective, HCX PL APO 40X/1.25-0.75NA oil immersion objective and HCX PL APO 63X/1.4NA oil immersion objective.

Laser lines available:

405 nm,

Argon Laser (458 nm, 476 nm, 488 nm, 496 nm, 514 nm)

DPSS 561 nm

HeNe 633 nm

Software: Leica LAS AF.

**SP2 AOBS**

Leica TCS SP2 AOBS laser confocal scanner mounted on a Leica DM IRE2 inverted microscope.

Objective: HCX PL APO 63X/1.4NA oil immersion objective and HCX PL APO 40X/1.25-0.75NA oil immersion objective.

Laser lines: Violet (405nm laser diode), blue (488nm argon laser), yellow

(561 nm laser diode) and red (633 nm laser diode).

Software: Leica Confocal Software (LCS).

**SP8 white laser**

Leica TCS SP8 laser confocal scanner mounted on a Leica DMi 8 inverted microscope equipped with motorized stage. Objectives: HC PL APO CS2 20X/0,75 dry objective, HC PL APO CS2 40X/1,30 oil immersion immersion objective and HC PL APO CS2 63X/1,40 oil immersion objective.

Laser Lines Available**:**

405 nm pulsed

Argon Laser (458 nm, 476 nm, 488 nm, 496 nm, 514nm)

White light laser tunable in the range: 470 nm-670 nm

Software: Leica Application Suite X (LASX) ver. 3.5.2.18963

**SP8**

Leica TCS SP8 laser confocal scanner mounted on a Leica DMI 8 inverted microscope equipped with motorized stage. Objectives: HC PL FLUOTAR 20X/0,55 dry objective, HC FLUOTAR 25X/0,95 water objective, HCX PL APO 40X/0,75-1,25 oil immersion objective, HC PL APO CS2 63X/1,40 oil immersion objective

Lasers:

405nm diode.

Argon (458 nm , 476 nm ,488 nm ,496 nm ,514 nm).

DPSS 561 nm.

HeNe 633 nm.

Incubation system: Okolab bold line.

Software: Leica Application Suite X, ver. 3.5.1.18803

**Statistical analysis**

All data are presented as the mean ± SD, with the exception of Figure 7B where the mean ± s.e.m is reported. The number of experiments as well as the number of samples analyzed is specified for each experiment and reported in the figure legends. A two-tails, Student's t-test with Welch corrections for two samples with un-equal variance was used to calculate the P values, with the exception of Fig 4A, *bottom graph*, and Fig. 4B, *bottom graph*, where a one-tail student t-test was used. * $p < 0.05$; ** $p < 0.01$, *** $p < 0.001$.

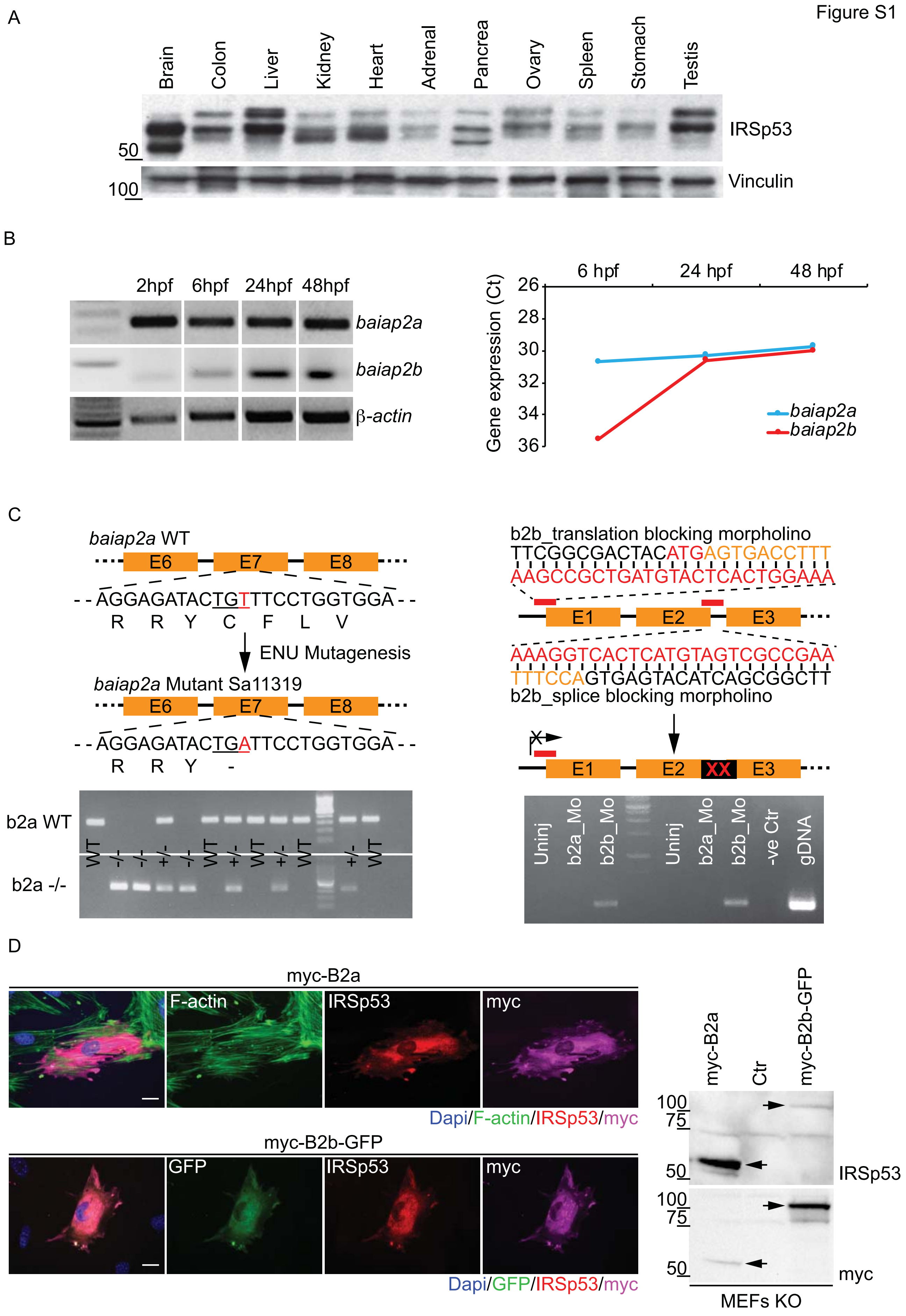
Figure S1
A
Brain
Colon
Liver
Kidney
Heart
Adrenal
Pancrea
Ovary
Spleen
Stomach
Testis
IRSp53
Vinculin
50
100
B
2hpf
6hpf
24hpf
48hpf
baiap2a
baiap2b
β-actin
6 hpf
24 hpf
48 hpf
Gene expression (Ct)
26
28
30
32
34
36
baiap2a
baiap2b
C
baiap2a WT
E6
E7
E8
- - AGGAGATACTGTTTCCTGGTGGA - -
R R Y C F L V
ENU Mutagenesis
baiap2a Mutant Sa11319
- - AGGAGATACTGAATTCCTGGTGGA - -
R R Y -
b2a WT
b2a -/-
WT
-/-
+/-
b2b_translation blocking morpholino
TTCGGCGACTACATGAGTGACCTTT
AAGCCGCTGATGTACTCACTGGAAA
E1
E2
E3
AAAGGTCACTCATGTAGTCGCCGAA
TTTCCAGTGAGTACATCAGCGGCTT
b2b_splice blocking morpholino
XX
Uninj
b2a_Mo
b2b_Mo
-ve Ctr
gDNA
D
myc-B2a
F-actin
IRSp53
myc
Dapi/F-actin/IRSp53/myc
myc-B2b-GFP
GFP
Dapi/GFP/IRSp53/myc
Ctr
100
75
50
myc
MEFs KO

**Figure S1. Characterization of the expression of IRSp53 in murine tissues and of orthologues Baiap2A and 2B in zebrafish**

A) IRSp53 is expressed in different mouse organs. Lysates derived from the indicated organs were processed by immunoblotting to analyse the expression levels of IRSp53 and vinculin.

B) *baiap2a* and *baiap2b* expression analysis during zebrafish embryogenesis. Total RNA was extracted from embryos at different developmental stages and retro-transcribed. The same amount of cDNA (500 ng) was used as template for Reverse Transcriptase-PCR (RT-PCR). *Left*: Semi-quantitative RT-PCR. *Right:* qRT-PCR analysis, data are expressed as Ct after normalization with the house keeping gene (β-actin).

C) Schematic of the genomic regions of the zebrafish genes *baiap2a* (on the left) and *baiap2b* (on the right) targeted by ENU mutagenesis or morpholinos. The mutation in *baiap2a* generates a premature stop codon in exon 7. The *baiap2b* translation-blocking morpholino recognizes a region in the 5'-UTR that causes a translation block; the *baiap2b* splice-blocking morpholino was designed on the junction between exon 2 and intron 2 (E2I2) of *baiap2b* and causes the retention of an intronic fragment in the mature mRNA. Below, the gel electrophoresis related to the genotyping of *baiap2a* mutant embryos (left) and to the specificity of the splice-blocking morpholino-*baiap2b* (right).

D) The rabbit polyclonal anti-IRSp53 antibody recognises the gene products of *baiap2a* and *baiap2b*. *Left:* Mouse embryonic fibroblasts derived from IRSp53-KO mice (MEFs KO) were transfected with either myc-Baiap2a (upper panels) or myc-Baiap2b-GFP (lower panels). Cells were fixed and stained as indicated. Scale bar, 25 µm. *Right*: Lysates from control MEFs KO (Ctr) and MEFs KO transiently expressing either myc-Baiap2a or myc-Baiap2b-GFP were immunoblotted with the indicated antibodies. Arrows indicate myc-Baiap2a and myc-Baiap2b-GFP respectively.

Figure S2

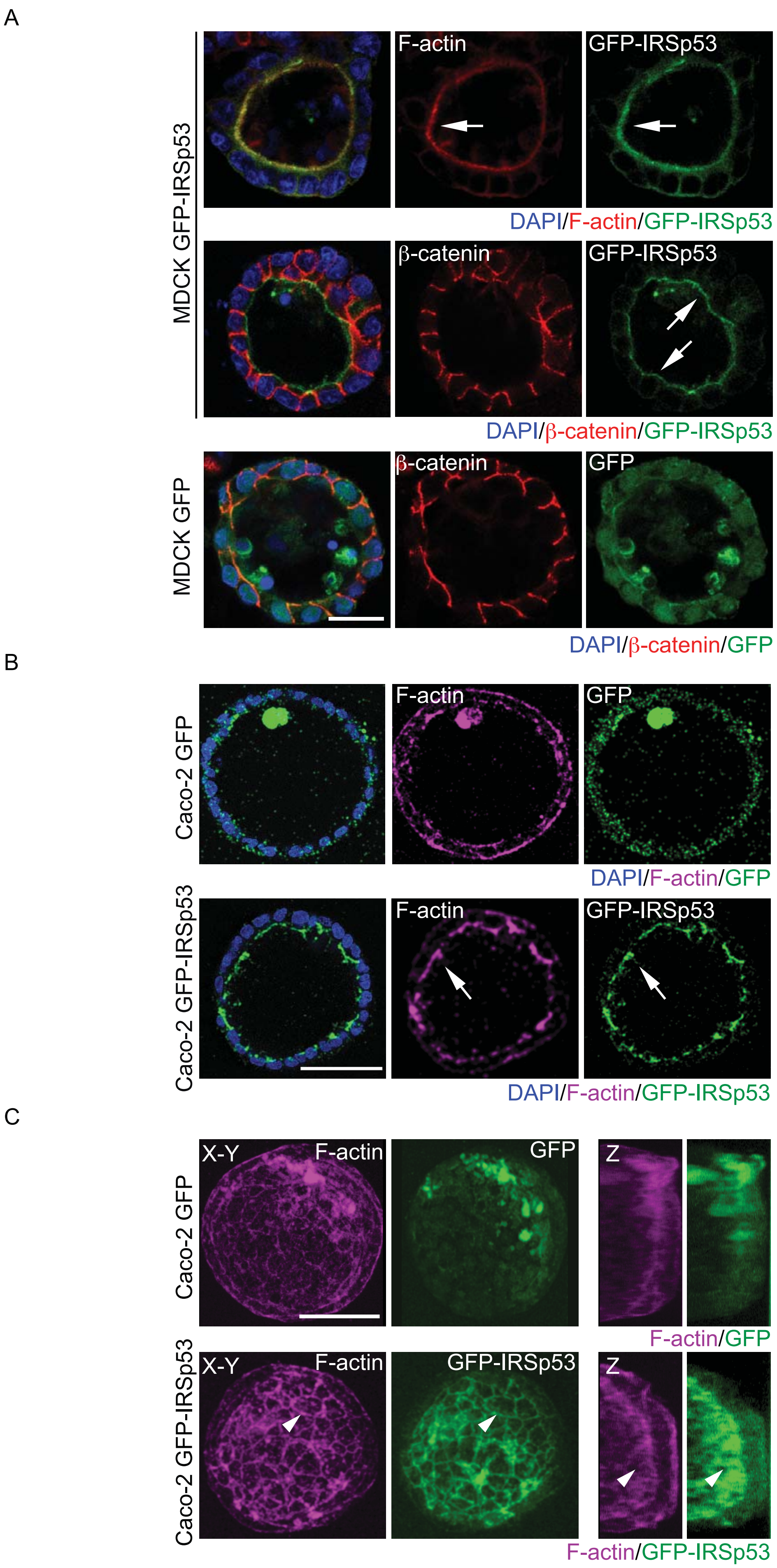
A
MDCK GFP-IRSp53
F-actin
GFP-IRSp53
DAPI/F-actin/GFP-IRSp53
β-catenin
GFP-IRSp53
DAPI/β-catenin/GFP-IRSp53
MDCK GFP
β-catenin
GFP
DAPI/β-catenin/GFP
B
Caco-2 GFP
F-actin
GFP
DAPI/F-actin/GFP
Caco-2 GFP-IRSp53
F-actin
GFP-IRSp53
DAPI/F-actin/GFP-IRSp53
C
Caco-2 GFP
X-Y
F-actin
GFP
Z
F-actin/GFP
Caco-2 GFP-IRSp53
X-Y
F-actin
GFP-IRSp53
Z
F-actin/GFP-IRSp53

**Figure S2. IRSp53 localization in 2D epithelial cells and 3D cysts**

A) MDCK-expressing GFP or GFP-IRSp53 were seeded as single cells on Matrigel and left growing to form 3D cysts. Cysts were fixed and processed for epifluorescence to visualize GFP or GFP-IRSp53 (green) and stained with rhodamine phalloidin (red) to visualized F-actin or anti-β-catenin (red) and Dapi (blue). Arrows indicate IRSp53 enrichment on the luminal side of the cyst. Scale bar, 18 µm.

B) Caco-2 cells, expressing either GFP or GFP-IRSp53, were embedded as single cells into Matrigel/Collagen matrix and left growing to form 3D cysts. Cysts were fixed and processed for epifluorescence to visualize GFP or GFP-IRSp53 (green), and stained with SiR-actin to detect F-actin (magenta) and Dapi (blue). Arrows indicate IRSp53 and F-actin enrichment at the luminal side of the cyst. Scale bar, 70 µm.

C) Maximum projection (X-Y plane, left panels) and 3D reconstruction (Z plane, right panels) of confocal Z-stack acquisitions of the cysts shown in D). Arrowheads indicate IRSp53 and F-actin enrichment at cell-cell junction (X-Y plane) and at the luminal side of the cyst (Z plane). Scale bar, 70 µm.

Figure S3

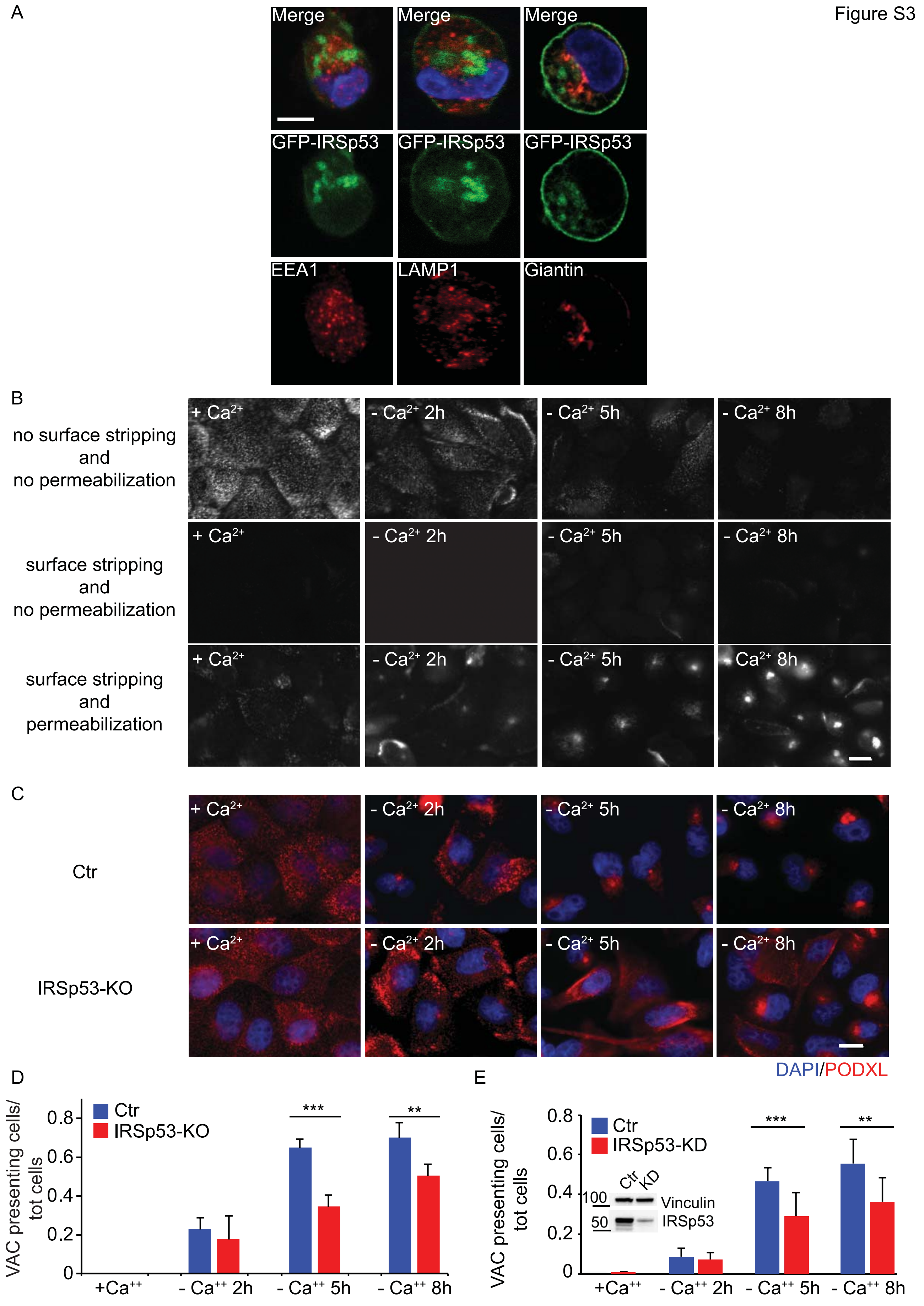
A
Merge
Merge
Merge
GFP-IRSp53
GFP-IRSp53
GFP-IRSp53
EEA1
LAMP1
Giantin
B
no surface stripping and no permeabilization
surface stripping and no permeabilization
surface stripping and permeabilization
+ Ca2+
- Ca2+ 2h
- Ca2+ 5h
- Ca2+ 8h
C
Ctr
IRSp53-KO
DAPI/PODXL
D
VAC presenting cells/ tot cells
Ctr
IRSp53-KO
0.8
0.6
0.4
0.2
0
***
**
+Ca++
- Ca++ 2h
- Ca++ 5h
- Ca++ 8h
E
VAC presenting cells/ tot cells
Ctr
IRSp53-KD
Ctr
KD
100
50
Vinculin
IRSp53
***
**

**Figure S3. IRSp53 impacts on Podocalyxin trafficking to the apical compartment**

A) MDCK expressing GFP-IRSp53 were seeded as single cells on Matrigel and fixed after 8h. Cysts were processed for epifluorescence to visualize GFP-IRSp53 (green) and stained with either anti-EEA1 (red), anti-LAMP1 (red) or anti-Giantin (red) and Dapi (blue). Scale bar, 10 µm.

B) PODXL re-localization assays at VAC (Vacuolar Apical Compartment)[20,21]. MDCK monolayers were grown without $Ca^{2+}$ for the indicated time points and fixed without surface stripping and permeabilization, or subjected to surface stripping and either permeabilized or not, as indicated. Cells were stained with anti-PODXL antibody. Scale bar, 10 µm.

C) IRSp53 depletion delays PODXL re-localization at VAC. MDCK control (Control) or IRSp53-KO monolayers were grown without $Ca^{2+}$ for the indicated time points, subjected to surface stripping and permeabilized after fixation. Cells were stained with anti-PODXL antibody (red) and DAPI (blue). Scale bar, 10 µm.

D) Quantification of PODXL re-localization at VAC, expressed as VAC-presenting cells/ total number of cells. Data are expressed as mean ± SD. At least 100 cells/time point/four different fields were analysed in the experiment. $^{**}p < 0.01$; $^{***}p<0.001$.

E) Quantification of PODXL re-localization at VAC in MDCK Ctr vs IRSp53-KD monolayer (not shown) expressed as VAC presenting cells/ total number of cells. Data are expressed as mean ± SD. At least 100 cells/time point/four different fields were analysed in two independent experiments. $^{**}p < 0.01$; $^{***}p<0.001$. The level of expression of IRSp53 and vinculin were analysed by immunoblotting to verify IRSp53 downregulation.

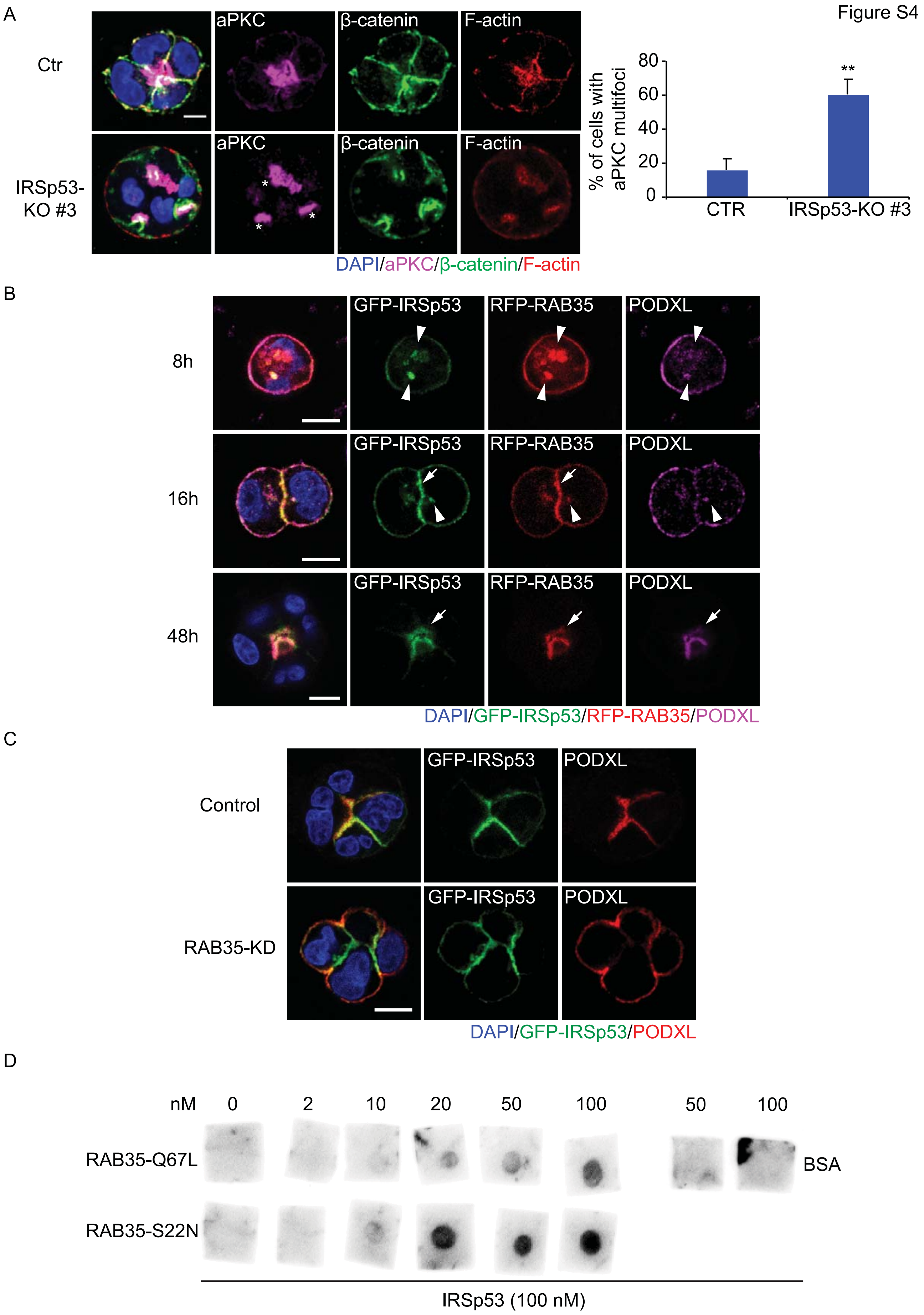
Figure S4
A
Ctr
IRSp53-KO #3
aPKC
β-catenin
F-actin
DAPI/aPKC/β-catenin/F-actin
% of cells with aPKC multifoci
0
20
40
60
80
**
CTR
IRSp53-KO #3
B
8h
16h
48h
GFP-IRSp53
RFP-RAB35
PODXL
DAPI/GFP-IRSp53/RFP-RAB35/PODXL
C
Control
RAB35-KD
GFP-IRSp53
PODXL
DAPI/GFP-IRSp53/PODXL
D
nM
0
2
10
20
50
100
50
100
RAB35-Q67L
RAB35-S22N
BSA
IRSp53 (100 nM)

**Figure S4. IRSp53 controls aPKC distribution, colocalizes and interacts directly with RAB35**

A) Caco-2 control (Ctr) or IRSp53-KO (#3), were embedded as single cells in Matrigel/Collagen matrix and left growing for 48/72h. Cysts were fixed and stained with anti aPKC (magenta), anti-β-catenin (green), rhodamine-phalloidin to detect F-actin (red) and Dapi (blue). Asterisks indicate aPKC multi-foci in IRSp53-KO cyst. Scale bar, 10 μm. *Right:* Quantification of cysts with multiple aPKC foci. Data are expressed as mean ± SD. Four-cells stage cysts were analysed. At least 15 cysts/experiment were analysed in three independent experiments. **$p < 0.01$.

B) Time course of IRSp53, RAB35, and PODXL localization in the early phases of cystogenesis. MDCK expressing GFP-IRSp53 and RFP-RAB35 were seeded as single cells on Matrigel. Cysts were fixed at the indicated time points, processed for epifluorescence to visualize GFP-IRSp53 (green) and RFP-RAB35 (red), and stained with anti-PODXL (magenta) and Dapi (blue). Arrowheads indicate IRSp53, RAB35, and PODXL colocalization in vesicle-like structures. Arrows indicate IRSp53 and RAB35 preceding PODXL re-localization on AMIS at early time points and the enrichment of the three proteins on the luminal side at later time points. Scale bar, 10 μm.

C) RAB35 silencing does not alter IRSp53 localization during cystogenesis. GFP-IRSp53 expressing MDCK treated with a scramble siRNA (Ctr) or a siRNA against RAB35 (RAB35-KD) were seeded as single cells on Matrigel and fixed after 24 hours. Cysts were processed for epifluorescence to visualize GFP-IRSp53 (green), stained with anti-PODXL (red) and Dapi (blue). Scale bar, 10 μm.

D) Recombinant purified GST-RAB35Q67L and GST-RAB35S22N proteins were spotted at the indicated concentrations on nitrocellulose filters. BSA was used as negative control. Nitrocellulose membranes were incubated with recombinant purified IRSp53 protein (100 nM) and immunoblotted with an anti-IRSp53 specific antibody.

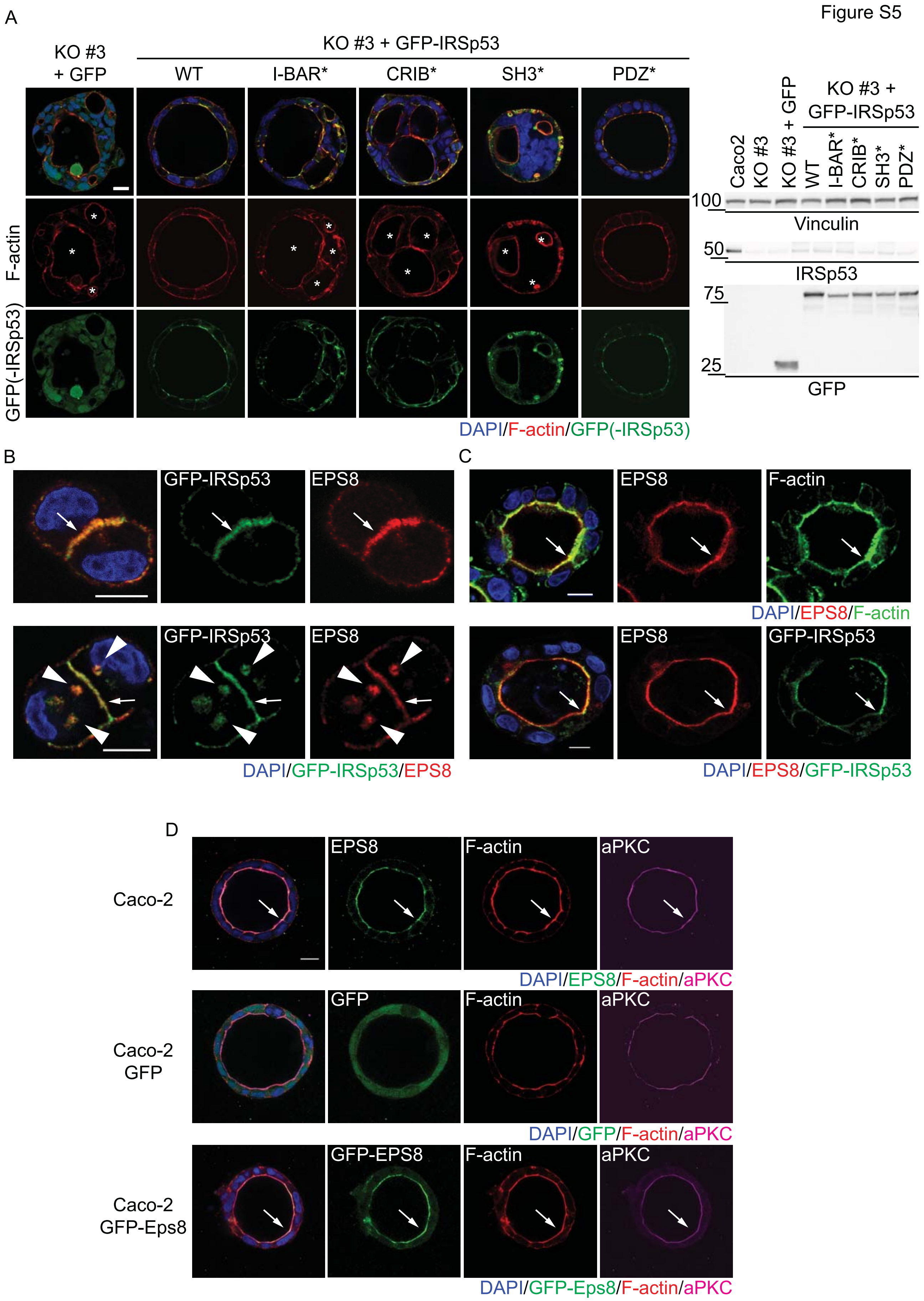
Figure S5
A
KO #3 + GFP
KO #3 + GFP-IRSp53
WT
I-BAR*
CRIB*
SH3*
PDZ*
F-actin
GFP(-IRSp53)
DAPI/F-actin/GFP(-IRSp53)
Caco2
KO #3
KO #3 + GFP
KO #3 + GFP-IRSp53
100
Vinculin
50
IRSp53
75
25
GFP
B
GFP-IRSp53
EPS8
DAPI/GFP-IRSp53/EPS8
C
EPS8
F-actin
DAPI/EPS8/F-actin
GFP-IRSp53
DAPI/EPS8/GFP-IRSp53
D
Caco-2
Caco-2 GFP
Caco-2 GFP-Eps8
aPKC
DAPI/EPS8/F-actin/aPKC
GFP
DAPI/GFP/F-actin/aPKC
GFP-EPS8
DAPI/GFP-Eps8/F-actin/aPKC

**Figure S5. Structure-Function analysis of IRSp53 and Eps8 localization in epithelial cysts**

A) Caco-2 IRSp53-KO #3 stably-infected with GFP or murine GFP-IRSp53 WT, or I-BAR*, CRIB*, SH3*, PDZ* mutants were embedded as single cells in Matrigel/Collagen matrix and fixed after 7 days. Cysts were processed for epifluorescence to visualize GFP or GFP-IRSp53 (green) and stained with rhodamine-phalloidin to detect F-actin (red) and Dapi (blue). Asterisks indicate multiple lumens. Scale bar, 20 μm. *Right:* The expression levels of IRSp53, GFP, GFP-IRSp53 WT and mutants, and vinculin were analysed by immunoblotting to verify the depletion of endogenous IRSp53 and the expression of the GFP-fusion constructs.

B) MDCK infected with murine GFP-IRSp53 were seeded as single cells on Matrigel and fixed after 12 hours. Cells were processed for epifluorescence to visualize GFP-IRSp53 (green) and stained with anti-EPS8 (red) and Dapi (blue). Arrows indicate EPS8 and IRSp53 co-localization at nascent AMIS; arrowheads indicate Eps8 and IRSp53 enrichment in vesicle-like structures. Scale bar, 10 μm.

C) MDCK (upper panels) or MDCK-expressing murine GFP-IRSp53 (lower panels) were seeded as single cells on Matrigel and mature cysts were fixed after 6 days. Cysts were stained with anti-EPS8 (red), FITC-phalloidin to visualize F-actin (green) and Dapi (blue) (upper panels) or processed for epifluorescence to visualize GFP-IRSp53 (green), and stained with anti-EPS8 (red) and Dapi (blue) (lower panels). Arrows indicate EPS8, F-actin and IRSp53 enrichment at the apical/ luminal side of the cysts. Scale bar, 10 μm.

D) Caco-2 (upper panels), Caco-2 infected with GFP (central panels) or Caco-2 infected with GFP-Eps8 were embedded as single cells in Matrigel/Collagen matrix and cysts were fixed after 7 days. Cysts were stained with anti-Eps8 (green), rhodamine-phalloidin to visualize F-actin (red), anti-aPKC (magenta) and Dapi (blue) (upper panels) or processed for epifluorescence to visualize GFP or GFP-EPS8 (green), and stained with rhodamine-phalloidin to visualize F-actin (red), anti-aPKC (magenta) and Dapi (blue) (central and lower panels). Arrows indicate EPS8, F-actin and aPKC enrichment at the apical/luminal side of the cysts. Scale bar, 20 μm.

Figure S6

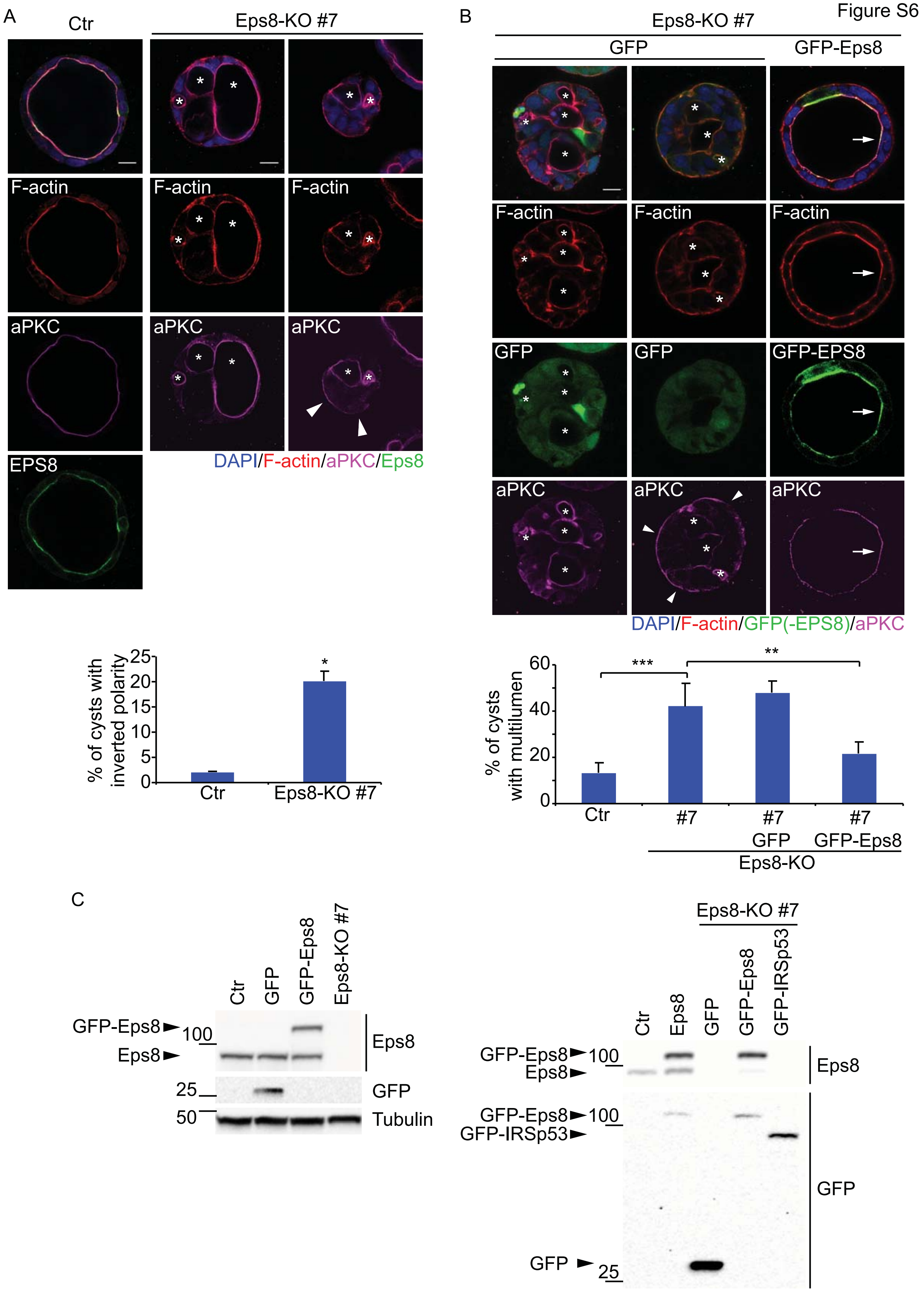
A
Ctr
Eps8-KO #7
F-actin
aPKC
EPS8
DAPI/F-actin/aPKC/Eps8
B
Eps8-KO #7
GFP
GFP-Eps8
GFP-EPS8
DAPI/F-actin/GFP(-EPS8)/aPKC
% of cysts with inverted polarity
25
20
15
10
5
0
*
Ctr
Eps8-KO #7
% of cysts with multilumen
60
40
20
0
***
**
Ctr
#7
#7 GFP
#7 GFP-Eps8
Eps8-KO
C
Ctr
GFP
GFP-Eps8
Eps8-KO #7
GFP-Eps8
100
Eps8
Eps8
25
GFP
50
Tubulin
Eps8-KO #7
Ctr
Eps8
GFP
GFP-Eps8
GFP-IRSp53
GFP-Eps8
100
Eps8
Eps8
GFP-Eps8
100
GFP-IRSp53
GFP
GFP
25

**Figure S6. The loss of Eps8 leads to the formation of Caco-2 cysts with multiple lumens**

A) Caco-2 control (Ctr) or Eps8-KO (clone #7), were embedded as single cells into Matrigel/Collagen matrix and left growing to form cysts. Cysts were fixed and stained with anti-EPS8 (green), rhodamine-phalloidin to detect F-actin (red), anti-a-PKC (magenta), and Dapi (blue). Asterisks and arrowheads indicate multiple lumens and inverted polarity, respectively. Scale bar, 20 µm. *Lower*: Quantification of the number of cysts with inverted polarity. Data are expressed as mean ± SD. At least 15 cysts/ experiment were analysed in two independent experiments. *$p < 0.05$. Quantification of multi-lumen cysts is shown in B (lower graph).

B) Caco-2 Eps8-KO #7, infected with GFP or GFP-Eps8, were embedded as single cells into Matrigel/Collagen matrix and left growing to form cysts. Cysts were fixed, processed for epifluorescence to visualize GFP or GFP-EPS8 (green), and stained with rhodamine-phalloidin to visualize F-actin (red), anti-aPKC (magenta), and Dapi (blue). Asterisks and arrowheads indicate multi-lumens and inverted polarity, respectively. Arrows indicate the enrichment of Eps8, F-actin, and a-PKC at the apical/luminal side of Eps8-KO cysts reconstituted with GFP-EPS8. Scale bar, 20 µm. *Lower:* Quantification of the number of cysts with multiple lumens, in comparison to Caco-2 control cells (C2 Ctr) and Caco-2 EPS8-KO #7. Data are expressed as mean ± SD. At least 15 cysts/experiment were analysed in two independent experiments. **$p < 0.01$; ***$p < 0.001$.

C) The expression levels of GFP, GFP-Eps8, GFP-IRSp53, Eps8, and tubulin were analysed by immunoblotting to detect Eps8 loss in Caco-2 #7 cells and the expression of the GFP-recombinant proteins.

Figure S7

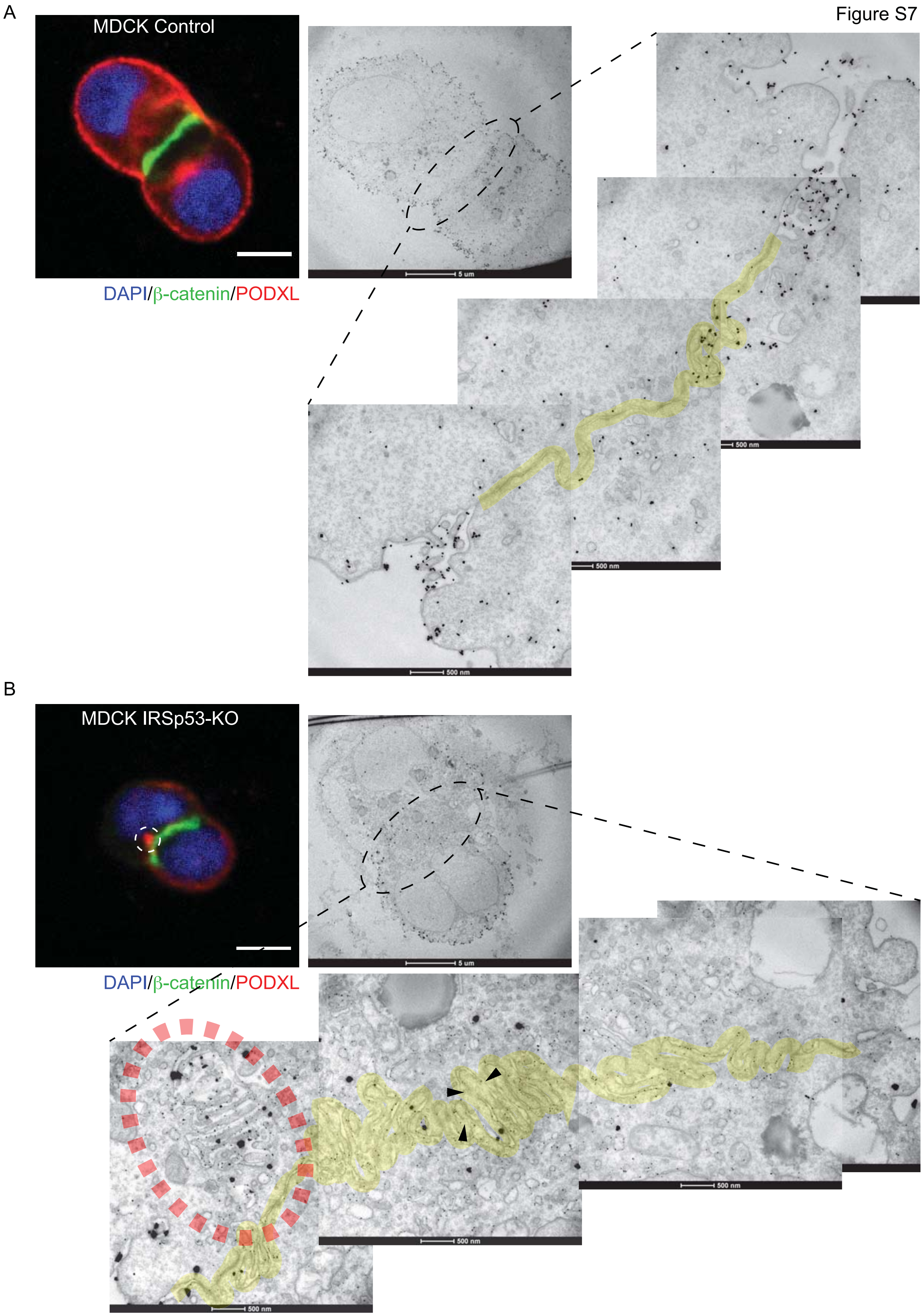
A
MDCK Control
DAPI/β-catenin/PODXL
5 um
500 nm
500 nm
500 nm
B
MDCK IRSp53-KO
DAPI/β-catenin/PODXL
5 um
500 nm
500 nm
500 nm

**Figure S7. TEM analysis of control and IRSp53-KO MDCK at two-cells stage of cystogenesis**

A) Control MDCK cells were seeded, stained with anti-ß-catenin (green), anti-PODXL (red), and Dapi (blue) and analysed by confocal microscope as described in Figure 7B. Two-cell stage cysts were identified on grids by confocal microscopy (left image, scale bar, 10 µm) and processed for EM analysis with immune-gold labelling against PODXL (middle image). Panels on the right are magnified reconstruction of the intervening plasma membranes region. Yellow transparent ribbon highlights the intervening plasma membranes.

B) IRSp53-KO MDCK were treated and analysed exactly as described above. Dashed white (in the IF image) and red (in the EM micrograph) circles highlight the ectopic lumen (or apical vacuole) in IRSp53-KO cells. Arrowheads indicate the inter-cytoplasmic bridges in the PMs. Scale bar of the EM micrographs are 5 µm, and 500 nm in the magnified images.

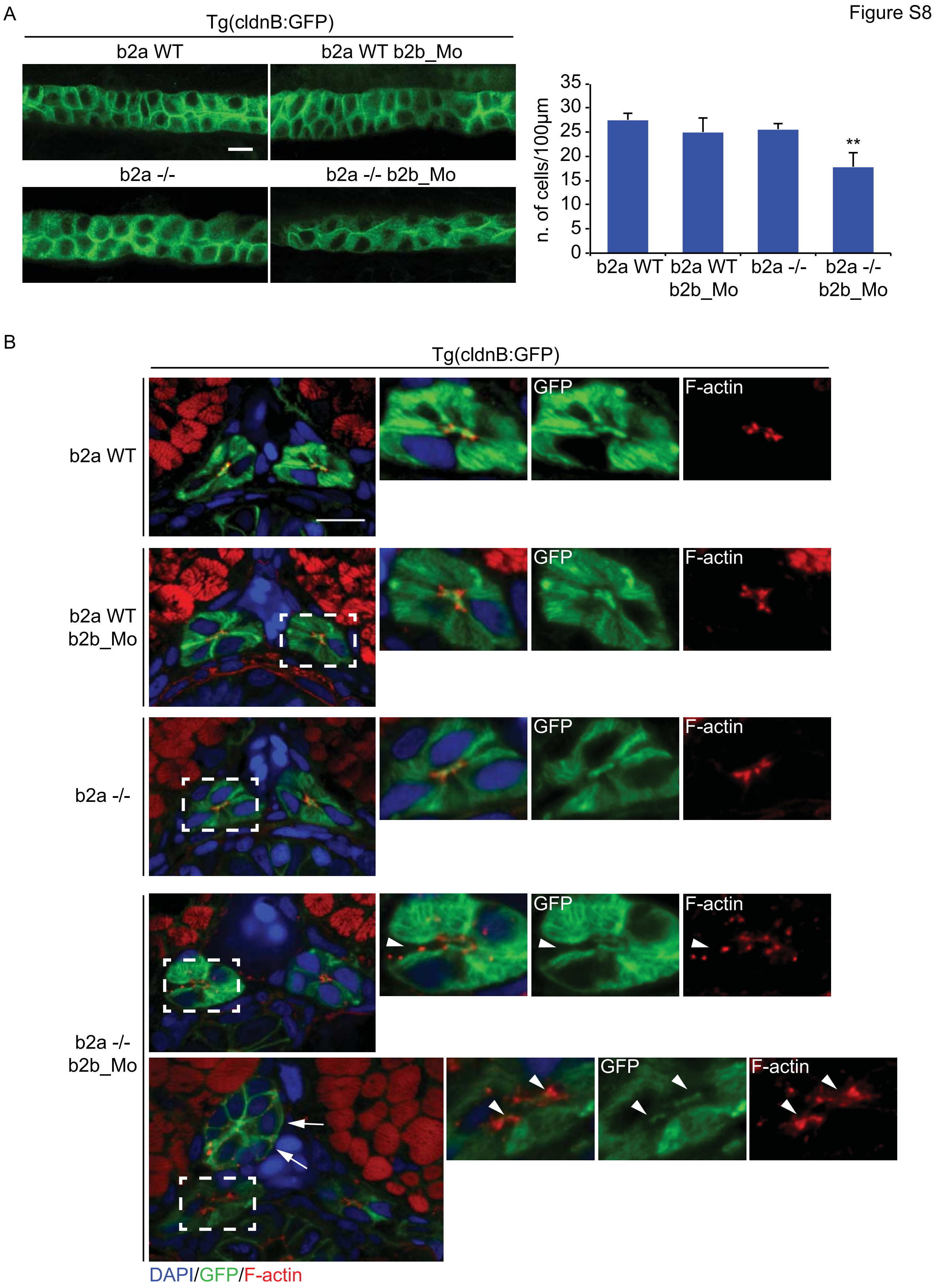
Figure S8
A
Tg(cldnB:GFP)
b2a WT
b2a WT b2b_Mo
b2a -/-
b2a -/- b2b_Mo
n. of cells/100µm
35
30
25
20
15
10
5
0
**
b2a WT
b2a WT b2b_Mo
b2a -/-
b2a -/- b2b_Mo
B
Tg(cldnB:GFP)
b2a WT
b2a WT b2b_Mo
b2a -/-
b2a -/- b2b_Mo
GFP
F-actin
DAPI/GFP/F-actin

**Figure S8. Pronephric duct in developing zebrafish after concomitant loss of *baiap2a* and *2b***

A) Digital light sheet confocal images of 72 hpf pronephric ducts. Embryos obtained by either WT treated with scramble oligo (b2a WT), *baiap2a* mutant (b2a -/-), *baiap2b* morphant (b2a WT b2b_Mo) and *baiap2a* mutant *baiap2b* morphant (b2a -/- b2b_Mo) females, in Tg(CldnB:GFP) genetic background, were fixed and mounted in agarose. Samples were stained with anti-GFP (green). Scale bars, 10 µm. *Right*: The number of nuclei/100µm was counted in different area of the PD in each genetic background. Data are expressed as mean ± SD. At least 4 different area of the PD of 4 embryos in each genetic background were analysed. $^{**}p < 0.01$.

B) Confocal images of 72 hpf pronephric ducts. Embryos obtained by either WT treated with scramble oligo (b2a WT), *baiap2a* mutant (b2a -/-), *baiap2b* morphant (b2a WT b2b_Mo) and *baiap2a* mutant *baiap2b* morphant (b2a -/- b2b_Mo) females, in Tg(CldnB:GFP) genetic background, were fixed and mounted in agarose. Samples were stained with anti-GFP (green), rhodamine-phalloidin to detect F-actin (red) and Dapi (blue). *Right*: 2x magnification of the dashed line areas on the left. Arrows and arrowheads indicate the ectopic-duct structure and the luminal aberrations (open lumen, multi-lumen) respectively in b2a -/- b2b_Mo embryo. ~67% of the b2a -/- b2b_Mo embryos show such defects (n = 8/12). Scale bar, 10 µm.

Figure S9

A

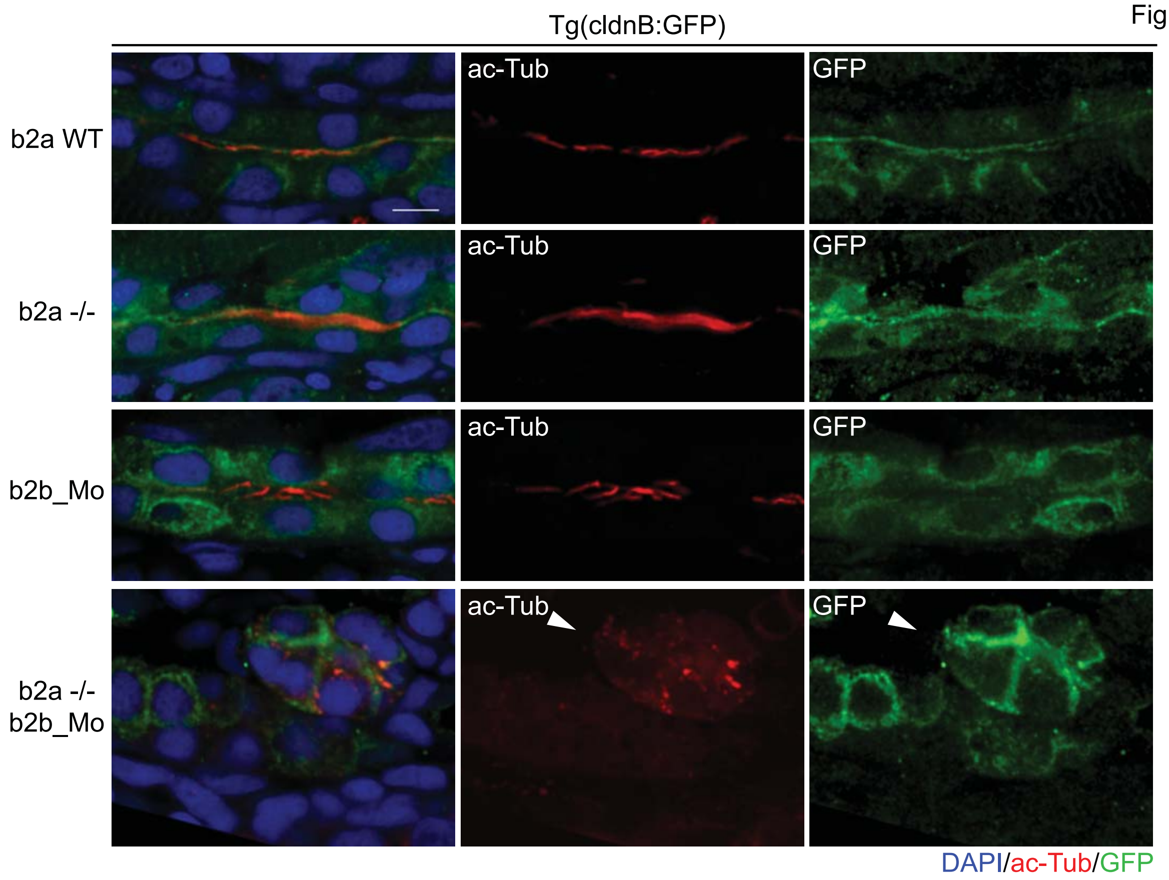


B

| ID | Cortical thinning | Architectural tubular disarray | Tubular anisokaryosis* | Vascular lacunae | Glomerular pseudo-aneurysm | Intratubular cast | Total score | Average |
|---|---|---|---|---|---|---|---|---|
| WT1 M | 0 | 0 | 0 | 1 | 0 | 1 | 2 | 1,75 |
| WT2 M | 0 | 1 | 0 | 1 | 0 | 1 | 3 | |
| WT3 M | 0 | 0 | 0 | 2 | 0 | 0 | 2 | |
| WT4 M | 0 | 1 | 0 | 1 | 0 | 0 | 2 | |
| WT1 F | 0 | 0 | 0 | 0 | 0 | 0 | 0 | |
| WT2 F | 0 | 1 | 0 | 0 | 0 | 0 | 1 | |
| WT3 F | 1 | 0 | 0 | 1 | 0 | 1 | 3 | |
| WT4 F | 0 | 0 | 0 | 0 | 0 | 1 | 1 | |
| KO1 M | 0 | 1 | 0 | 0 | 0 | 0 | 1 | 10 |
| KO2 M | 1 | 2 | 1 | 0 | 0 | 1 | 5 | |
| KO3 M | 1 | 2 | 1 | 0 | 0 | 0 | 4 | |
| KO4 M | 3 | 3 | 2 | 2 | 2 | 2 | 14 | |
| KO5 M | 1 | 2 | 2 | 1 | 2 | 2 | 10 | |
| KO6 M | 1 | 2 | 2 | 1 | 2 | 2 | 10 | |
| KO1 F | 2 | 3 | 2 | 2 | 2 | 2 | 13 | |
| KO2 F | 2 | 3 | 2 | 2 | 3 | 2 | 14 | |
| KO3 F | 3 | 3 | 3 | 3 | 2 | 3 | 17 | |
| KO4 F | 1 | 2 | 2 | 2 | 2 | 2 | 11 | |
| KO5 F | 2 | 3 | 2 | 2 | 0 | 2 | 11 | |
| KO6 F | 3 | 3 | 2 | 0 | 1 | 1 | 10 | |
| KO7 F | 0 | 2 | 1 | 0 | 0 | 2 | 5 | |
| KO8 F | 2 | 3 | 3 | 2 | 2 | 2 | 14 | |
| KO9 F | 1 | 3 | 2 | 3 | 2 | 3 | 14 | |
| KO10 F | 1 | 2 | 2 | 0 | 1 | 1 | 7 | |

0 = absent; 1 = focal presence; 2 = multifocal presence; 3 = diffuse presence
Tubular anisokaryosis*: irregular distribution and nuclear shape in the tubules

**Figure S9. Structural defects of pronephric duct in zebrafish and of adult kidney in WT and IRSp53-KO mice**

A) Confocal analysis of 72 hpf pronephric ducts. Embryos obtained by either WT treated with scramble oligo (b2a WT), *baiap2a* mutant (b2a -/-), *baiap2b* morphant (b2a WT b2b_Mo) and *baiap2a* mutant *baiap2b* morphant (b2a -/- b2b_Mo) females, in Tg(CldnB:GFP) genetic background, were fixed, processed and paraffin embedded. Slices were stained with anti-GFP (green), anti-acetylated-Tubulin (ac-Tub, red) and Dapi (blue). Arrowhead indicates the irregular distribution of acetylated-tubulin in the extra-duct structure of b2a -/- b2b_Mo embryo pronephric duct. Scale bar, 10 μm. B) Detailed analysis of renal defects. Kidneys from adult (4 and 8 months) mice were analysed for the defect described in the table and a score between 0 (absent) and 3 (diffuse presence) was assigned. Total scores represent the sum of the scoring for each of the described defect. The average represents the Renal Damage Score (see also Figure 8B). WT (n = 8; 4M, 4F) and IRSp53-KO (n = 16; 6M, 10F).

Figure S10

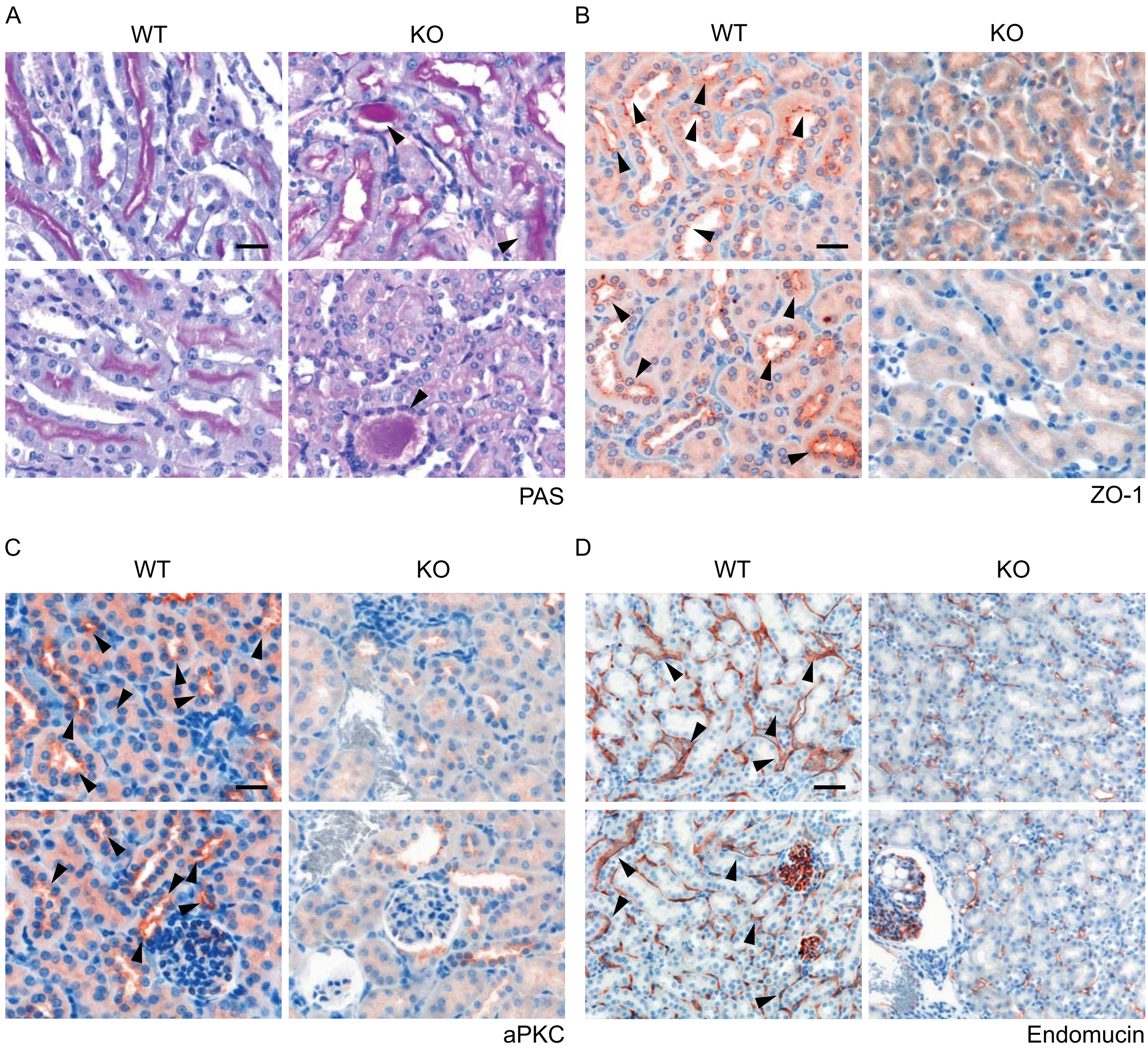

**Figure S10. Genetic removal of IRSp53 leads to polarity defects in mouse kidney.**

A) Periodic acid-Schiff (PAS) analysis in kidney from IRSp53 WT and KO mice. Arrowheads indicate the luminal accumulation of eosinophilic acellular material. Scale bar, 50 μm. B) IHC analysis of ZO-1 expression and localization in kidney from IRSp53 WT and KO mice. Arrowheads indicate ZO-1 apical/luminal enrichment in WT kidney tubules. Scale bar, 50 μm. C) IHC analysis of aPKC expression and localization in kidney from IRSp53 WT and KO mice. Arrowheads indicate aPKC apical/luminal enrichment in WT kidney tubules. Scale bar, 50 μm. D) IHC analysis of endomucin expression and localization in kidney from IRSp53 WT and KO mice. Arrowheads indicate endomucin basal enrichment in WT kidney tubules. Scale bar, 50 μm.

Figure S11

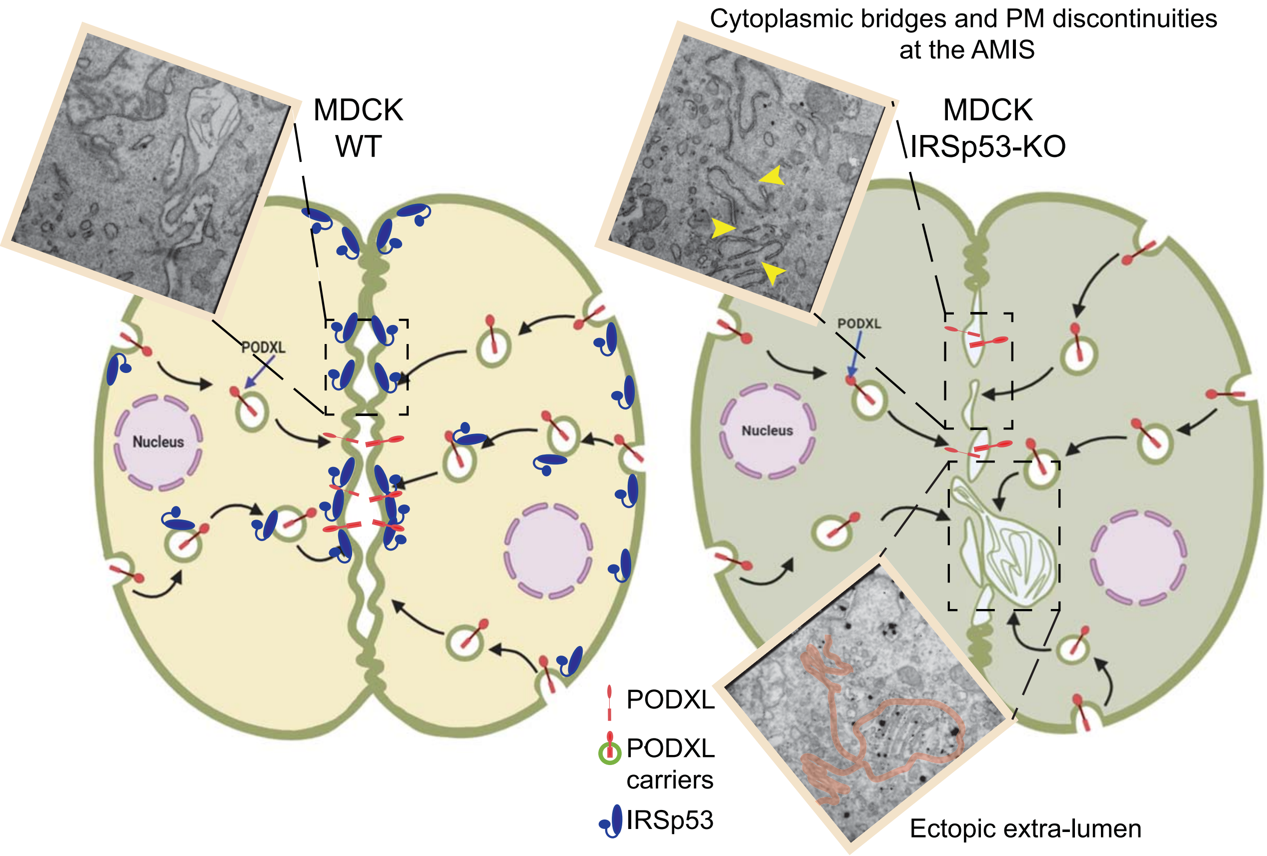
Cytoplasmic bridges and PM discontinuities at the AMIS
MDCK WT
MDCK IRSp53-KO
PODXL
PODXL
Nucleus
Nucleus
PODXL
PODXL carriers
IRSp53
Ectopic extra-lumen

**Figure S11. Shaping the plasma membrane and controlling trafficking of polarized determinants via IRSp53 during the early phase of lumen formation in epithelial cystogenesis.** Control MDCK cells plated on Matrigel in Matrigel-containing media undergo symmetry-breaking events immediately after the first cell division, when after cytokinetic furrow ingression a midbody is formed[25,26]. Around the midbody, the apical membrane initiation site (AMIS) is assembled establishing the location of the nascent lumen. AMIS assembly requires the polarized transport of highly-charged and -glycosylated transmembrane proteins, such as Podocalyxin (PODXL), that from the outer cell face become endocytosed and traffic via vesicular recycling carriers to the newly formed opposing plasma membrane. At this site, repulsive charges contribute to generate interconnected mini-lumens (leftmost inset show a Correlative Light Electron Microscopy-CLEM-analysis of MDCK cells capture after the very first cell division) that will evolve to form a single apical lumen, typical of an epithelial cyst. IRSp53 is required both for directing the trafficking of PODXL and to ensure its targeting at the AMIS via stabilizing the GTPase RAB35 (not shown), and through its interaction with the actin capping protein Eps8 (not shown). At the AMIS, IRSp53 through its membrane deforming and curvature sensing I-BAR domain ensures the structural integrity, continuity and correct shape of the opposing plasma membrane at two-cells stage. Consistent with this, IRSp53 genetic loss leads to interruption of the continuity of the PM at the AMIS (top rightmost inset, yellow arrowhead indicate membrane discontinuities), with the formation of inter-cytoplasmic bridges, and occasionally ectopic lateral lumen (bottom inset, magenta line outlines the intervening PM and the extra lumen).

#### SUPPLEMENTARY MOVIES LEGENDS

**Movie S1. IRSP53 precedes PODXL localization at the apical membrane since the first cytokinesis of 3D cyst development**

A) Time lapse of GFP-IRSp53 and RFP-PODXL during early phases of cystogenesis. MDCK expressing GFP-IRSp53 and RFP-PODXL were seeded as single cells on a Matrigel. 6 hours after seeding cells were subjected to time-lapse analysis using Confocal Spinning Disk microscope. Images (Bright Field, GFP and RFP channel respectively) were acquired every 5 minutes for 15 hours. White arrowheads indicate GFP-IRSp53 accumulation at cleavage furrow immediately after the first cell division; yellow arrowheads indicate enrichment of RFP-PODXL at nascent AMIS at later stages. Scale bar, 10 µm.

**Movie S2. Loss of IRSp53 alters facing PMs at the nascent AMIS**

Electron Microscopy Tomography of MDCK Ctr (left) and IRSp53-KO (right) two-cells stage intervening membranes during early cystogenesis. Cells were seeded and processed as in Figure 7B. Dashed red and black lines define the plasma membrane, while the arrowhead indicates the inter-cytoplasmic bridge of the intervening membranes during AMIS formation at 2-cells stage.

**Movie S3. Genetic removal of IRSp53 leads to defects in *zebrafish* kidney development**

3D rendering of deconvoluted confocal images of pronephric ducts of Tg(CldnB:GFP)-WT, treated with scramble oligo (b2a WT), or Tg(CldnB:GFP)-*baiap2a* mutant zebrafish strain treated with splice-blocking and translational-blocking morpholinos (b2a -/- b2b_Mo).

**ORIGINAL UNCROPPED IMMUNOBLOTS AND GELS FOR ALL FIGURES**

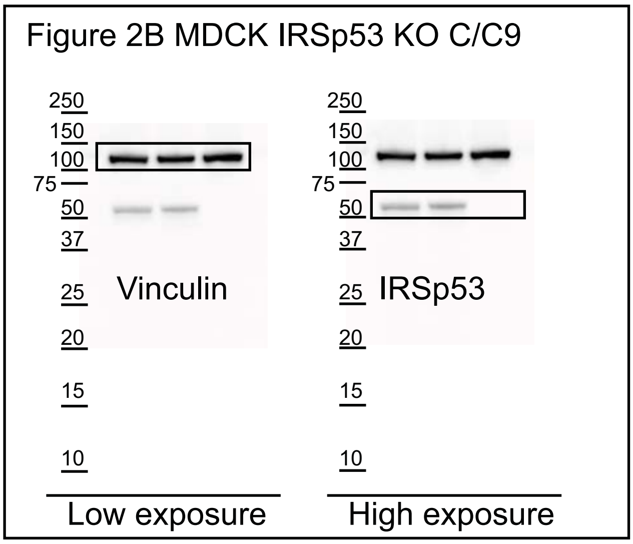


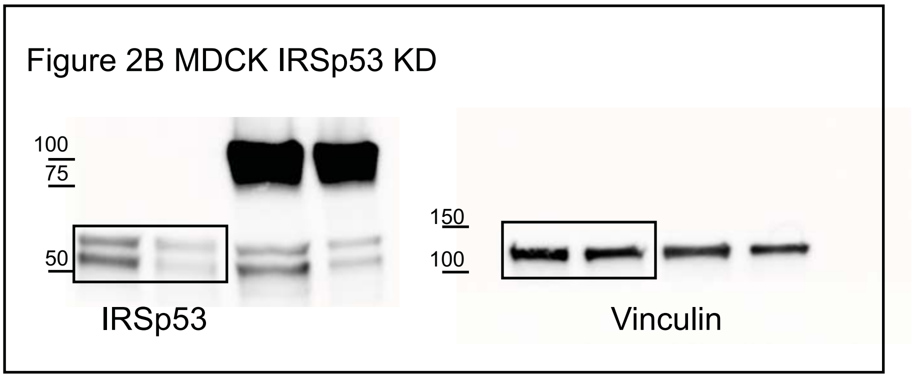


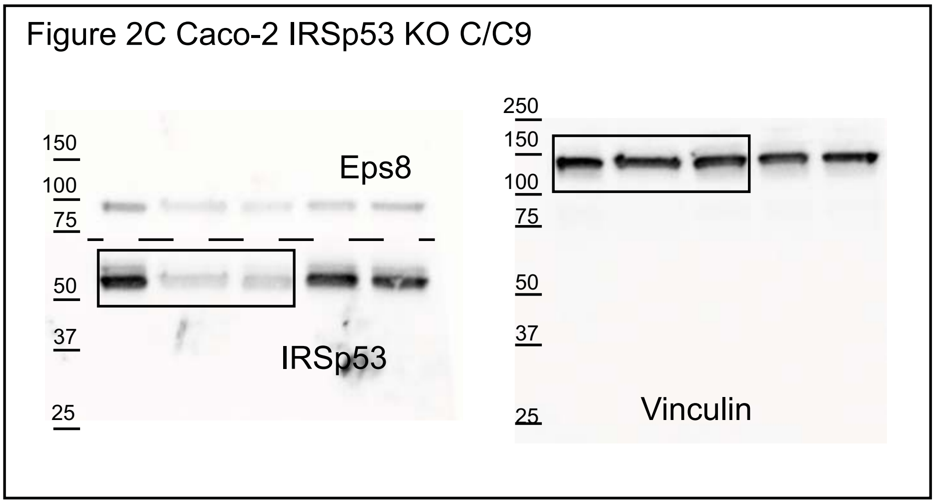

Figure 4B MDCK IRSp53 KD; Rab35 KD

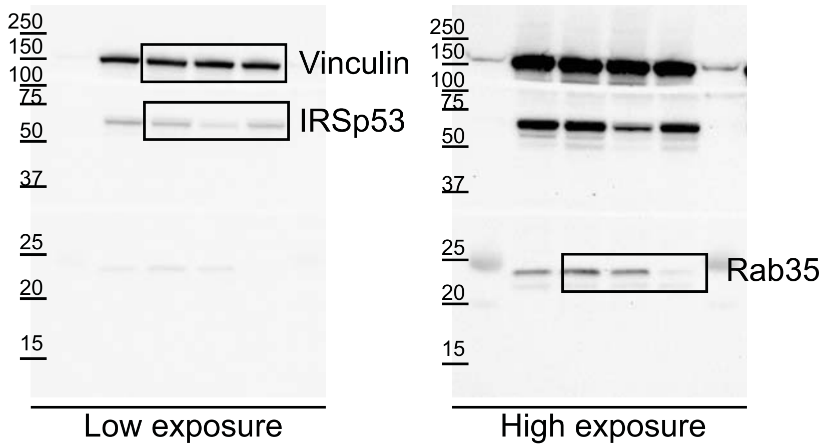

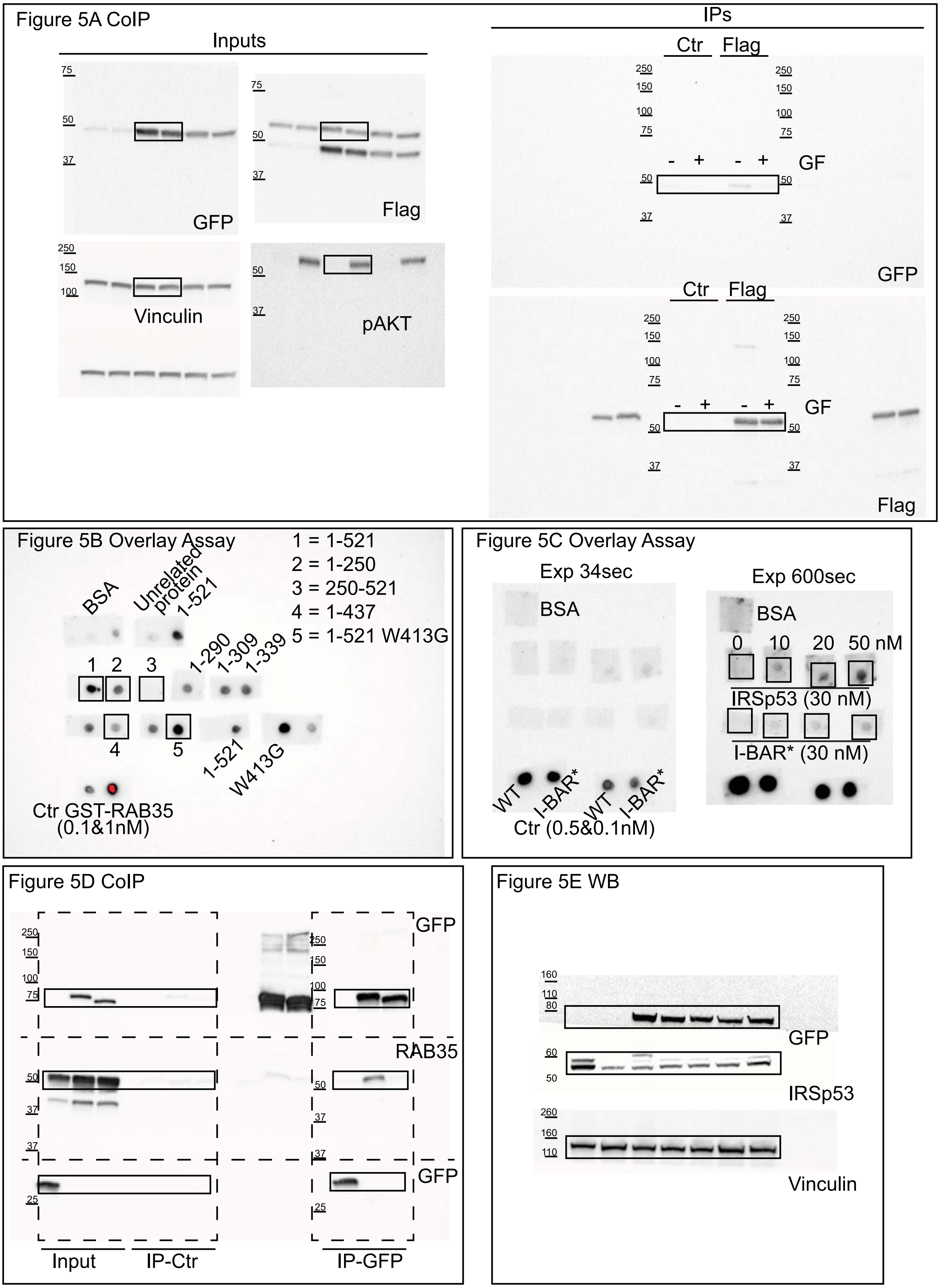
Figure 5A CoIP
Inputs
IPs
Ctr Flag
- + - + GF
GFP
Flag
Vinculin
pAKT
Figure 5B Overlay Assay
1 = 1-521
2 = 1-250
3 = 250-521
4 = 1-437
5 = 1-521 W413G
BSA
Unrelated protein
1-521
1-290
1-309
1-339
1 2 3
4
5
1-521
W413G
Ctr GST-RAB35
(0.1&1nM)
Figure 5C Overlay Assay
Exp 34sec
Exp 600sec
BSA
0 10 20 50 nM
IRSp53 (30 nM)
I-BAR* (30 nM)
WT
I-BAR*
Ctr (0.5&0.1nM)
Figure 5D CoIP
GFP
RAB35
Input
IP-Ctr
IP-GFP
Figure 5E WB
GFP
IRSp53
Vinculin

Figure S1 A WB

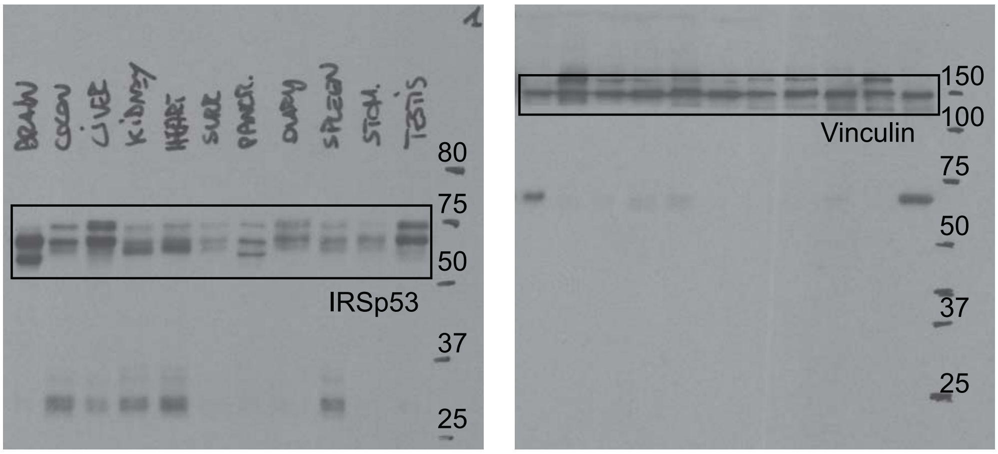


Figure S1 B

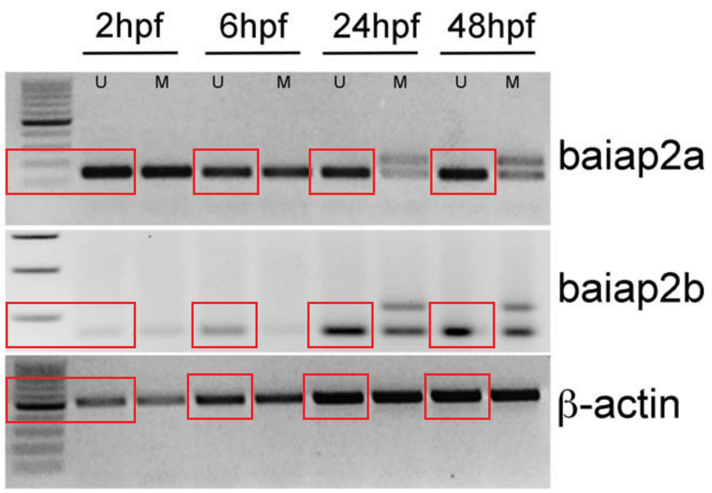


Figure S1 C Agarose Gels

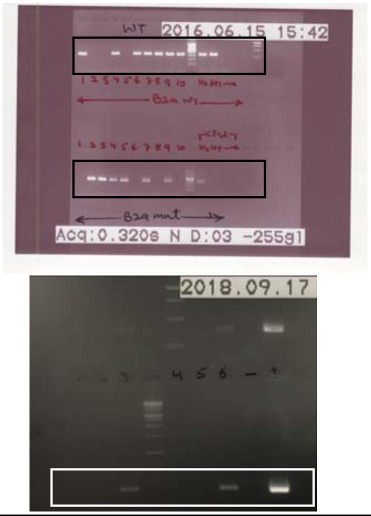


Figure S1 D WB

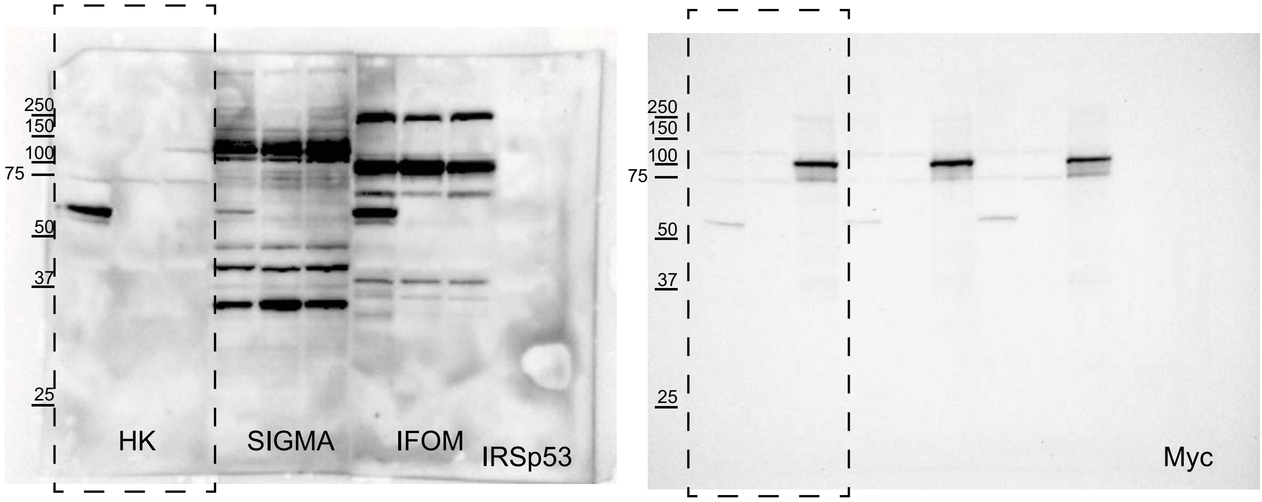

Figure S3 E WB

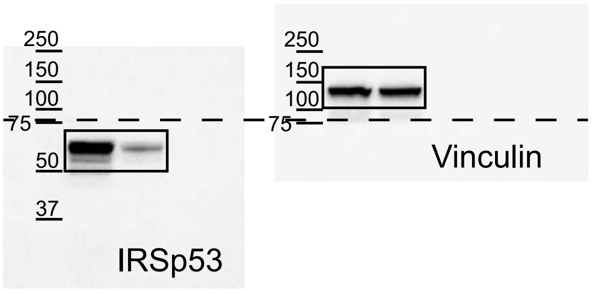

Figure S4 D Overlay assay

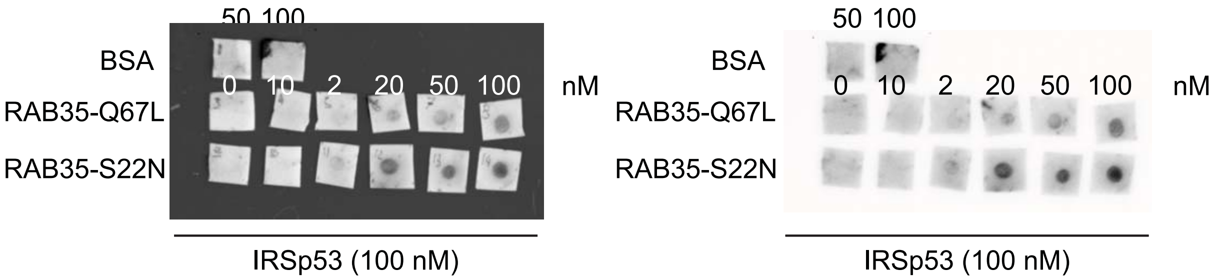

Figure S5 A WB

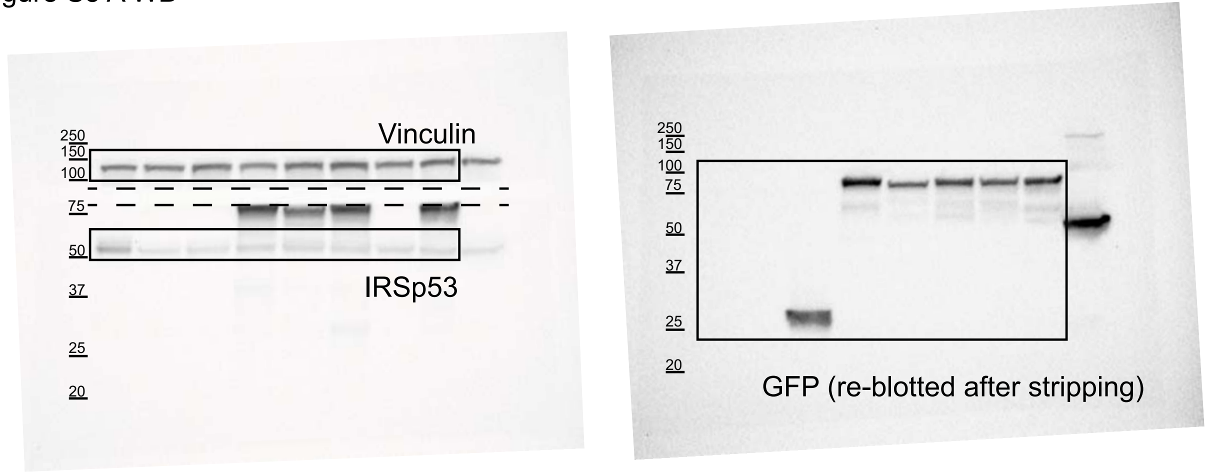

Figure S6 C WB

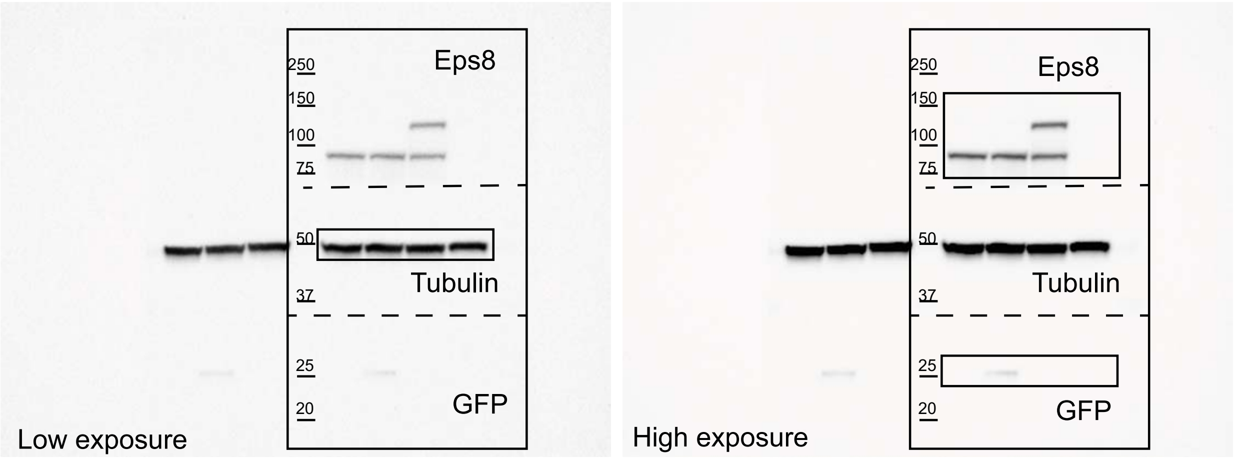


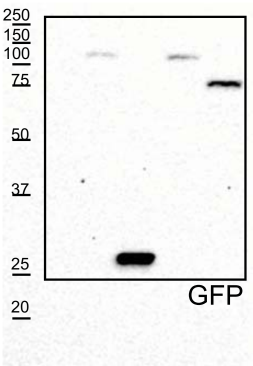


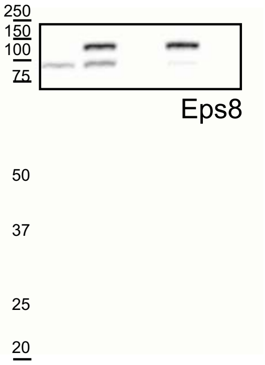
